# Bayesian analysis dissects kinetic modulation during non-stationary gene expression

**DOI:** 10.1101/2023.06.20.545522

**Authors:** Christian Wildner, Gunjan D. Mehta, David A. Ball, Tatiana S. Karpova, Heinz Koeppl

## Abstract

Labelling of nascent stem loops with fluorescent proteins has fostered the visualization of transcription in living cells. Quantitative analysis of recorded fluorescence traces can shed light on kinetic transcription parameters and regulatory mechanisms. However, existing methods typically focus on steady state dynamics. Here, we combine a stochastic process transcription model with a hierarchical Bayesian method to infer global as well locally shared parameters for groups of cells and recover unobserved quantities such as initiation times and polymerase loading of the gene. We apply our approach to the cyclic response of the yeast *CUP1* locus to heavy metal stress. Within the previously described slow cycle of transcriptional activity on the scale of minutes, we discover fast time-modulated bursting on the scale of seconds. Model comparison suggests that slow oscillations of transcriptional output are regulated by the amplitude of the bursts. Several polymerases may initiate during a burst.

## Introduction

Transcription is one of the fundamental processes of cellular life. RNA synthesis consists of the three major steps of initiation, elongation and termination. A closer look reveals that the individual steps are highly regulated and subject to intrinsic as well as extrinsic stochastic effects^1–4^. The details of transcriptional regulation on the molecular level are still far from understood.

Single-cell measurements of RNA have revealed that transcription is not only heterogenous between cells or within the genome but can also change for the same gene over time. The frequently observed pattern of high transcriptional activity interspersed with periods of silence is known as transcriptional bursting^5^. Recent evidence suggest that bursting may occur on multiple superimposed timescales^6^. It is unclear how these different timescales are regulated. A common question of interest is whether transcriptional output is regulated by burst amplitude, burst frequency, or burst duration^7–9^. Traditionally, this is investigated using RNA counting data. Continuous transcription with exponentially distributed time intervals between initiation events leads to a Poisson distribution of mature mRNA. Therefore, deviations from the Poisson distribution may indicate bursty transcription. Early work in this direction modeled the promoter as a telegraph process that stochastically switches between transcriptionally active and inactive states. In the active state, transcriptional output follows a Poisson process. For inference, predicted distributions of the model are matched to empirical RNA count histograms^5, 10, 11^. Later, the method was extended to multi-state models and to include nascent mRNA by solving the chemical master equation of the underlying system numerically^2, 9^. While successful at confirming bursting, information theory suggests that RNA counting data is fundamentally limited in distinguishing different multi-state promoters^12, 13^. In addition, the employed models oversimplify elongation and termination. Therefore, all variance in the observed data is necessarily attributed to the initiation process which may bias results towards more complex promoter models.

Novel imaging techniques such as the stem loop approach now enable time-resolved measurements of single transcription sites (TS) in live cells and in real time by fusing a fluorescent marker to a binding protein that attaches to hairpin structures formed by the nascent mRNA^14, 15^. Observed by a fluorescence microscope, the TS appears as a moving diffraction-limited spot with fluctuating intensity. An idealized time trace of a single polymerase consists of an approximately uniform increase in intensity, followed by a plateau phase when all stem loops have been formed and a sharp drop when the transcript detaches from the transcript site^16^. For actively transcribed genes, the observed intensity is a superposition of several polymerases that have initiated with varying interevent times. Together with other sources of noise accumulated during image acquisition the resulting trace may seem highly random. Thus, while time-resolved measurements of single transcriptions sites are potentially more informative than RNA counting data, they are also much more challenging from an inference perspective. Phenomenologically, bursting has been studied by binarizing the fluorescent traces via thresholding. Modifying suspected regulators then allows to measure the corresponding effect on burst statistics^17^. However, for fast switching dynamics, extracting bursts directly from the trace is prone to error^18^. First, a simple Poisson signal convolved with non-Gaussian observation noise and measured with a detection threshold may give an impression of burstiness. Second, if time between two bursts is shorter than the production time of the nascent mRNA, a bursty signal may be classified as continuous. A more sophisticated method for trace analysis that allows to determine initiation rate and expected production time of nascent transcripts is based on the autocorrelation function of the intensity signal^16, 19^. From a stochastic kinetic model of transcription, a theoretical autocorrelation function for the system is computed and then fit to match the empirical autocorrelation function of the traces. This provides a computationally efficient approach to extract average mRNA production times. An adaptation for dual-color labelling of the same transcript is also available^20^. However, by design, fluctuation analysis works best for stationary systems and long observation times. Extensions to non-stationary settings or more complex transcription models involving multi-state promoters or interactions between individual polymerases are challenging and currently rely on phenomenological corrections^20, 21^. An alternative idea is to split the problem into two parts. In a first step, initiation times are reconstructed from a fluorescent trace by a deconvolution algorithm. In a second step, the recovered initiation time distribution is then compared to theoretical predictions of different promoter models^22^. While this form of analysis proved effective, it requires reliable extraction of initiation time sequences which may not be possible for more irregular signals. Bayesian inference in combination with stochastic process models of transcription provides a principled framework to extract information from individual single-site traces. Due to the high computational demands, studies in this direction have been limited to simplified models where elongation and termination are treated as deterministic processes^23^. More complex models with stochastic elongation and termination have so far only been used within moment-based or simulation-based inference frameworks^24, 25^.

In this work, we use a kinetic model with stochastic treatment of the main transcription steps and develop a hierarchical Bayesian framework that performs joint inference on a collection of traces. The hierarchical approach allows one to jointly infer cycle-independent parameters shared by all cells and cycle-dependent parameters shared by cells within the same time window. This improves accuracy significantly compared to inferring data from individual traces and then pooling the results. We use this approach to investigate dynamic changes in the kinetics of transcription for *CUP1* promoter in *Saccharomyces cerevisiae*. Previously, *CUP1* has been shown to undergo a slow cycle of transcriptional activity with variable transcriptional output on a timescale of minutes in response to a heavy metal stressor^26^. Within this slow cycle, fast bursts of transcription on the scale of seconds regulated by fast interdependent cycling of transcription activator and chromatin remodeler were inferred from smFISH modeling^27^. In this work, we investigated the first period of the slow cycle by monitoring transcription sites in live cell using the stem loop approach and Bayesian inference. To account for the non-stationary setting, we split the cycle into short windows with higher frame rate and used the hierarchical Bayesian inference framework. By employing stochastic variational inference^28^, the method can handle datasets consisting of several thousand traces. Model comparison of several candidate models reveals fast bursting of *CUP1* on the scale of seconds, indicated by quasiperiodic transcription in individual cells through the slow cycle of bursting. This bursting on a faster timescale on the order of seconds is comparable in timescale with previously observed cycling of transcription activator on *CUP1* promoters^27^. Our discovery supports the hypothesis of fast bursts of transcription activated by fast cycling of TF. Regulation of the *CUP1* transcriptional output occurs most likely via modulation of the burst amplitude. In addition to parameter posterior distributions our method can recover via latent state inference based on stochastic filtering unobserved dynamic quantities such as initiation times and polymerase loading. We demonstrate that multiple polymerases may be loaded onto the same promoter during a burst. We also observe that the elongation speed of RNAP II varies, undergoing a slow cycle correlated with the slow cycle of transcription output. Our method reveals a delay between rise-and-fall patterns of observed bursts and the actual intervals of activity. While we demonstrated the method on non-stationary transcription data, our approach is applicable to a wide range of systems that can be modeled as a Markov jump process and can be straightforwardly adapted to other sources of heterogeneity. We provide a corresponding Python toolbox available at XXX.

## Results

### In continuous presence of Cu^2+^, *CUP1* undergoes bursts of transcription

*CUP1* encodes metallothionein protein Cup1 that protects the cells from heavy metal stress. *CUP1* is present in 10 tandem copies per chromosome VIII in *Saccharomyces cerevisiae*^26^. In yeast cells activated with Cu^2+^, quantification of mature mRNA by RT-qPCR reveals oscillations in *CUP1* mRNA transcriptional output in the cell population (Fig. 1a). This indicates that as described for several but not all systems, in continuous presence of an activator, *CUP1* transcription occurs not continuously but in oscillations: periods of transcriptional activity are interspersed with periods of transcriptional silence (Fig. 1h). Moreover, transcription output is modulated through the cycle. As shown previously, these oscillations are not dependent on cell cycle^26^. In living cells, we can monitor nascent mRNA formation at *CUP1* TS via the stem loop approach (Fig. 1b,c) ^14, 15^. A single ORF within the array of *CUP1* genes was replaced in one chromosome of the diploid yeast by a reporter encoding PP7 stem loops, visualized by the PP7 phage coat protein (PCP) tagged with GFP as a single green spot. Signal to noise was optimized in this system by low-level expression of PCP-GFP under a constitutive promoter *pSEC61* (see *SI Appendix*, Sec. S5.1). As the mRNA of the reporter contains only stem loops, it is not translated due to abundant stop codons, and thus, no protein is produced. Reporter is controlled by a natural *pCUP1* within the natural *CUP1* array; thus, expression of the reporter characterizes initiation from the *CUP1* promoters. However, the reporter sequence is different from natural *CUP1* ORF, and the transcript is longer. Thus, the production time for this reporter may not reflect the production of the wild-type *CUP1* ORF sequence.

**Figure 1.**
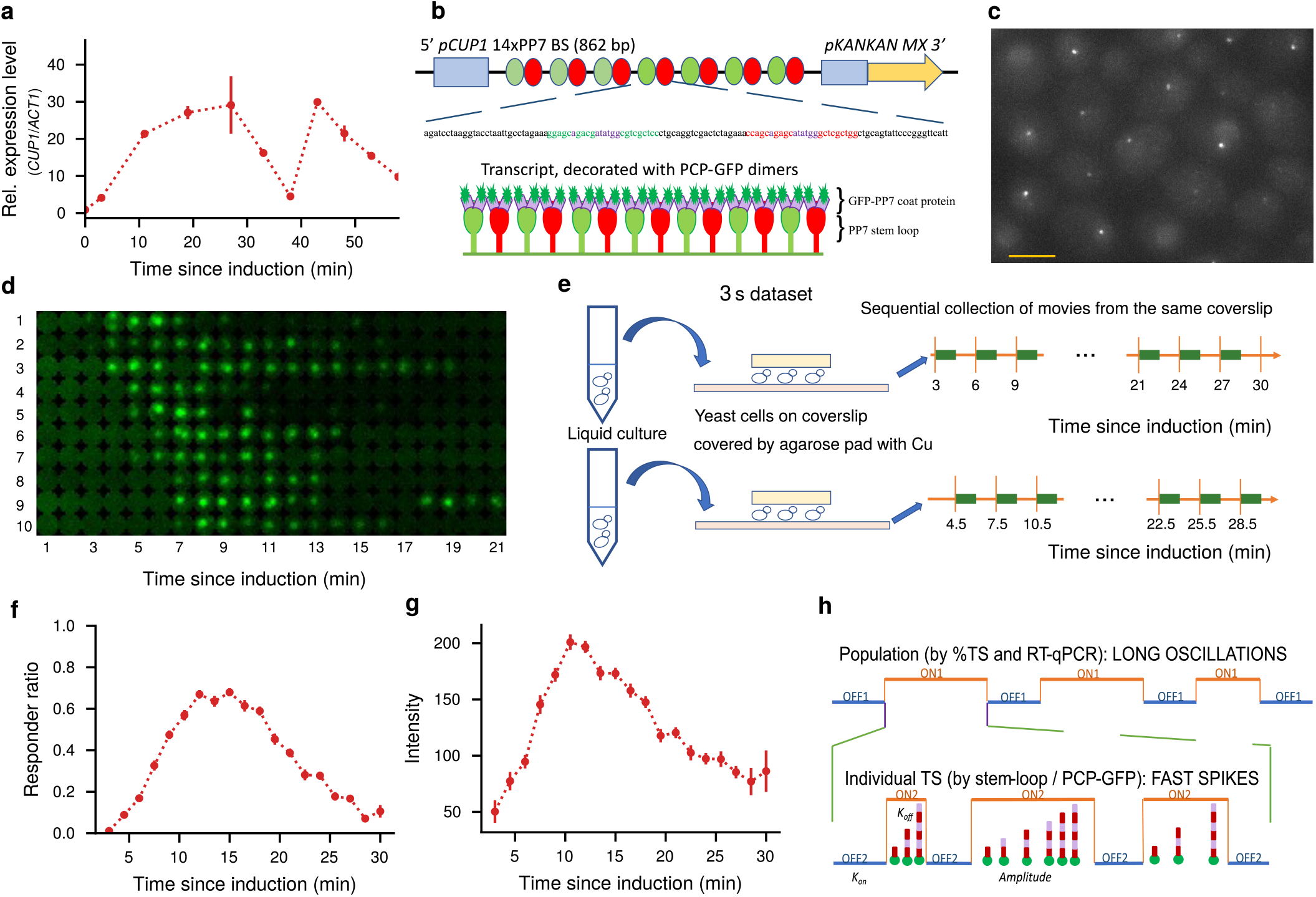
Engineering, visualization, and characterization of the *CUP1* transcription site. **a** Oscillations in *CUP1* mRNA expression level quantified by RT-qPCR and normalized by expression of the housekeeping gene *ACT1*. Error bars represent standard error of the mean (SEM) from two biological replicates. **b** Schematic of the 14x PP7 reporter replacing one copy of the *CUP1* ORF in chromosome VIII of *S. cerevisiae*. Two types of the stem loop sequence (red and green stems, purple loops, or bulges) are present in the reporter sequence, each stem is bound by a PP7-GFP dimer. Hence, 28 GFP molecules associate with single mRNA. The length of reporter transcript is 862 bp, however a few transcripts may be longer as there is no terminator immediately after 14xPP7. The expression of *KANMX* observed by RT-qPCR is constitutive and does not follow oscillations of 14X PP7 transcripts, which indicates that great majority of 14xPP7 transcripts stop before *KANMX* ORF (data not shown). **c** Example field view of cells with active TS containing nascent 14x PP7 reporter transcripts. Cells were imaged after 9 min of Cu^2+^ induction. Z-stack of the entire cell volume is presented as a maximum intensity projection for the GFP channel. Scale: 5 μm. **d** TS in individual cell display independent spikes of activity. TS dynamics from 10 representative cells are presented for the first 21 min since Cu^2+^ addition, imaged with 1 min time-lapse. Maximum intensity projections of the entire z-volume of the cells were cropped by keeping the TS in the center of the 13x13 pixels area. **e** Illustration of the movie collection for datasets imaged with 3 s time-lapse. Movies on the same coverslip are started every 3 min and are recorded for 90 s (green blocks). The remaining time is used to move the microscope to the next position on the coverslip. By collecting several such movie sequences starting either 3 min or 4.5 min after induction, the whole first cycle is covered. **f** The fraction of cells in the population showing an active TS follows the oscillation pattern of expression of the whole *CUP1* locus. Cells with TS were counted in independent fields imaged sequentially with 3 min time interval. Error bars represent standard error of the percentage (SEP). **g** TS in individual cells express more transcripts at the peak of the oscillation. In sequential 90 s movies collected after Cu^2+^ addition with 3 s time-lapse, the spot intensities were measured in the first frame of every movie. The graph shows the population average of these spot intensities indicating that transcriptional output of individual cells follows the oscillation pattern of the whole system. **h** Schematics of *CUP1* multi-scale bursting. Top – long oscillations between transcription (ON1 state) and no-transcription phases (OFF1 state) on the population level (cf. **a** and **d**). Bottom – short transcription spikes (ON2 and OFF2 states) as observed for individual TS (**f**). Transcriptional amplitude of the spikes is defined by the average number of mRNA produced per spike (striped lines represent different nascent mRNA produced during spike). The gene switches from an inactive OFF2 state to an active ON2 state with rate *k*_on_, and back to OFF2 state with rate *k*_off_. As this work focuses on the first ON1 phase, subscripts are dropped from hereon.

By counting the fraction of the cells with active nascent reporter mRNA over time (responder ratio), we confirmed a time-modulated response to constant activation by Cu^2+^ (first oscillation is quantified in Fig. 1f). Interestingly, in the movies of TS the average brightness of the individual TS also changed depending on the time of activation, indicating that at the peak of oscillation each TS produces more mRNA (Fig. 1g). This implies that the time-varying transcriptional output on the population level observed by RT-qPCR cannot be explained by a change in the number of responding cells alone but that parameters of transcription are modulated over time. Therefore, *CUP1* transcription is not in a steady state. Interestingly, individual TS observed through the first oscillation show independent dynamics, and the majority of the cells display several bursts of activity (Fig. 1d, exemplary cells no. 1, 3, 5, 7, 9). This suggest modulation on two scales: long-term oscillations on the order of several minutes governing transcriptional output of the cell population (slow cycle) and fast spikes (bursting) on the order of seconds regulating output of individual TS (Fig. 1h). The long-term oscillations are schematically represented by interspersed ON1 and OFF1 states. The short-term spikes are represented by interspersed ON2 and OFF2 states.

In this paper, we focused on the characterization of transcriptional activity within the first transcription phase of roughly 30 min. Due to photobleaching, such a long period cannot be imaged with our setup under a high frame rate. However, more importantly, the observed modulation of transcriptional output indicates that averaging through the long periods of observation may hide putative oscillatory variability in transcription parameters. We therefore split the cycle into small windows and performed parameter inference on all traces within the same window (cf. Fig. 4a). The first dataset was recorded with 12 s time-lapse for 300 s. We observed that a time-lapse of 12 s was too long for the relatively short gene template leading to problems separating elongation speed and termination rate. In addition, a duration of 5 min turned out too long to capture dynamic parameter changes adequately. Thus, we collected a second dataset with a duration of 90 s imaged with a timelapse of 3 s (Fig. 1e) leading to a high frame rate covering of the first 30 min after induction. After quality control, we extracted fluorescent traces with a custom tracker based on three-dimensional Gaussian fitting of the transcription site combined with ideas from stochastic filtering (*SI Appendix*, Sec. S12). Thus, we obtained two datasets of traces, each containing more than 3000 traces, with varying starting times during the first transcription phase. A detailed description of the pre-processing and the collected datasets is provided in *SI Appendix*, Sec. S6. To quantitatively analyze these non-stationary datasets we developed a Bayesian inference approach based on a stochastic kinetic model of transcription discussed in the following sections.

### Stochastic kinetic model of transcription

Our model (Fig. 2) is based on the totally asymmetric exclusion process (TASEP), where the DNA template is partitioned into *L* sites^30^. Polymerases initiate at site 1 with rate *k*_i_. After initiation, elongation proceeds in discrete steps with rate *k*_e_. Termination occurs at site *L* with rate *k*_t_ (Fig. 2a). Early termination is not permitted. While typical models of transcription assume independent polymerases, the TASEP model permits at most one polymerase per site allowing possible interactions in actively transcribed genes. Our model adopts this behavior for all but the termination site accounting for the possibility of transcripts residing at the TS after elongation. A typical progression of a single polymerase is shown in Fig. 2c.

**Figure 2.**
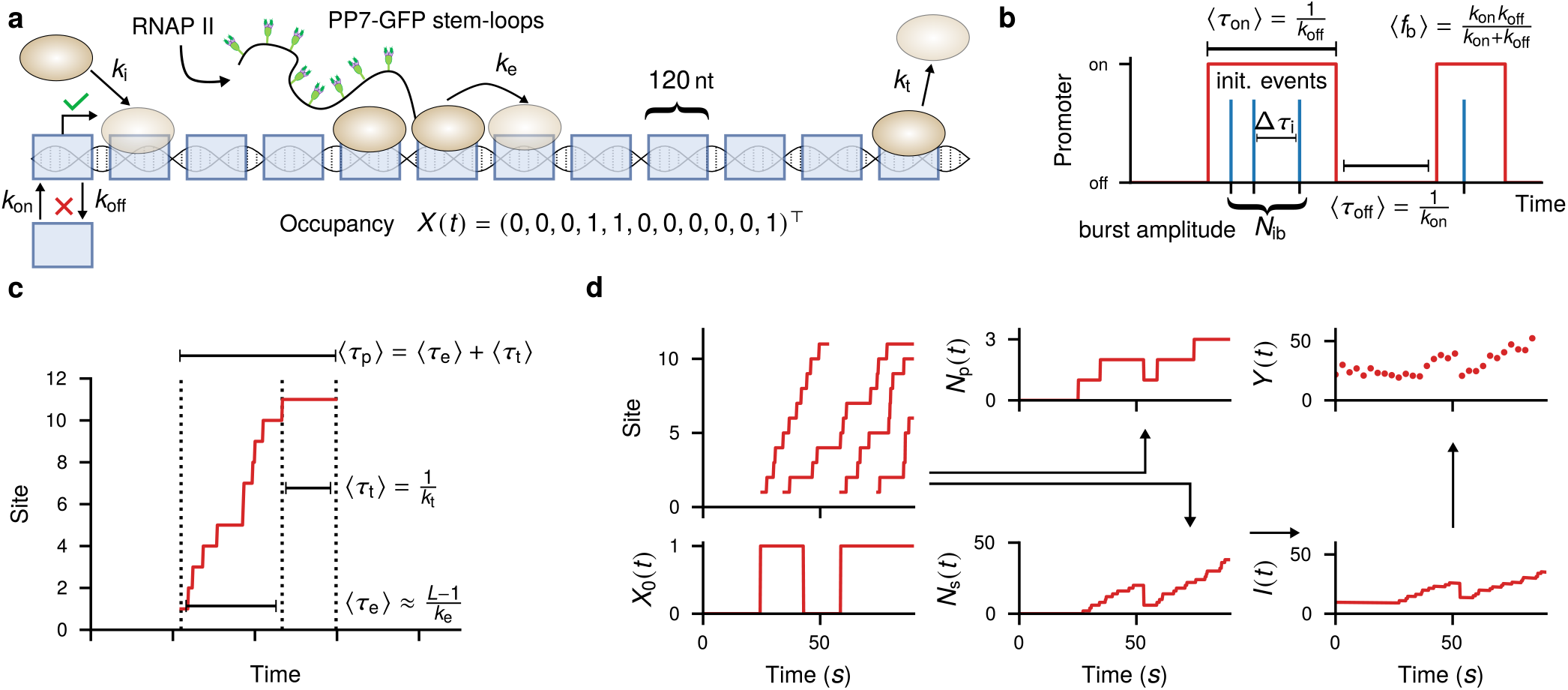
Stochastic kinetic model of transcription. **a** Kinetic transcription model based on TASEP. The DNA template is coarse-grained and partitioned into sites of 120 nt corresponding roughly to twice the footprint of a stem loop. Therefore, each of the first sites is associated with two dual GFP. The promoter switches between a transcriptionally active state (green) and an inactive state (red) with rates *k*_on_ and *k*_off_. In the active state, Polymerases initiate with rate *k*_i_, step along the lattice at rate *k*_e_ and terminate with rate *k*_t_. **b** Illustration of a two-state promoter model switching between an inactive and an active state with rates *k*_on_ and *k*_off_. These parameters implicitly define other bursting-related quantities such as the burst duration *τ*_on_, the time between bursts *τ*_off_ and the burst frequency *f*_b_. The burst amplitude *N*_b_ is defined as the number of initiation events per burst. **c** Illustration of a single polymerase initiating and progressing on the lattice. The three kinetic parameters of the TASEP model determine quantities such as termination time *τ*_*t*_, elongation time *τ*_*e*_ and mRNA production time *τ*_*p*_. The relation for expected elongation time is only valid when the polymerase density is low, in general there is now closed form expression available due to possible polymerase interactions. **d** Illustration of how to simulate synthetic traces. First, a trajectory from the telegraph-augmented TASEP model is created using the Gillespie algorithm. This typically produces a superposition of several polymerases. This is illustrated in form of a kymograph plot that shows the probability of a site to be occupied over time. From the full occupancy *X*(*t*) we can extract unobserved quantities of interest such as the polymerase loading *N*_p_(*t*), or initiation, elongation and production times of mRNAs (panel **c**) and the time between initiation events Δ*τ*_i_ (panel **b**). To simulate the measured fluorescence intensity, we first extract the number of stem loops *N*_s_(*t*) by summing the contributions of all polymerases. The spot intensity *I*(*t*) is then formed by multiplication with intensity per GFP *γ*, addition of background levels *b*_0_ and *b*_1_ and exponential bleaching with rate *λ*. Finally, the continuous-time intensity is sampled at equidistant time points and convolved with multiplicative noise to obtain the simulated measurements *Y* (*t*).

The times between transitions are exponentially distributed. Thus, the model is a continuous-time Markov chain (CTMC) describing the stochastic movement of RNAP II on the DNA template by the occupancy vector *X*(*t)* = (*X*_1_ (*t*), …, *X*_*L*_(*t*)). In order to model bursting, we introduce an additional promoter site *X*_0_(*t*) that switches between an active and an inactive state with rates *k*_on_ and *k*_off_ (Fig. 2b). In the extended model, initiation at site *X*_1_(*t*) is only allowed if the promoter site *X*_0_(*t*) is in the active state. This behavior is akin to the random telegraph model^31^ often used for the analysis of RNA counting data^9, 10, 17, 32^. Within this model, transcription dynamics are governed by the vector of parameters *θ* = (*k*_on_, *k*_off_, *k*_i_, *k*_e_, *k_t_*). The extended model is still a CTMC, therefore samples can be generated by the Gillespie algorithm. We denote such a full sample path as *X*_[0,*T*]_. The transient probability *p* (*x, t*) ≡ Pr (*X* (*t*) = *x*) satisfies a master equation

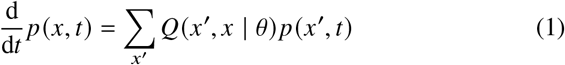

where the sum is over all possible configurations of the lattice and *Q* is the transition function of the process parametrized by *θ*. Details on how to construct *Q* are given in *SI Appendix*, Sec. S8.1.

In order to compare simulated to measured traces, the occupancy *X*(*t*) has to be converted to predicted intensity (Fig. 2d). As the positions of stem loops are known, one can compute the number of GFPs attached to every nascent mRNA from the lattice occupancy *X*(*t*). To convert the number of GFPs to predicted intensity, one requires a scaling factor *γ* and the bleaching rate *λ* and background variables *b*_0_, *b*_1_. The predicted intensity is then sampled at measurement times *t*_1_, …, *t*_*n*_ and passed through a multiplicative noise model to obtain a synthetic trace *Y* = (*Y*_1_, …, *Y*_*n*_). For later use, we combine all parameters related to the observation model into the vector *ω*. A detailed description of the observation model can be found in *SI Appendix*, Sec. S8.2.

### Calibration of the observation model

The parameters of the observation model have a major impact on the simulated measurements. Since this can cause issues with parameter identifiability, we perform independent calibration measurements for the scaling factor *γ* and the bleaching rate *λ*. To estimate the scaling factor, we engineered three yeast strains with sub-cellular structures attached to a known number of GFP molecules (see *Methods - Yeast strains and plasmids*) and measured spot intensities for a number of cells. As shown in Fig. 3a, the intensity distributions of the three constructs roughly follow a linear shape. Next, we combined the observation model for intensity prediction with vague priors and computed a posterior distribution using Monte Carlo. A similar Bayesian calibration was applied to time-lapse data of a static construct to determine the bleaching rate. An in-depth description is given in *SI Appendix*, Sec. S9. The posterior distributions from the calibration measurements where used as priors for the time trace inference.

**Figure 3.**
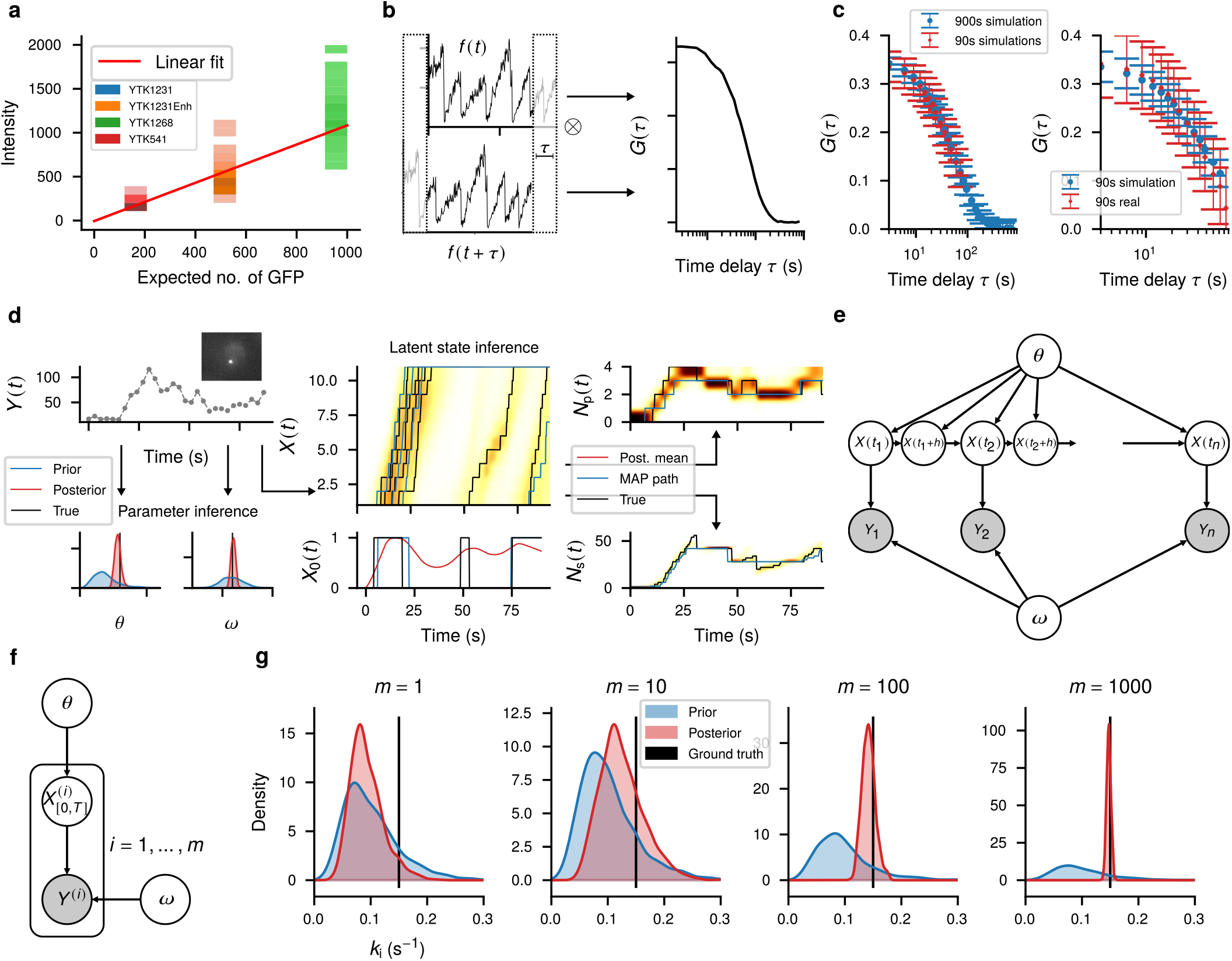
Inference from single cell traces. **a** Intensity distribution of the three strains used for calibration over the expected number of GFPs. Darker regions indicate a higher density. The red line indicates a linear regression fit to the data. **b** The autocorrelation of a single TS is generated by multiplying a signal shifted in time by a delay, *τ*, and multiplying it with the original signal and integrating. By repeating this process over many values of *τ*, the ACF function is generated for a single trace. Autocorrelation function averaged for many traces can reveal temporal characteristics of the system. **c** Left panel: Average autocorrelation of 1000 simulated trajectories for a time-lapse of 3 s observed over 90 s and 900 s showing an ideal autocorrelation function that could be analyzed to extract transcription parameters. Right panel: Average autocorrelation of CUP1 transcription sites from 282 cells imaged for 90 s with 3 s time-lapse after 9 min of Cu^2+^ activation compared to simulated results. **d** Illustration of Bayesian inference for single cell traces. From a measured trace *Y*(*t*), we obtain the posterior distributions of model parameters *θ*, and observation parameters *ω*. Note that this is an illustration as *θ* and *ω* are vectors of multiple parameters. In addition, latent state inference recovers the most likely sample paths of the unobserved lattice process *X*(*t*) and the promoter state *X*_0_(*t*). From these traces, other dynamic quantities of interest such as the polymerases loading *N*_*p*_(*t*) and the number of active stem loops *N*_*s*_(*t*) can be extracted. **e** Probabilistic graphical model representation of the single trace inference problem. Arrows indicate conditional relationships in the data generating process, grey color indicates that the corresponding node is observed^29^. The process *X*(*t*) is sampled at times *t*_1_, …, *t*_*n*_ and observed via noisy measurement *Y* = (*Y*_1_, …, *Y*_*n*_). As illustrated by the nodes *X* (*t*_*i* +_ *h)*, *X*(*t*) is continuous in time. **f** Graphical model of the joint inference problem with pooling of *m* traces to infer shared parameters. A plate indicates multiple conditionally independent variables given the parent^29^. Every pair 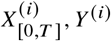 is of the form shown in **e. g** Gaussian kernel density representation of the results of joint Bayesian inference of the initiation rate on an increasing number of pooled simulated traces. As the number of traces increases, the posterior concentrates around the true value used to generate the data.

### Autocorrelation analysis is not applicable

A traditional approach to estimate kinetic parameters of live transcription sites is to use the autocorrelation function (ACF) of the intensity traces^16, 19, 21^ (Fig. 3b). However, ACF analysis is designed for long traces in a steady state setting. To test the applicability of this analysis to *CUP1* transcription, we analyzed traces for a single time window starting 9 min after the addition of Cu^2+^ (*Methods*) where TS activity is close to its peak (cf. Fig. 1g). Average ACF of real data, collected for 90 s total, is difficult to interpret and cannot be used for extracting kinetic parameters (Fig. 3c, right panel). As the ACF remains declining even at the longest delays, 90 s appear to be an insufficient amount of time for correlation analysis to be used. In fact, our simulations suggested that a TS has to be observed for at least 15 min (Fig. 3c, left panel). Although it may be possible to obtain an interpretable ACF by collecting over a longer time period, this would also lead to averaging over time-points from different parts of the slow cycle imposing the steady state assumption.

### Pooled Bayesian inference identifies model parameters

The goal of Bayesian inference is essentially to invert the data generation process depicted in Fig. 2. Given a measured trace *y* = (*y*_1_, …, *y*_*n*_) we want to reconstruct kinetic parameters *θ*, observation parameters *ω* (parameter inference). In addition, a stochastic process model also allows to reconstruct the most probable lattice configurations over time *x*_[0,*T*]_ given the data (state inference, Fig. 3d). More formally, this corresponds to computing the joint posterior distribution *p* (*θ, ω, x* _[0,*T*]_ | *y*_1_, …, *y*_*n*_). A probabilistic graphical model (PGM) representation of the single trace inference problem is given in Fig. 3e.

Sampling from this posterior distribution using Markov chain Monte Carlo (MCMC) involves evaluating the marginal data likelihood *p*(*y*_1_, …, *y*_*n*_ | *θ, ω*) which in turn requires integration of the master equation (1) for many different configurations of model parameters *θ* and observation parameters *ω*. We developed an efficient approach to evaluate the marginal likelihood and its gradient in parallel for multiple traces which allows to apply efficient gradient-based inference algorithms such as Hamiltonian Monte Carlo (HMC) and stochastic variational inference (SVI) (*SI Appendix*, Sec. S10).

Inference from single traces is often challenging due to issues with parameter identifiability^33^. To test identifiability for our setup, we simulated a set of synthetic traces following the steps illustrated in Fig. 2 using a fixed parameter configuration (*SI Appendix*, Table S14). Indeed, Bayesian inference of a single trace essentially reproduces the prior distribution indicating that a single trace does not contain sufficient information to identify the system (Fig. 3g, left panel). Pooling multiple traces and performing inference jointly (Fig. 3f) can improve the results substantially. Indeed, inference accuracy increases with the number of pooled cell, implying the system is identifiable given sufficient data (Fig. 3g, right panel). We stress that the pooling performed in joint Bayesian inference is a principled approach and more reliable compared to performing inference on single traces and then comparing the posterior means. An extended plot showing the posterior of more parameters is provided in *SI Appendix*, Fig. S19.

### Hierarchical Bayesian model captures slow cycle of transcription

In order to analyze the dependence of the parameters on the slow cycle, we split the dataset into subgroups pooling all traces that share the same time window since induction. As individual movies are short (90 s), we can assume constant parameters during individual windows leading to a joint Bayesian inference problem (cf. Fig. 3f,g) for each window. This straightforward approach has two problems. First, in some of the windows the number of traces is quite small (*SI Appendix*, Table S13) leading to unreliable inference. Second, not all of the parameters are expected to depend on the slow cycle. To take full account of the pooling, we developed a collection of mixed hierarchical models where some parameters are shared locally between traces in the same window and others are shared globally between all traces (Fig. 4a). A PGM representation of one such model with local initiation rate is shown in Fig 4b. For each of these models, we also included a version with a constitutive promoter (i.e. *X*_0_(*t*) = 1 for all times) to investigate if the data supports the bursting hypothesis. While increasing inference accuracy, a hierarchical model of 3000 traces was too computationally expensive for MCMC, as every step of MCMC requires evaluation of the log-likelihood of all traces in the dataset. Inference of the full dataset was therefore done by SVI^28^.

Allowing local variability of different parameter combinations gives rise to a collection of models. Bayesian model selection based on the marginal likelihood provides a systematic approach of finding the most likely model given the data and automatically penalizes models with too many free parameters^34^. As the marginal likelihood is costly to compute, we used the evidence lower bound (ELBO), that is obtained by variational inference, as an approximation. To prevent possible issues with model mismatch that cannot be detected from the marginal likelihood alone, we designed an additional metric based on the posterior predictive^35^ that relies on the Wasserstein distance between predicted and measured cycle (Fig. 4c and *Methods — Model selection*).The results of the model selection are shown in Fig 4d. A small value of the Wasserstein distance indicates that that data simulated from the learned model (posterior predictive) agrees well with measured data. In contrast, a higher value of ΔELBO suggest that the learned model is more probable than other models, given the data. Consequently, the overall best models are found in the lower right region of the graph. We also observe that most of the investigated models are close to a line in the two-dimensional evaluation space, indicating consistency of the two scores. A full account of all tested models on both datasets is given in *SI Appendix*, Table S15. We observe that a time-dependent elongation rate alone cannot explain the cycle. Additionally, the graph suggests that a switching promoter is more probable than a constitutive promoter. The best explanation of the slow cycle is a combination of time-dependent initiation rate with time-independent promoter switching rates. This implies that for *CUP1* the cyclic response is likely regulated by burst amplitude rather than burst frequency. The best ranking models are able to reproduce the intensity pattern over the cycle fairly well (Fig. 4e). Corresponding parameter posteriors of the best ranking model are shown in Fig. 4f,g. The posterior initiation rate (Fig. 4f) closely follows the cycle pattern known from responder ratio and spot intensity distribution (cf. Fig. 1f, g). Interestingly, the variance of the posterior is larger close to the cycle peak.

Previously we demonstrated that heterogeneity in *CUP1* transcriptional response is exacerbated by depletion of chromatin remodeler RSC^27^. This implies that this heterogeneity is caused by variable accessibility of the binding sites in the promoter to transcription activator Ace1p. Thus, we propose gradual changes in promoter accessibility through the slow cycle of transcription. Current observation is compatible with the following hypothesis - the rate of increase in accessibility is the same for all cells, but the maximal opening of the binding sites at the peak of the cycle may vary, providing heterogeneity in transcriptional output.

Posterior distributions for global model parameters are given in Fig. 4f. The full graph including all observation parameters can be found in *SI Appendix*, Fig. S20. As distributions of *k*_on_ and *k*_off_ are close, the promoter seems to be active approximately half of the time with an active time of ≈25 s.

**Figure 4.**
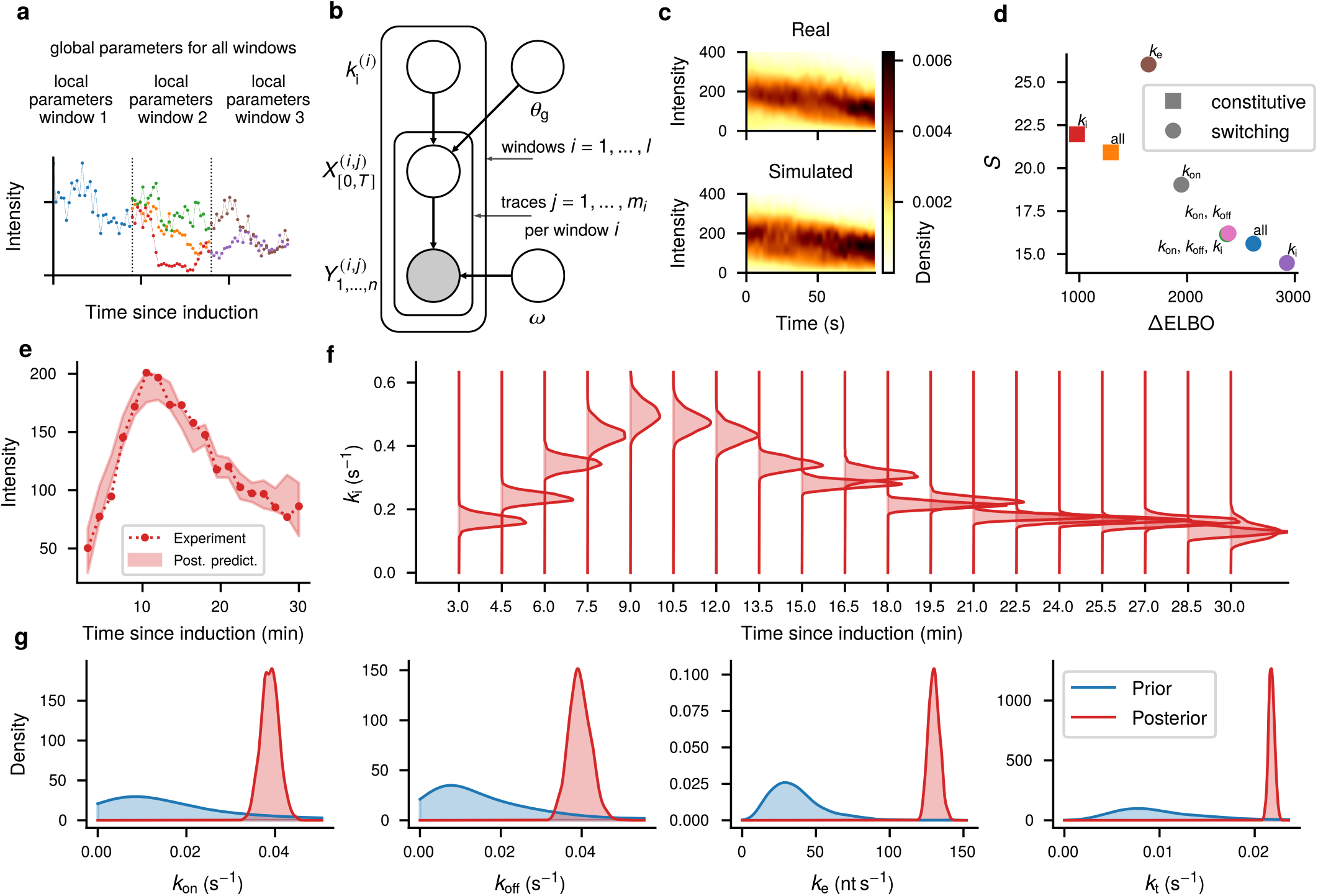
Hierarchical Bayesian inference of the first slow cycle. **a** Illustration of the hierarchical model for the first slow cycle. Traces are grouped into windows according to the starting time with respect to Cu^2+^ induction. Parameters are split into local and global parameters. Local parameters are shared between traces of the same window, global parameters are shared between all traces. While for illustration purposes, only three windows are shown, the traces of the 3 s dataset are assigned to 19 windows. **b** A PGM representation of the hierarchical inference problem for a model with local initiation rate and all remaining parameters constant. **c** Illustration of the predictive score *S*. Synthetic traces are simulated from the fitted model. The intensity distribution of the simulated data is compared to the intensity distribution of the real data by the Wasserstein metric, a general similarity measure for probability distribution. The score *C* is obtained by averaging over all windows and the parameter posterior. **d** Two-fold model selection based on the posterior predictive score *S* based on the Wasserstein distance and approximate Bayes factor ΔELBO. For *S* smaller values indicate that simulated data is more similar to measured data (smaller is better), for ΔELBO higher values indicate a larger probability of the model compared to a reference model. The reference model, assuming no switching and only global parameters, is the same for all data points. **e** Mean intensity of the first frame of traces over the slow cycle. The dashed line indicates experimental results of the 3 s dataset (cf. Fig 1), the shaded regions corresponds to a 90 % credible interval of the posterior predictive distribution. **f, g** Gaussian kernel density representations of the SVI approximate parameter posteriors of the best ranking model. **f** Local initiation rate per time-window since induction. **g** Global kinetic parameters.

### Stochastic filtering reveals time dependence of unobserved quantities

The true power of a continuous-time stochastic process is the possibility for latent state inference (cf. Fig. 3d). Given a measured trace, we can reconstruct the trajectories of the latent stochastic process that have most likely produced the observations by the backward filtering forward sampling approach (*Methods - Bayesian inference*). Results for three exemplary traces are shown in Fig. 5a. These traces are selected to reflect typical cases present in the dataset: continuous expression, a single burst and a double burst. The posterior predictive plots in the top panel demonstrate that our model is capable of explaining qualitatively different traces of the dataset. The reconstructed distributions of the number of polymerase and stem loops over time closely follows the shape of the observations (Fig. 5a, second and third row). The polymerase distribution indicates that *CUP1* is a highly transcribed gene that binds multiple polymerases simultaneously. This observation agrees with earlier predictions from smFISH modeling^27^. From the kymograph plot (fourth row), we can observe the movements of individual polymerases along the lattice. The bottom row demonstrates that the model is capable of reconstructing the promoter activity. Importantly, the active intervals are shifted in time with respect to the rise-and-fall patterns of the measurements and the inactive phase is identified even though the intensity does not drop to base level. This is an advantage of a model-based method compared to approaches that identify bursts directly from the traces. As shown in the kymograph, several polymerases may initiate during a burst. From the posterior path distribution, arbitrary path statistics such as the number of initiation events in a given time interval or the distribution of time between initiation events can be computed. By comparing such statistics of posterior paths over the cycle, we can investigate the time dependence of unobserved quantities. We stress that this is different from analyzing sample paths simulated from the fitted model. As the model is a Markov process, forward simulations will always exhibit exponential inter-event time distributions between fundamental events. In contrast the posterior process is non-homogenous, and can recover non-exponential inter-event times if the data provides evidence accordingly. As one path statistic of interest we investigated local elongation times by which we mean the average dwell time of the polymerases on individual sites as it progresses the lattice (cf. Fig. 2c). A graphical representation of the elongation times over the lattice and over the cycle is shown in Fig. 5b. Interestingly, polymerases tend to progress slower during the peak of the cycle. As during the peak of the cycle, the number of transcribing polymerases is also higher, this indicates a higher polymerase density is associated with lower average speed per polymerase. This could be caused, e.g. by steric hinderance from tightly spaced polymerase or competition for elongation factors.

**Figure 5.**
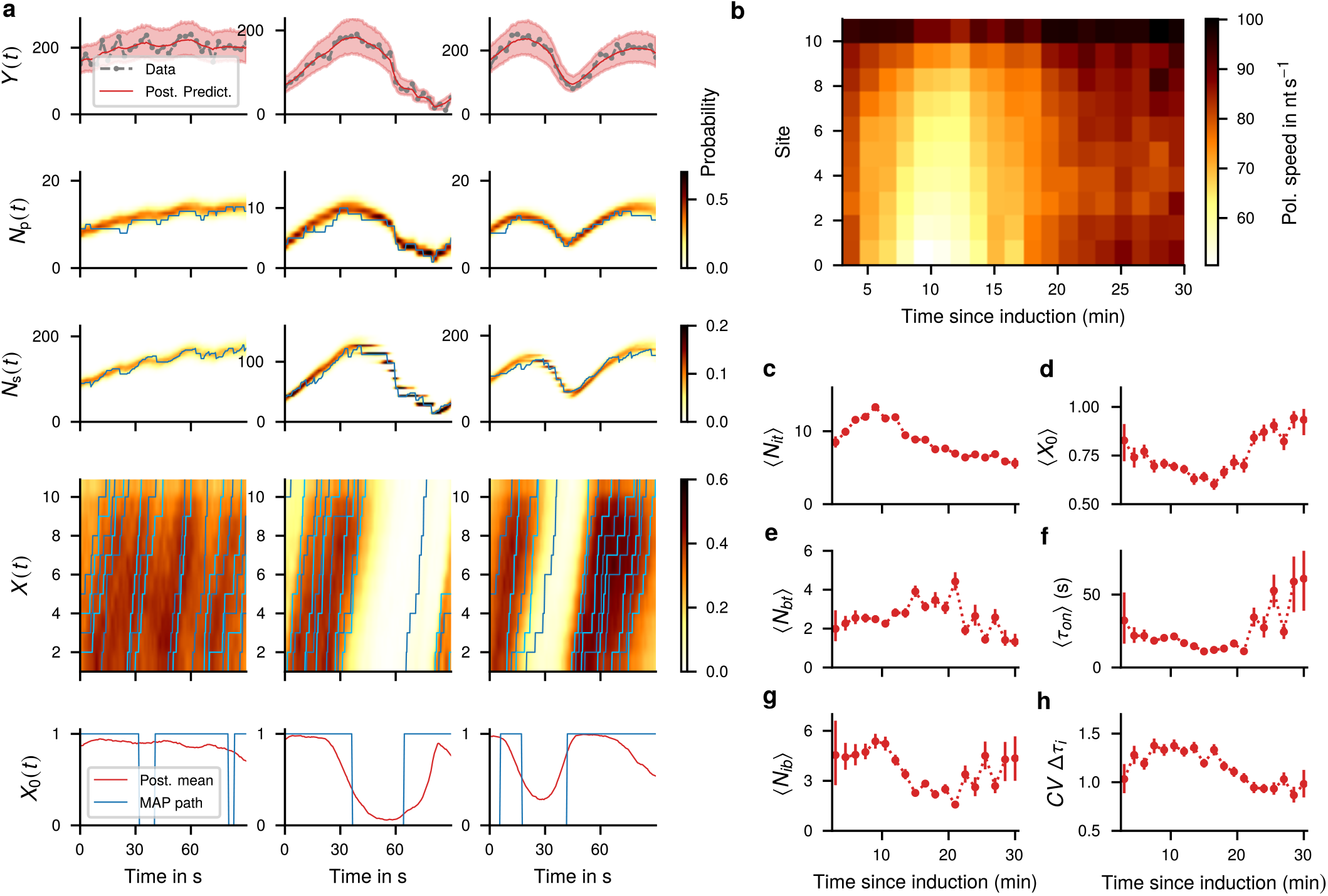
State inference. **a** Posterior path distributions for three characteristic traces from the 3 s dataset. The top panel shows the measured data *Y*(*t*) (gray dots) compared to the corresponding posterior predictive mean (red line) with 90 % credible interval (red shaded). Lower panels show the posterior distribution over time of the polymerase loading *N*_*p*_(*t*) including termination site, the number of active stem loops *N*_*s*_(*t*), kymograph of actively elongating polymerases and promoter activity *X*_0_(*t*). Here, darker colors indicate higher probability. The three selected traces represent typical cases in the dataset: a spot that is already active when imaging starts and stays active during the interval (a), a single rise followed by a decay (b), and a more dynamic site with multiple separated phases of activity. **b** Average local elongation speeds per site and over the slow cycle. Elongation speeds were computed by simulating traces from the state posterior as shown in panel **a**. For these traces, we extracted the times polymerases remained at each individual sites. Result were collected for all traces of a window and averaged over the posterior paths. **c-h** Initiation-related summary statistics over the first slow cycle. Statistics were computed by simulating a posterior sample (one smoothing sample for every trace in the dataset) and then pooling over windows. The results were then averaged over the parameter posterior. Red dots indicate the posterior mean, error bars correspond to 90 % credible interval. **c** Average number of initiation events per trace. **d** Average fraction of time spent in the active promoter state. **e** Average number of bursts per trace. **f** Average duration of a burst. **g** Average number of initiation events per burst (burst amplitude). **h** Coefficient of variation of the distribution of times between initiation events.

We investigated a number of selected path statistics related to initiation dynamics (Fig. 5c-h). The number of initiation events per trace closely follows the cycle. This is expected as the number of initiation events is directly related to the initiation rate *k*_i_ (Fig. 5c). In contrast, the mean activity of the promoter shows an inverse relation which seems counter intuitive (Fig. 5d). A possible explanation is the small number of initiation events away from the peak. When there are few events, it is not possible to reliably distinguish bursty and non-bursty behavior. Therefore, inference favors the simpler explanation of constitutive expression with small rate. An alternative explanation would be a multi-state promoter that is in a leaky baseline state away from the peak and switches to a more active regime during the cycle peak. The average burst duration (Fig. 5f) points in a similar direction: In the later part of the cycle, burst durations are longer with smaller number of initiation events per burst. The number of bursts per trace does not show a clear dependence on time since induction (Fig. 5e). The burst amplitude follows the cycle in a slightly shifted form (Fig. 5g). Interestingly, the peak of the cycle shows shorter burst times combined with higher burst amplitude meaning that time between initiation events is much shorter during the cycle peak. This suggest that efficiency of the initiation machinery is one target of regulation during the slow cycle. Finally, we studied the distribution of times between initiation events (Fig. 5g) by means of the coefficient of variation (CV). A CV of one corresponds to an exponential distribution and suggest constitutive expression. A CV larger than one indicates a heavy-tailed distribution which suggest burstiness. Indeed, in the beginning and in the end, the coefficient of variation of the inter-event time distribution is close to one suggesting exponential behavior while close to the peak we observe larger values indicating burstiness. Note that for some quantities the uncertainty is significantly larger away from the peak. This is explained by the smaller number of selected traces in the corresponding windows (cf. *SI Appendix*, Table S13).

## Methods

### Yeast strains and plasmids

We utilized haploid strains of *Saccharomyces cerevisiae* (BY4742 and BY4741) for live transcript analysis. The strains were engineered to include 14X PP7 binding sites and a MET3 integrative vector for expressing PP7-NLS-GFP. For photobleaching correction and GFP calibration, we used strains YTK541, YTK1231, and YTK1268, each containing a known number of GFP molecules per locus. Further details are provided in the *SI Appendix*, Sec. S5.1.

### Media and growth conditions

YTK1799 cells were grown in CSM-URA media under specific conditions for live transcript analysis (*SI Appendix*, Sec. S5.2). The growth protocol involved a series of inoculations, refrigeration, and daily inoculations to maintain consistent results. Cells were harvested and placed under a CSM-URA agarose pad for imaging. Strains YTK541, YTK1231, and YTK1268 were grown under similar conditions for photobleaching correction and GFP calibration.

### Quantitative RT-PCR (RT-qPCR)

RNA was extracted from samples at specified time points post-Cu induction. The extracted RNA was used to prepare cDNA, which was then used for quantitative real-time PCR (qPCR). The expression of the housekeeping gene ACT1 was used for normalization. The process was repeated at least twice, with qPCR performed in duplicates for each experiment.

### Microscope settings and imaging conditions

Live cells were imaged using a DeltaVIsion Elite Microscope under specific conditions (*SI Appendix*, Sec. S5.4). 3D time-lapse movies were acquired at room temperature using a specific imaging regime to cover the entire slow cycle. The same conditions were used for imaging strain YTK1231 for photobleaching correction.

### Trace extraction

We developed a custom 3D method based on sequential filtering to track fluorescence spots and quantify fluorescence levels. The method involves a state estimation using an approximate recursive filter based on the Laplace approximation and includes a binary state to take account of vanishing spots. Details are provided in *SI Appendix*, Sec. S6.2 and Sec. S12.

### Calibration measurements

We performed calibration measurements using strains YTK541, YTK1231, and YTK1268 containing a known number of GFP per locus. Spot intensities were measured as for the live cell experiments but only for a single time point. Assuming a linear relationship between number of GFP and intensity, we extracted estimates of the scaling factor by Bayesian log-linear regression. Similarly, an independent estimate for the bleaching rate was obtained by recording videos of strain YTK1231 and assuming an exponential decay of spot brightness. A detailed description is given in *SI Appendix*, Sec. S9.

### Stochastic modeling

The model is an instance of a Markov jump process that satisfies the master equation 1. The system’s state represents the occupation of lattice sites by RNAP II and the transition matrix is defined by initiation, elongation and termination along with the exclusion principle (*SI Appendix*, Sec. S8.1). By splitting the transition matrix into contributions corresponding to individual parameters, the master equation can be solved efficiently for a given parameter vector for fairly large state spaces by the Krylov subspace approximation for matrix exponentials^41, 42^. The lattice state is converted to fluorescence intensity by assuming an affine-linear dependence on the number of formed stem loops and a multiplicative noise model with a correction for small intensities. Details are provided in *SI Appendix*, Sec. S8.2.

### Autocorelation analysis

The average autocorrelation functions (ACF) were calculated as described in^21^ using the intensities of all TS tracks over time. The ACF was calculated for 3s intervals and 12s intervals after correcting the individual traces for photobleaching.

### Bayesian inference

The joint posterior *p* (*θ, ω, x*_[0,*T*]_ | *y*_1_, …, *y*_*n*_) of model parameters *θ*, observation parameters *ω* and latent lattice trajectory *x*_[0,*T*]_ was split into marginal parameter posterior *p* (*θ) p* (*ω) p* (*y* | *θ, ω*) and the conditional state posterior *p x*_[0,*T*]_ |*θ, ω, y*_1_, …, *y*_*n*_). Sampling from the marginal parameter posterior was realized by HMC and SVI using the probabilistic programming language Pyro^45^. To integrate the stochastic process model with Pyro, we developed a procedure to evaluate to evaluate the marginal data likelihood log *p* (*y* | *θ, ω*) by stochastic filtering. Combined with a modified backward filter to compute the gradients, we designed a differentiable inference procedure applicable to general Markov jump process models. To recover the full posterior, we used backward filtering forward sampling approach to generate posterior paths from the conditional state posterior *p* (*x*_[0,*T*]_ | *θ, ω, y*_1_, …, *y*_*n*_). For a more comprehensive description, we refer to *SI Appendix*, Sec. S10. The corresponding code is available as a Python package at XXX.^48^

### Model selection

We used a two-fold approach to Bayesian model selection. The evidence lower bound obtained from variational inference was used to approximate the marginal likelihood of different models and corresponding Bayes factors. In order to check for model mismatch, we used an additional metric based on the Wasserstein distance of the posterior predictive distribution and the empirical data distribution. The details are given in *SI Appendix*, Sec. S11.

## Acknowledgements

C.W. and H.K. acknowledge support by the European Research Council (ERC) within the CONSYN project, grant agreement number 773196. T.S.K. and G.D.M. were supported by Intramural Research Program of National Institutes of Health (NCI, CCR). The authors gratefully acknowledge the computing time provided to them on the high-performance computer Lichtenberg at the NHR Centers NHR4CES at TU Darmstadt. We would like to thank Dr.Tineke Lenstra for the plasmid pTL031, and Dr. Daniel Larson for the discussion of autocorrelation analysis.

## Author contributions statement

T.S.K. and G.D.M. conceived and conducted wet lab experiments. C.W. and H.K. designed model and inference procedures. C.W. developed inference software and performed Bayesian data analysis. D.A.B. performed fluctuation analysis. T.S.K. and C.W. wrote the initial manuscript. T.S.K. and H.K. supervised the project. All authors prepared, reviewed and edited the manuscript.

## Additional information

The authors declare no competing interests.

**SI Appendix**

### S1 Experimental details

#### S1.1 Yeast strains and plasmids

For live transcript analysis, we engineered the haploid strains of *Saccharomyces cerevisiae* (BY4742 and BY4741), which are isogenic to S288C (Research Genetics/Invitrogen, Huntsville, AL). 14X PP7 binding sites (hairpins) were amplified from pTL031^36^ using primers T1053, T1054 and integrated in one of the haploids by replacing one of the *CUP1* ORFs in the yeast genome by homologous recombination^37^. For expressing PP7-GFP coat protein, we constructed a MET3 integrative vector (pTSK630) to express PP7-NLS-GFP from *SEC61* promoter and integrated this vector by SacI XhoI digestion in both the haploids. This vector can be available upon request. For the photobleaching correction, we used a diploid strain YTK1231 in which both the *CUP1* arrays are replaced by 256 copies of *LacO* and lacI-GFP-NLS is expressed from *pHIS3*. For preparing the calibration curve for the number of GFP molecules verses brightness/intensity, we used three yeast strains (YTK541, YTK1231 and YTK1268) with known numbers of GFP molecules per locus. YTK541, contains a tandem array of 10 copies of *CUP1* locus with 40 binding sites for the transcription activator Ace1p-GFP. *CUP1* is activated by Cu, and at the peak of activity *CUP1* array binds 120 molecules of GFP. In YTK1231, each lacO binding site may be associated with a dimer of the lac Repressor (LacI-GFP)^38^. Therefore, the array of 256 tandem lacO binding sites is associated with 512 LacI-GFP molecules. In YTK1268, the spindle pole body of the diploid yeast strain contains approximately 1000 molecules of Spc42-GFP. Strain genotypes are provided in Table S8, plasmids in Table S9 and primer sequences in Table S10.

#### S1.2 Media and growth conditions

For live transcript analysis, cells of YTK1799 were plated on CSM-URA plate (from −80°C frozen glycerol stock) and grown for 48 h at 28 °C. 3 to 5 colonies were inoculated in 3 mL CSM-URA media (in 14 mL polypropylene tubes, Cat no. 352059, Falcon, Maxico) and grown for overnight at 28 °C, 230 RPM. 250 μL of this overnight grown culture was inoculated in 25 mL CSM-URA (in 250 mL flask) and grown at 28 °C, 230 RPM for 24 h. This flask was removed from the shaker and kept in refrigerator at 4 °C. We used this refrigerated culture for daily inoculations for a month to get consistent results (to avoid day to day variations in transcription induction kinetics due to difference in the age of the culture). From this refrigerated culture, we inoculated 60 μL in 3 mL of fresh CSM-URA media (in 14 mL polypropylene tubes, Cat no. 352059, Falcon, Maxico) in the morning and grew the cultures for 5 h at 28 °C, 230 RPM. Cells were harvested by centrifugation (2200 RPM for 1 min) and cells were placed under the CSM-URA agarose pad (100 μM CuSO_4_) for imaging. For the photobleaching correction and GFP calibration curve, strains YTK541, YTK1231 and YTK1268 were grown under the same conditions, except YTK541 and YTK1268 were grown in CSM-HIS media.

#### S1.3 Quantitative RT-PCR (RT-qPCR)

Samples were harvested at indicated time points after Cu induction. RNA was extracted (from yeast cells) using the ISOLATE II RNA Mini kit (Bioline, UK, Cat no. BIO-52072). cDNA was prepared using the iScript cDNA synthesis kit (BioRad, Cat no.: 1708891) starting with 1 mg of total RNA. Quantitative real-time PCR (qPCR) was performed as described^27^. For normalization, the expression of the housekeeping gene ACT1 was quantified. Primers used for this quantification are listed in Table S10 (T531, T532 for CUP1 and T1055, T1056 for ACT1). To confirm the absence of contaminating genomic DNA in cDNA preparations, reverse transcriptase negative (-RT) samples were used as a control, which produced the Ct value difference of >10 cycles between “-RT” and “+RT”samples, indicating a negligible amount of genomic DNA contamination in cDNA samples. mRNA extraction, cDNA synthesis, and qPCR were repeated at least twice, and qPCR was performed in duplicates for each experiment. Error bars indicate SEM.

#### S1.4 Microscope settings and imaging conditions

For imaging live cells, 5 h grown cultures were harvested by centrifugation (2200 RPM for 1 min) and 3 μL of cell pellet were placed in Lab-Tek II chambered coverglass (1.5 Borosilicate Gass, Nunc, ThermoFisher Scientific, MA, US), mixed with equal volume of 200 μM CuSO_4_ containing CSM-URA and covered by 1cm x 1cm CSM-URA agarose pad (100 μM CuSO_4_). A timer was started immediately upon mixing the cells with 200 μM CuSO_4_ containing CSM-URA. 3D time-lapse movies were acquired at the room temperature on the DeltaVIsion Elite Microscope, using 100x 1.4 NA oil immersion objective lens, sCMOS camera, FITC filter set (15 ms exposure, Ex 488/27; Em 505/45, Chroma Technology Corp, Bellows Falls, VT), 15 z-steps at every 400 nm, 1x1 binning and 1024x1024 pixels. Time-lapse movies with 3 s time interval were acquired for 90 s, followed by changing the field within next 90 s, imaging new field for another 90 s and so on. This imaging regime was repeated 9 times within 30 min to cover the entire slow cycle. E.g. to cover the entire slow cycle of 30 min with 3 s time interval, first set of movies were started for new field of cells after 3, 6, 9, 12, 15, 18, 21, 24, 27 min after copper induction and acquired for 90 s. Remaining 90 s (between 3 min and 6 min time points, between 6 min and 9 min time points, and so on) were used for moving to the next field of cells. Second set of movies were acquired for new field of cells after 4.5, 7.5, 10.5, 13.5, 16.5, 19.5, 22.5, 25.5 and 28.5 min after copper induction and acquired for 90 s to compensate the missing time points from the first set. Similarly, time-lapse movies for 12 s time interval were recorded for 5 min, followed by changing the field of view within the next 1 min. A first set of movies was started after 3, 9, 15, 21, 27 min after induction. A second set of movies was stared at 6, 12, 18, 24 min after induction. For the photobleaching correction, strain YTK1231 was imaged under the same condition with 3 s time interval.

**Table S1.**
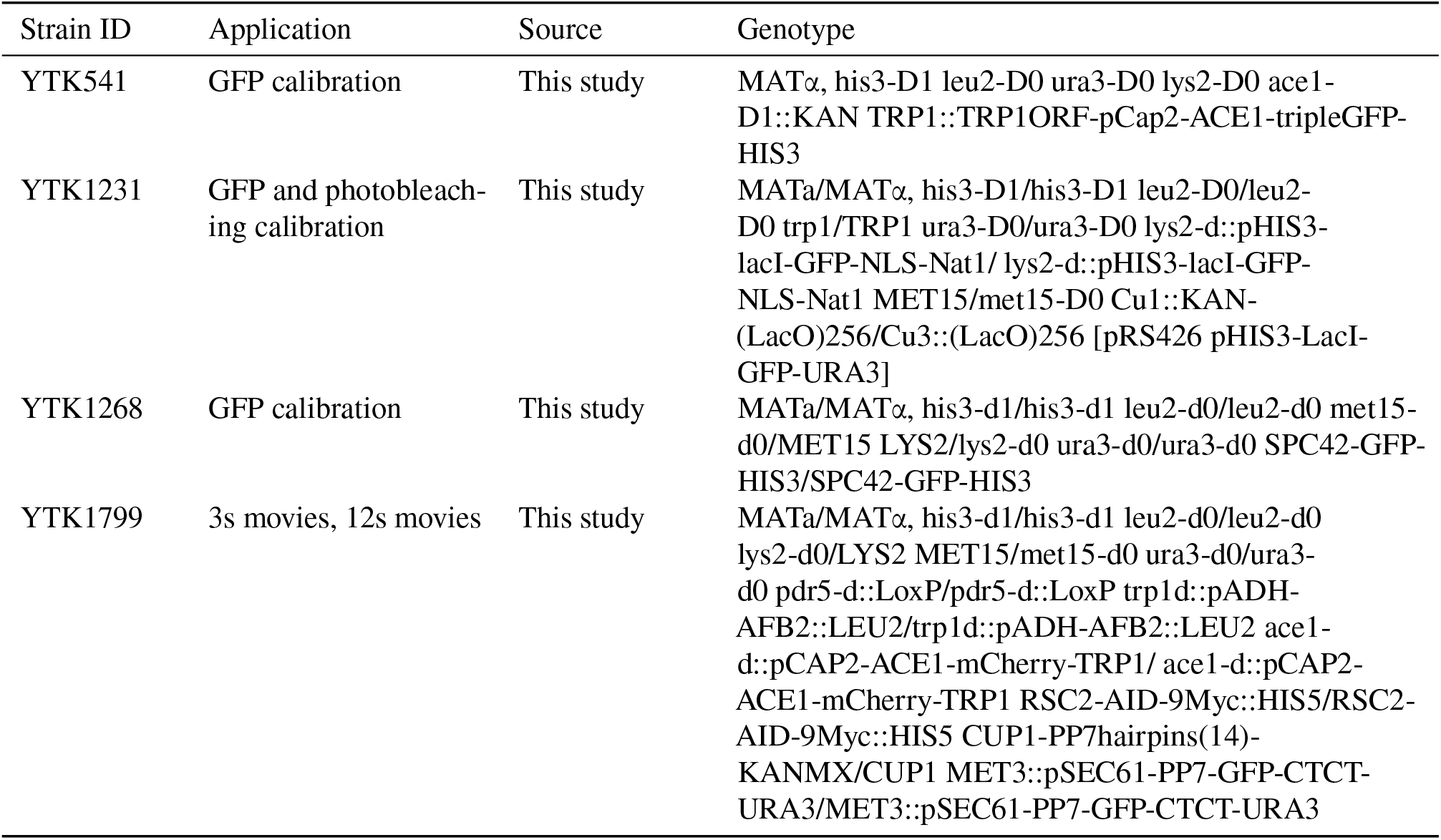
Yeast strains used in this study

**Table S2.**
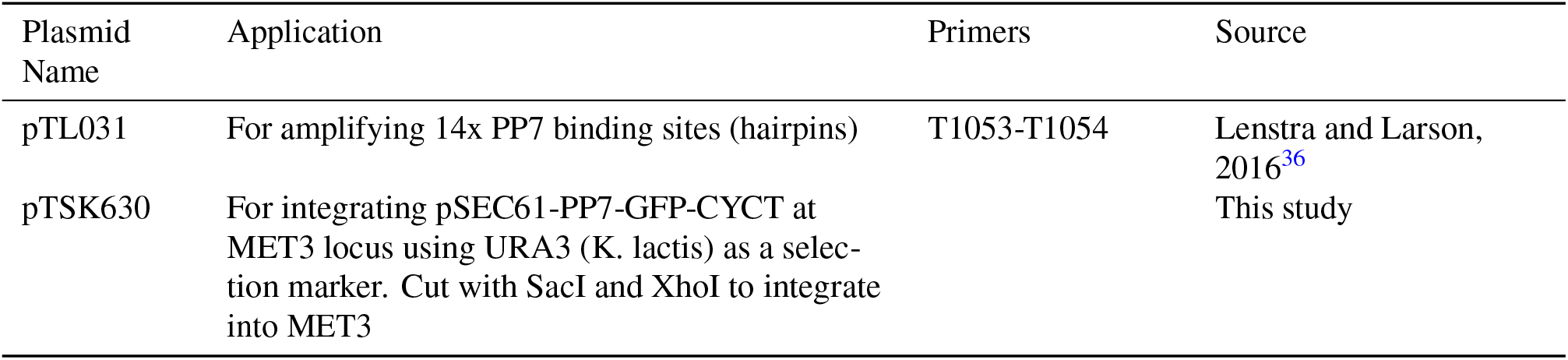
Plasmids used in this study

**Table S3.**
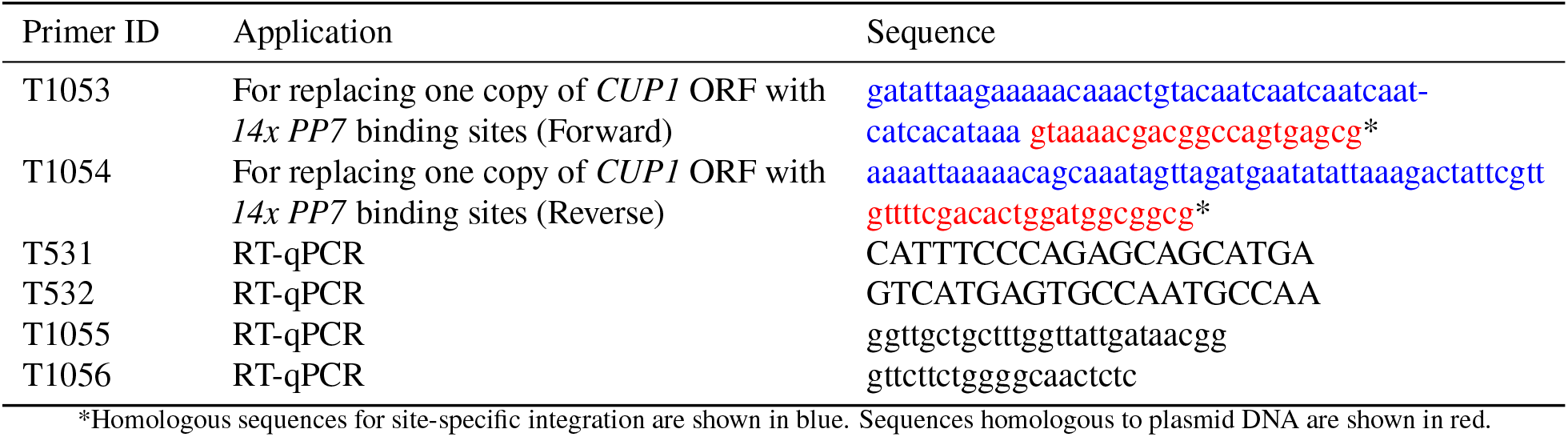
Sequences of the primers used in this study

**Table S4.**
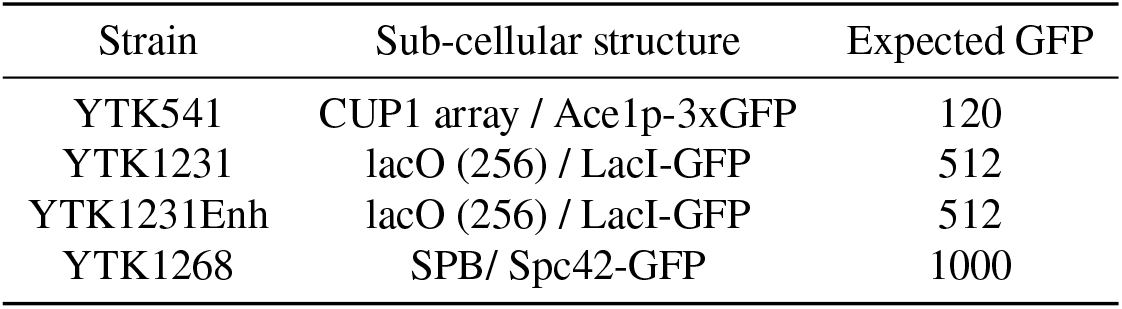
Structures used for GFP calibration

**Table S5.**
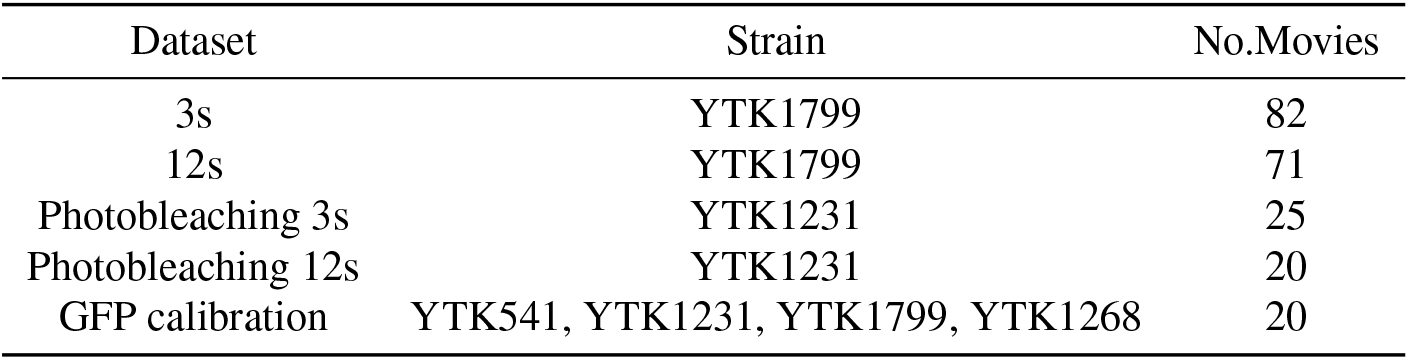
Datasets collected in this study

### S2 Data acquisition and pre-processing

#### S2.1 Cell selection

As a first step, we extracted movies of individual cells from the full movies. In order to extract cells with spots, we computed two projections: a maximum projection over all frames leading to a 2D movie and a maximum projection over all time frames and slices producing a single image. The latter single image allowed to identify all cells that develop an active transcription site during imaging. For the identified cells we drew ROIs in the initial time frame of the 2D video. These initial ROIs were tracked over time and used to extract 3D movies of individual cells. For each cell, the full image stack over time was stored for trace extraction. Fluorescence intensity traces were obtained as described in Sec. S6.2. A more detailed description of the custom tracker is given in Sec. S12.

#### S2.2 Trace extraction

To track fluorescence spots and quantify fluorescence levels, we developed a custom 3D method based on sequential filtering. The location of the spot within an image stack at time *t*_*k*_ is given by *r*_*k*_ = (*x*_*k*_, *y*_*k*_, *z*_*k*_). The spot is modeled as a diffraction limited point source such that the intensity 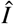 of the pixel at position *r* can be described as

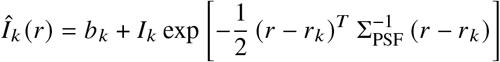

where the diagonal matrix Σ_PSF_ describes the shape of the point spread function of the optical system, *b*_*k*_ is a local background and *I*_*k*_ is the peak intensity of the spot that is related to the underlying intensity of the point source by a constant factor. In addition, we introduce a binary variable *s*_*k*_ that describes whether the spot is visible in frame *k*. This leads to a total state *x*_*k*_ = (*r*_*k*_, *b*_*k*_, *y*_*k*_, *s*_*k*_). The observation model is given by the predicted intensity per pixel with additive Gaussian noise. State estimation is carried out using the standard recursive filtering approach^43^. Due to the non-linear observation model, a Laplace approximation is used to evaluate filter updates. The details are given in S12. The filter provides estimates of *I*_*k*_ for *k* = 1, …, *n* which are treated as the noisy measurements *y*_1_, …, *y*_*n*_ for the main analysis. Traces showing dividing spots due to DNA replication were excluded from further analysis. In this study we do not take into account the noise from the fraction of the cells that completed the replication of the reporter gene, but did not separate sister chromatids, which may lead to transcription of two alleles in the same spot.

#### S2.3 Quality Control

In the images, some of the cells showed too high or too low GFP expression or were otherwise malfunctioning. In the 12s data set, we identified 3260 cells where an active TS is visible in at least one time frame during imaging. After tracking, all traces were compared manually to the corresponding cell video for quality checking. During this screening, we removed all potentially problematic traces, e.g. floating or dividing cells. The remaining 3036 high quality traces were used for further analysis. The same procedure was applied to the 3s data set leading to a selection of 3685 of an initial 4053 cells. A detailed summary of extracted cells per time interval is given in Table S13.

**Table S6.**
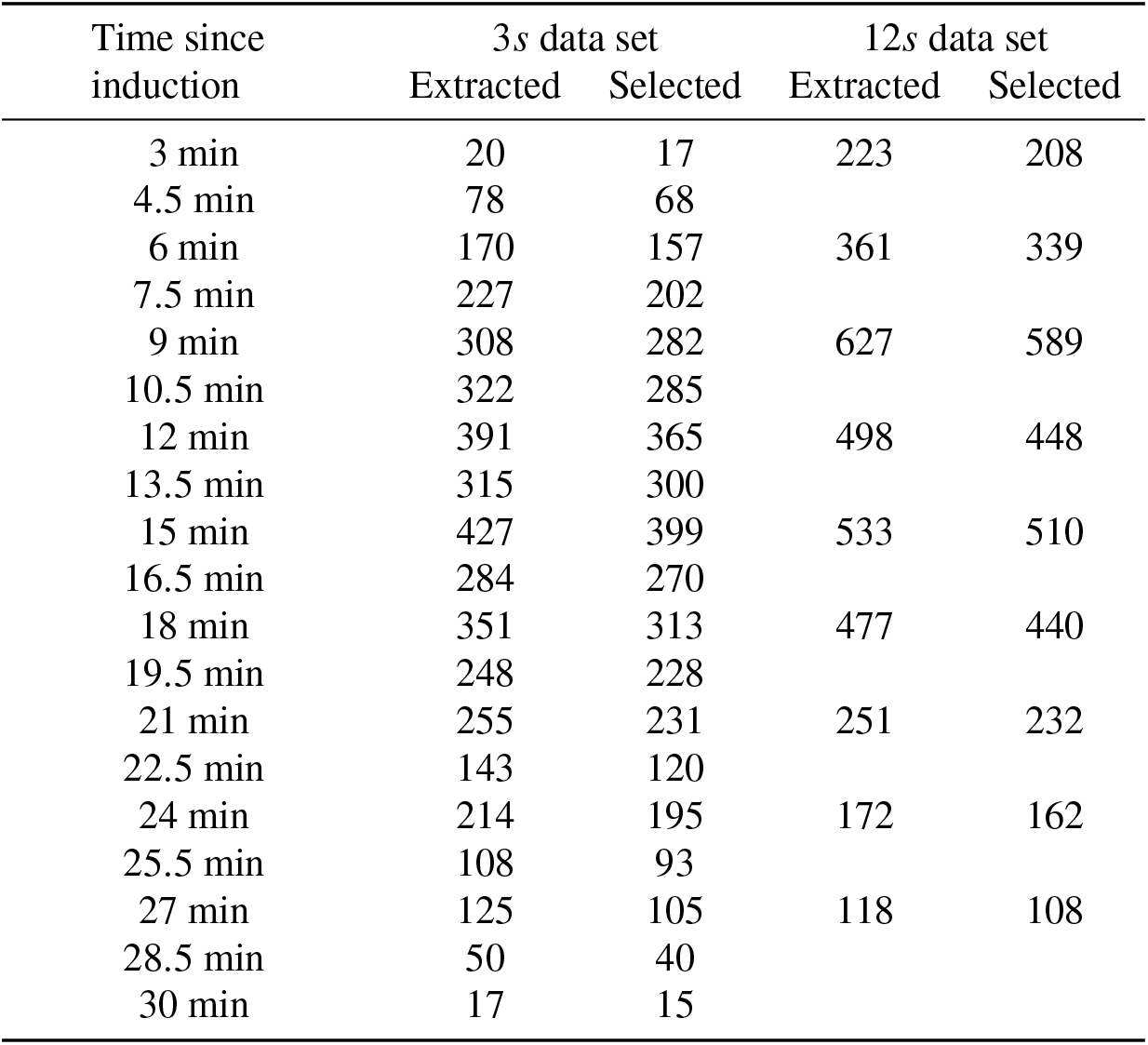
Cells extracted from the 3 *s* and 12 *s* data sets

### S3 Exploratory analysis

Before turning to the model-based Bayesian inference approach, we performed descriptive statistical analysis to confirm the cyclic activity of the *CUP1* promoter known from earlier studies.

#### S3.1 Total image brightness

The movies within different dataset were collected on different days. To ensure that these movies are comparable, we check the total intensity distributions of the first time frame of every movie. By focusing on the first time step, we can avoid confounding effects of bleaching. Boxplot representations of the intensity distributions (Fig. S11) reveal some variations in the intensity distribution. To understand these variations, it is important to note that the images consist mostly of extra-cellular background. Biological differences mainly affect the nuclear brightness and can thus not be responsible for large distribution shifts. Possible factors that could affect total brightness are the agarose medium in the background or variations in illumination or other factors of the optical system.

The most striking difference is that the movies collected on day 4 in the 3s dataset are significantly brighter than all other movies in this dataset. To make the dataset more homogenous, we exclude the day 4 movies from all further analysis. We also observe that the 12s dataset is significantly brighter than the 3s dataset. The reason for this difference is unclear as several months are between the recording dates. However, as we analyze the dataset separately and optical factors such as the gfp scaling are inferred from the dataset, we do not view this as a problem. The third observation we discuss here is that the brightness tends to decrease for movies take from the same coverslip. This is most likely due to some cross-bleaching from imaging neighboring positions. Since the effect is rather small, we will ignore it in the following analysis.

**Figure S6.**
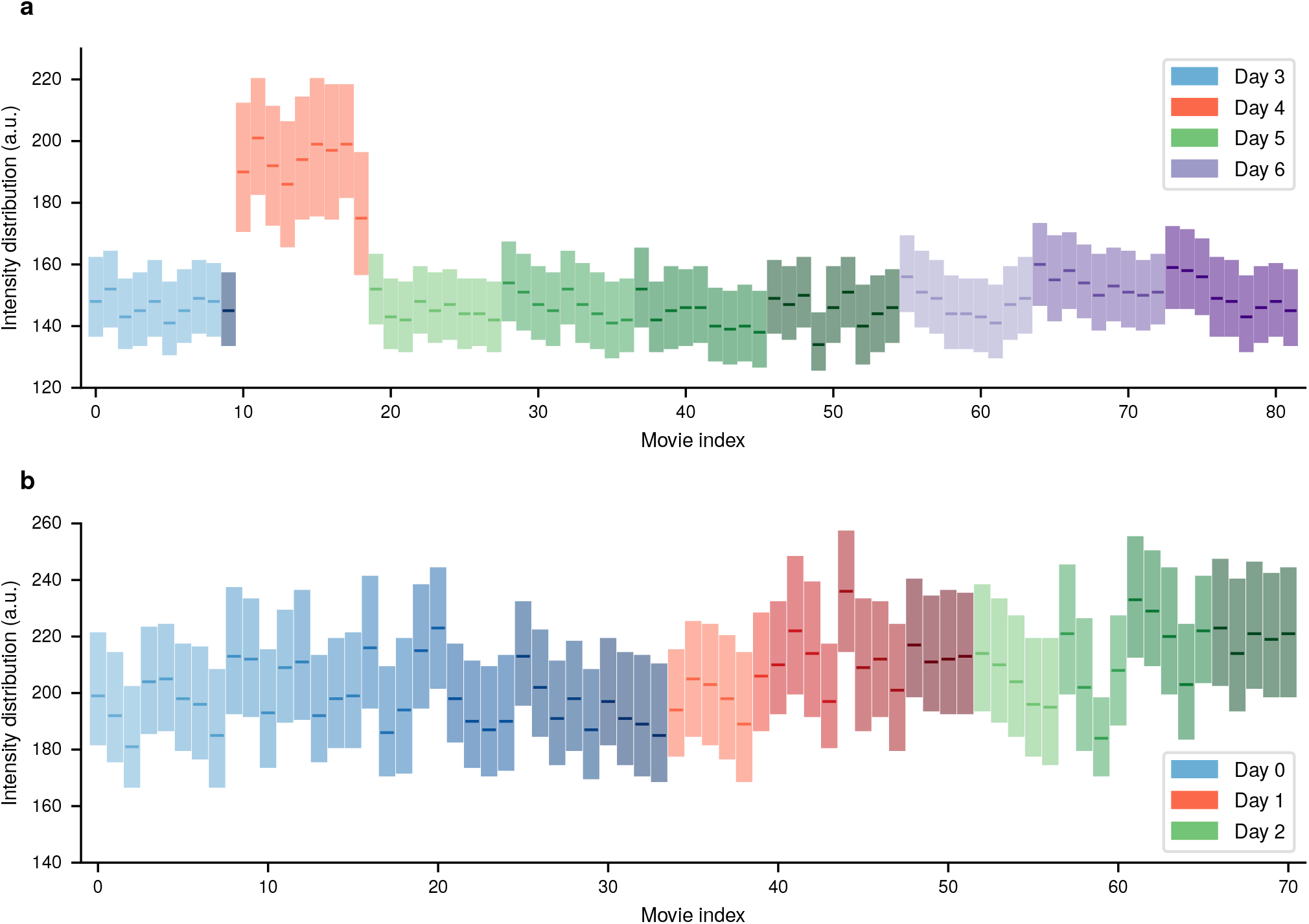
Pixel intensity distributions for the first time frame of every movie in form of a box plot shown for the 3 s dataset (**a**) and the 12 s dataset (**b**). Lower and upper border of the rectangles represent 25th and 75th percentile, respectively. The bold line within the box indicates the median. Every color family represents one experiment day, while the shading withing a color family indicates movies taken from the same coverslip

#### S3.2 Spot Brightness

We now take a first look at the spot brightness. Similarly as an the previous section, we consider only the first frame of every movie to avoid effects of bleaching. For every movie, the spot intensities of all active cells in the first frame were pooled. The corresponding distributions are shown in the form of boxplots for both 3s and 12s dataset in Fig. S12.

Especially in the 3s dataset one can observe a clear dependence of spot brightness on the time since induction. Since the movies taken from the same coverslip are ordered sequentially, they correspond to spot measurements every 3 minutes or every 6 minutes for 3 s dataset and 12 s dataset, respectively. The observed sharp increase followed by a longer decrease is well in accordance with measurements from RT-qPCR. In addition, the results from different coverslips are quite similar and show less variation than the total intensity distributions in Fig. S11. The reason for this is that spot intensities are measured relative to local nuclear background such that additive noise affecting the total brightness of images is filtered out by the spot tracking. The one exception from this are the movies from day 4, which we have already decided to treat as outliers due to their much higher overall brightness.

The dependence on time since induction is also visible in the 12 s dataset albeit less clear than for the 12 s dataset. One reason for this is that there are fewer measurements per coverslip due to the longer duration of the individual movies. In addition, some series of the movies are shifted by 3 minutes leading to a less regular appearance.

Finally, it seems that the spots in the 12 s dataset are overall brighter than in the 3 s dataset. Two possible explanations come to mind. The first possibility is that the cell cultures differ in activity which is possible due the time between the collection of the datasets. Alternatively, multiplicative noise affecting the total intensity distribution could explain the differences as it would not be filtered out by the spot tracking. In practice, it could also be a combination of both factors. We deal with this by estimating the observation model (cf. Sec. S9) parameters individually for every dataset. Any multiplicative noise will be captured by the estimated GFP scaling factor such that remaining differences can be attributed to Biological causes.

**Figure S7.**
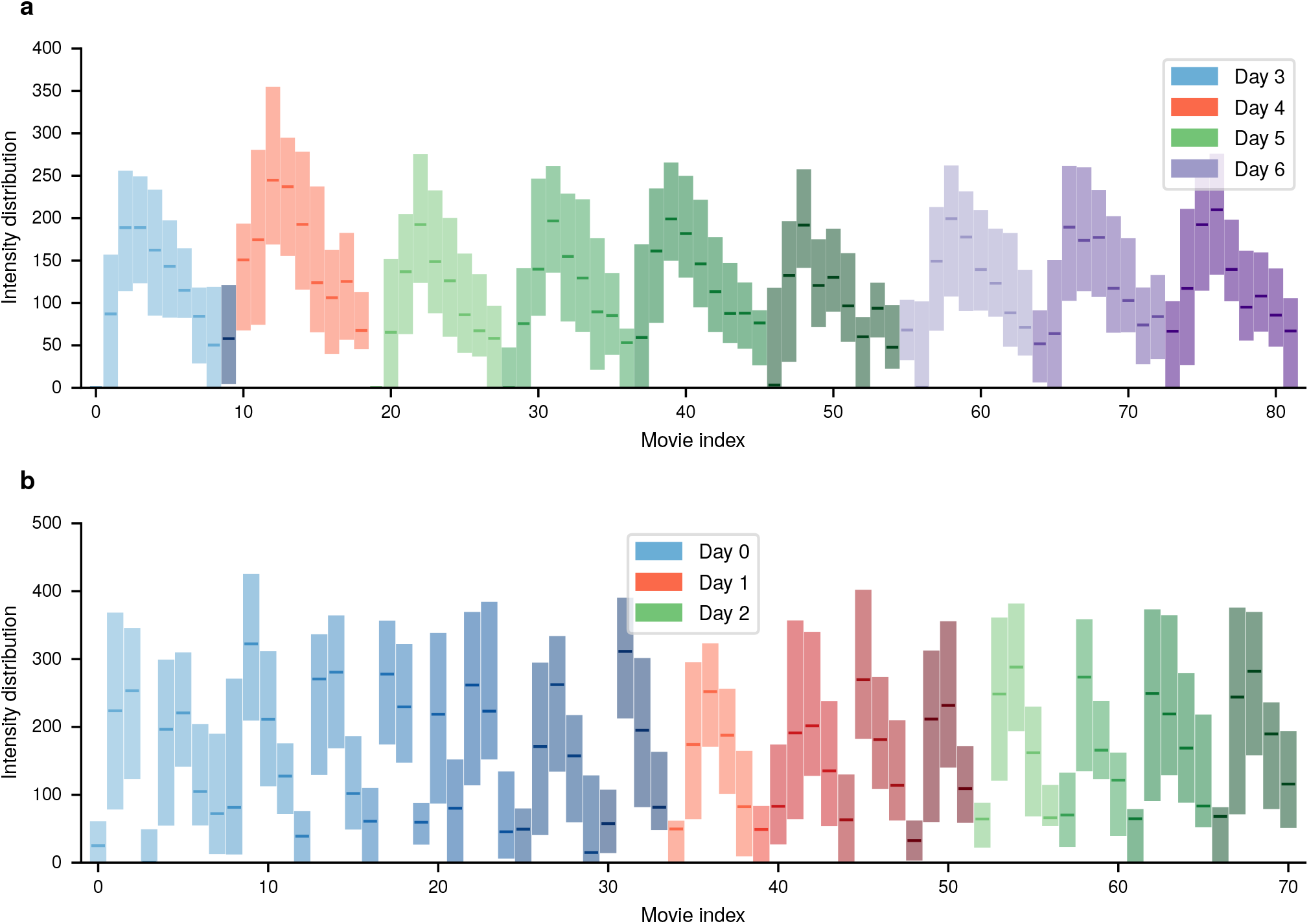
Pixel intensity distributions for the first time frame of every movie in form of a box plot shown for the 3 s dataset (**a**) and the 12 s dataset (**b**). Lower and upper border of the rectangles represent 25th and 75th percentile, respectively. The bold line within the box ndicates the median. Every color family represents one experiment day, while the shading withing a color family indicates movies taken from the same coverslip.

#### S3.3 Investigating time-dependant activity

The presence of non-stationary transcription dynamics can be already seen from the movie histograms in Fig. S12. We will now take a closer look by inspecting the average spot intensities pooled over all movies that started with the same time delay with respect to induction. As before, we will only consider the first time frame of each movie to avoid the effects of bleaching. The first row in Fig. S13 shows the average spot intensity over the cycle. For both 3 s dataset (left) and 12 s dataset (right) wie see a sharp rise of the intensity in the early part of the observed interval followed by a slower decay. While the general shape of both curves is similar, the spots in the 12s dataset are generally brighter as we have seen from the per-movie distributions (cf. Fig. S12). The second row of Fig. S13 shows the number of responding cells divided by the total number of cells per time window. To assign cells to the class of responders or non-responders, we used a standard Gaussian mixture classifier on the spot intensity distributions of the datasets. In order to avoid bias by the overall different brightness, we applied this classifier individually on each dataset. The resulting responder ratios of both datasets agree well.

**Figure S8.**
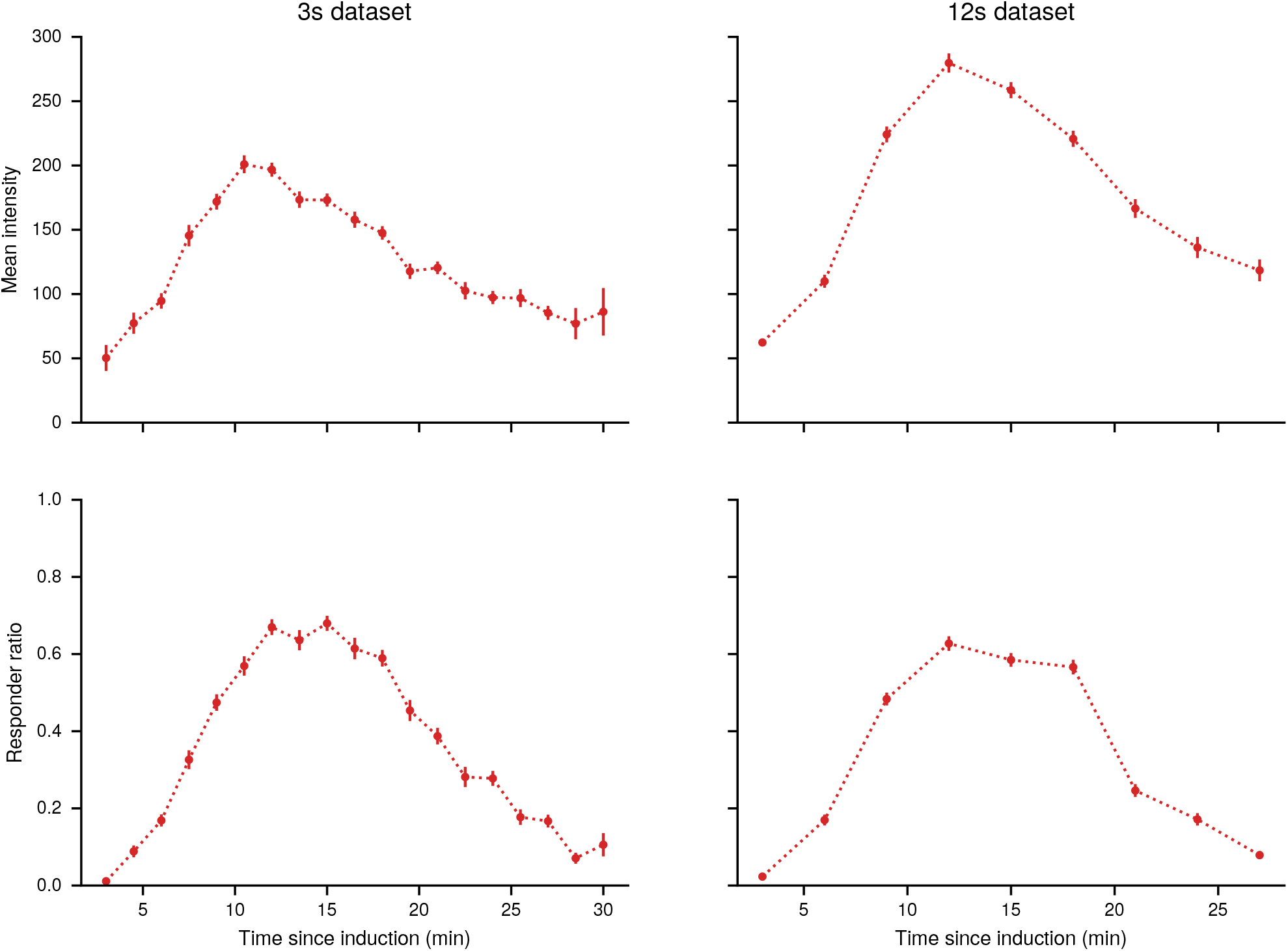
Dependence of summary statistics on the time since induction. The first row shows the mean of the spot intensities pooled over all videos with the same time delay since induction. The second row shows the number of responders divided by the total number of cells per time window. Error bars indicate standard error. Results for the 3 s dataset are also shown in the main text (Fig. 1f, g)

### S4 Model

#### S4.1 Kinetic Transcription Model

As described in the main text, the model is an example of a Markov jump process that satisfies the master equation (1). The transition function *Q* defines the probability of an event to happen in an infinitesimal interval *h*

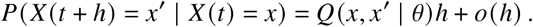

The transition function is fully specified by the vector of parameters *θ* = (*k*_on_, *k*_off_, *k*_i_, *k*_e_, *k*_t_)^⊤^ and the conditions under which transitions can occur, leading to

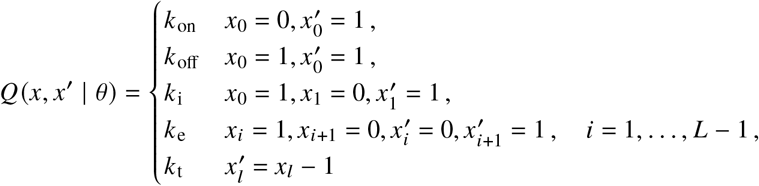

For parameter inference, we need to evaluate the system for many different parameter configurations. It is therefore convenient to represent the transition functions as

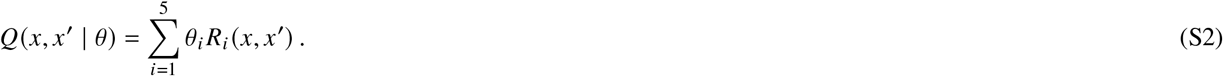

The operators *R*_*i*_ are independent of the parameters. By enumerating the states of the system, the probability *p* (*x, t*) can be represented by a vector **p**(*t*) and the transition function *Q* becomes a sparse matrix **Q**. With this, the master equation becomes

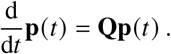

A formal solution of this system is given by the matrix exponential

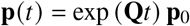

with initial distribution **p**_0_. For sparse **Q**, this can be efficiently solved for fairly large state spaces by the Krylov subspace approximation for matrix exponentials^41, 42^.

#### S4.2 Observation Model

The kinetic model discussed above is a continuous time model. In practice, one cannot observe such a systems in continuous time but rather at discrete sample times *t*_1_, …, *t*_*n*_. In addition, we do not observe the process *X*(*t*_*k*_) directly. Here, our measurement *Y*(*t*_*k*_) is provided by the total intensity of the fluorescence spots as measured by the tracking algorithm. In the following, we construct a model that relates *X* and *Y*. First, note that as the polymerase traverses the gene, an additional stem loop is added for every site until at some point the maximum number of stem loops is acquired. For the remaining part of the transcription process, the number of stem loops stays constant. After termination, the mRNA is released and rapidly diffuses away from transcription site. The corresponding spot is thus no longer visible and we observe a sharp drop in intensity. Hence, if *a* ∈ ℕ^*L*+1^ encodes the number of stem loops associated with the sites of the gene, the variable

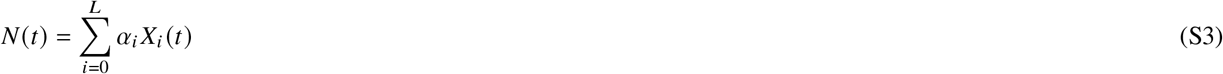

corresponds to the number of visible stem loops at time *t*. Assuming that stem loops are occupied by GFP fast compared to the elongation speed, the total spot signal can be described as

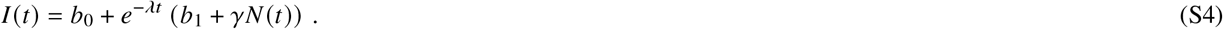

Here, *λ* is the bleaching factor and *b*_0_, *b*_1_ correspond to baseline background and a bleachable part of the background respectively. The factor *γ* encodes the intensity contribution per GFP molecule. During image acquisition and intensity estimation, the signal is corrupted by various forms of noise such as z-diffusion of the transcription site, photon counting noise on the camera chip, variations in the media, mismatch of the point-spread function with the Gaussian approximation, irregular background illumination, etc. We subsume all these effects into a single multiplicative noise variable leading to the relation

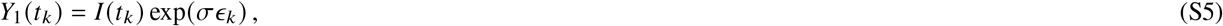

where *ϵ*_*k*_ are independent and standard normally distributed. While (S10) is a reasonable approximation for larger signals, it is not suitable for very small signals. The reason for this is that at low intensity, due to fundamental limitations of the spot detection, there is an increased probability that a spot is missed or that a local fluctuation is confused with a signal. To take account of this effect, we introduce the random background signal

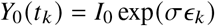

and an additional unobserved random variable *Z* (*t*_*k*_) ∈ {0, 1} with

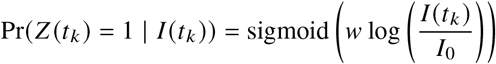

where sigmoid(*x*) = (1 + exp(−*x*))^−1^ and *w* is a response parameter. The final observation model is then given by

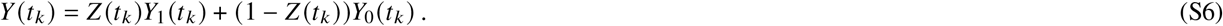

This can be understood as a soft threshold for spot detection. When the true spot intensity *I* is larger than *I*_0_, *Z* will likely be one and we measure the signal *Y*_1_ with high probability. When *I* is smaller than *I*_0_, *Z* is likely zero and we observe the spurious signal *Y*_0_ with high probability. The likelihood corresponding to (S11) is that of a mixture of two lognormal distributions with parameters log(*I*) and log(*I*_0_). An overview of all parameters involved in kinetic and observation model is given in Table S14.

#### S4.3 Elongation times

Many models for transcription assume independent movement of individual polymerases. This leads to a simple relation between elongation rate and elongation time as 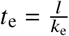 where *l* is the size of the translated region. Due to possible interactions between polymerases, this relation is not valid in the TASEP model. As shown in Fig. S14, the TASEP model requires a higher value compared to an independence model to produce the same expected elongation time. This difference becomes smaller for higher elongation rates, since a fast movement of individual polymerases decreases the probability of interaction. As an example, consider the dotted black line in Fig. S14 corresponding to an elongation time of 12 s. To produce such an elongation time, the independent model requires an elongation rate of 100 nt s^−1^, while the TASEP model requires and elongation rate of roughly 120 nt s^−1^.

**Figure S9.**
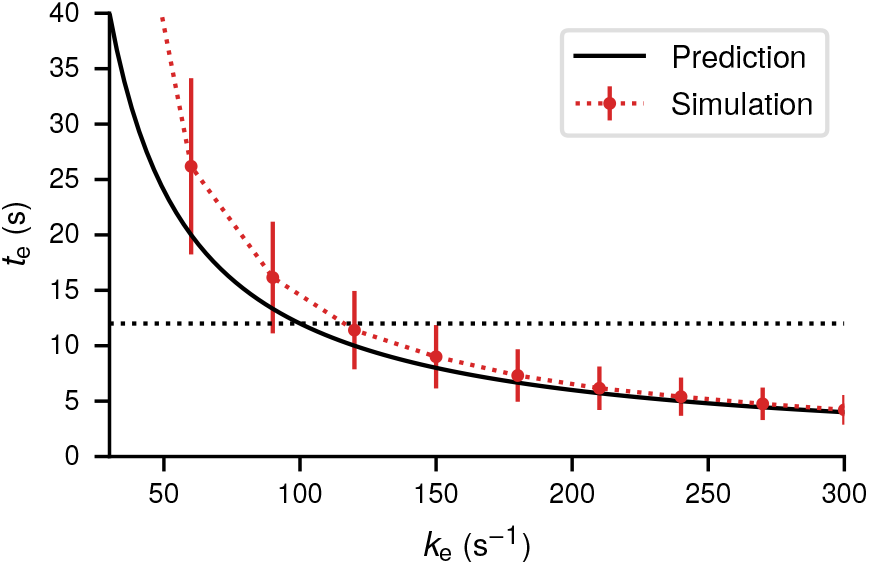
Expected elongation times of a gene template with size *l* = 1200 nt for different values of the elongation rate *k*_e_. The thick black curve indicates the value obtained from independent polymerase movement by the relation 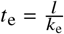. Red dots show the expected elongation time from simulations of a TASEP model with a site size of 120 nt. Error bars indicate the standard deviation of the distribution. Initiation rate *k*_i_ and termination rate *k*_t_ are fixed and chosen such that typically multiple polymerases are on the template and no traffic jams are caused by the termination site.

#### S4.4 Coarse-graining

The most natural quantization of the gene into sites would be to associate every site with a single nucleotide. This would, however, lead to more than 1000 sites and since the state space of the model scales as 2^*L*^ would make inference intractable. In addition, we only observe the system by one-dimensional summary statistic every few seconds such that most of the detailed dynamics are not captured. It is thus necessary to combine several nucleotides into a single site. A good candidate for such a coarse-graining is the DNA footprint of a stem loop, which is roughly 60 nt, as the appearance of a stem loop is the most fine-grained observable. For a DNA template consisting of 1200 nt, this would lead to *L* = 22 sites. While the master equation this model is still tractable, it requires significant computational effort such that only a small number of cells can be handled this way. As argued in the main text, it is important to pool many traces in order to overcome structural identifiability limitations of the system. As a compromise between tractability and resolution, we choose a partition size of 120 nt corresponding to two stem loops leading to a system of *L* = 12 sites.

The coarse-graining changes the waiting time distribution between appearance of two stem loops. If one starts from a fine-grained model where every site corresponds to a single nucleotide, the waiting time for jumps of size 120 bp is much more peaked compared to the exponential distribution. To investigate the robustness of the inference against this kind of mismatch, we simulated 100 trajectories from a fine-grained model described in^23^ Supp. M. The model is similar to the TASEP model we use for inference, but uses one site for every nucleotide. In addition, RNAP molecules have a footprint of 40 sites and individual stems loops appear every 60 sites with 14 stem loops in total. The number of sites was set to 1200 roughly corresponding to the gene investigates experimentally. The elongation rate was set to *k*_elong_ = 100 nt s^−1^, for all other parameters we used the standard values of the coarse-grained model. We generated 100 trajectories with 3 s time-lapse and performed pooled inference. For simplicity, we assume a constitutive promoter an drop the switching site. The results are presented in Fig. S15a. They indicate that while initiation and termination rate are inferred quite accurately, the estimated elongation rate is, however, significantly larger than the ground truth used to generate data in the fine-grained model. This can be explained by the exclusion property of the TASEP process. In the coarse-grained model, collisions will happen more frequently on average since the space is limited. In order to achieve a similar total elongation time, the rate has to be increased to account for the possible blocking. To verify this, we generated trajectories from fine-grained model and coarse-grained model and extracted the distribution of total transcription times (Fig. S15b). The mean elongation time predicted by the learned coarse-grained model is quite close to the mean elongation time of the fine-grained model. This demonstrates that even though there is a mismatch, the model learns physically meaningful quantities. However, we stress that this is merely an example to illustrate model mismatch. In practice, the distribution of elongation times predicted by the fine-grained model is likely too narrow as it ignores effects such as pausing, reverse steps, chromatin remodelling, etc. Thus, a coarse-grained model with wider waiting time distribution may be a better approximation to the real data, as long as these effects are not modeled explicitly.

#### S4.5 Overview of the model parameters

**Figure S10.**
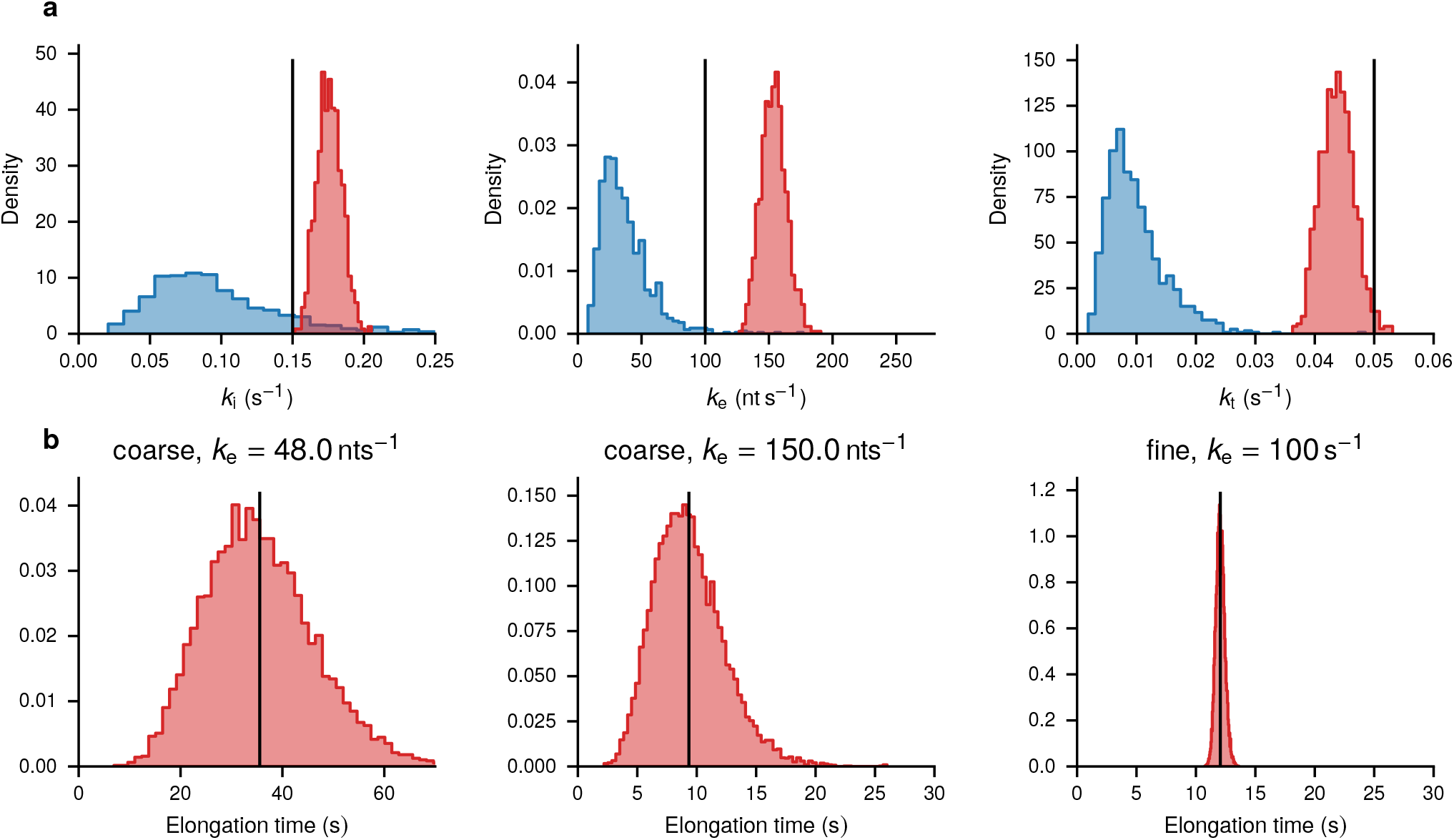
**a** Pooled posterior inference of a non-switching model on traces generated by the fine-grained TASEP model with a time laps of 3 s. Observation parameters are not shown but were also estimated during inference. The rows show histogram approximations of the prior distribution (blue) and the posterior distribution (red) for the model parameters. Black lines indicate the parameter value used to generate the data. **b** Distribution of total elongation times obtained from forward simulation of the coarse-grained model (left and middle) and the fine-grained model (right). The elongation right of the left plot is a typical value of the prior distribution, the elongation rate of the middle plot is a typical value of the posterior distribution. Black lines indicate the empirical mean.

**Table S7.**
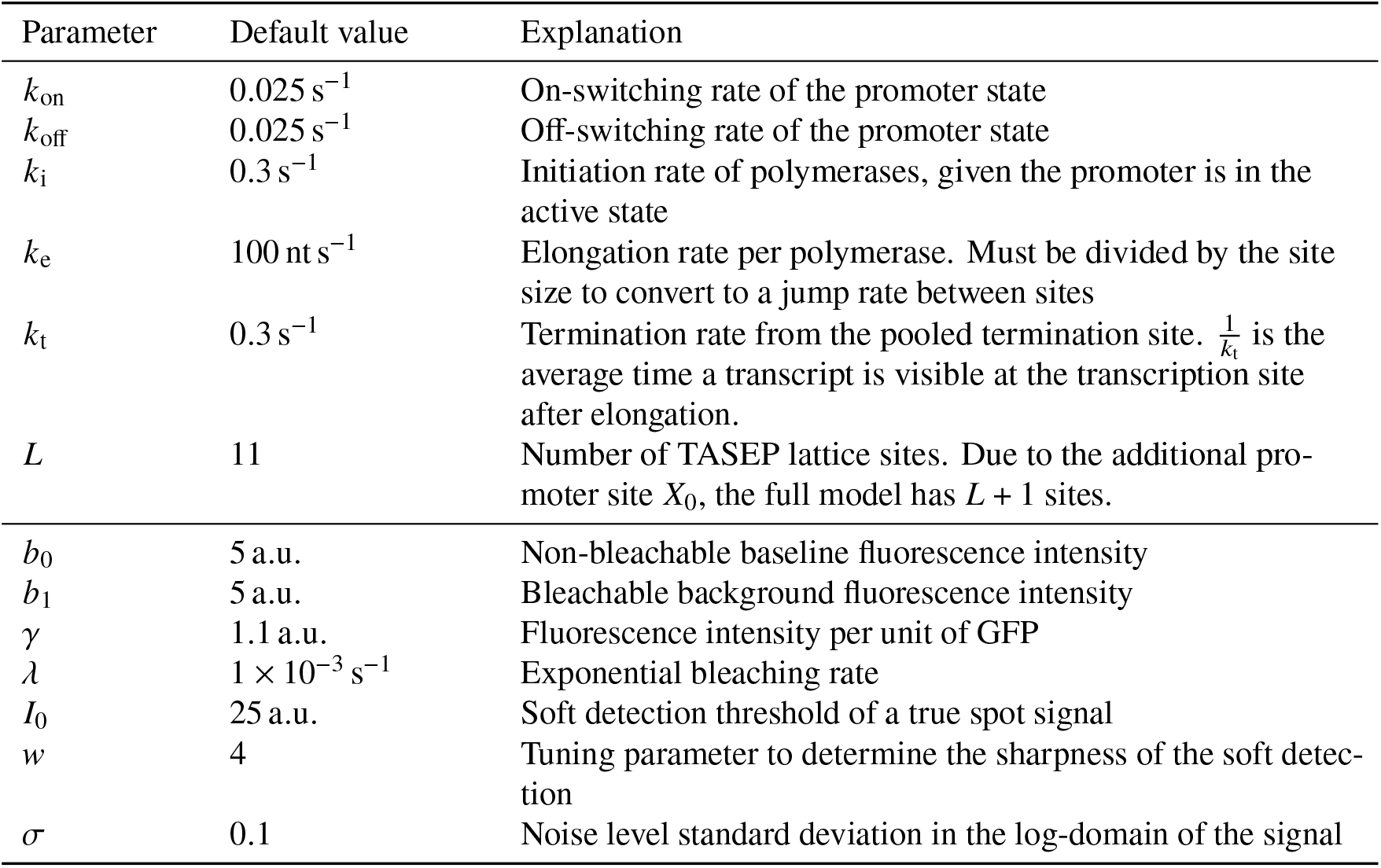
Overview of the model parameters

### S5 Experimental details

#### S5.1 Yeast strains and plasmids

For live transcript analysis, we engineered the haploid strains of *Saccharomyces cerevisiae* (BY4742 and BY4741), which are isogenic to S288C (Research Genetics/Invitrogen, Huntsville, AL). 14X PP7 binding sites (hairpins) were amplified from pTL031^36^ using primers T1053, T1054 and integrated in one of the haploids by replacing one of the *CUP1* ORFs in the yeast genome by homologous recombination^37^. For expressing PP7-GFP coat protein, we constructed a MET3 integrative vector (pTSK630) to express PP7-NLS-GFP from *SEC61* promoter and integrated this vector by SacI XhoI digestion in both the haploids. This vector can be available upon request. For the photobleaching correction, we used a diploid strain YTK1231 in which both the *CUP1* arrays are replaced by 256 copies of *LacO* and lacI-GFP-NLS is expressed from *pHIS3*. For preparing the calibration curve for the number of GFP molecules verses brightness/intensity, we used three yeast strains (YTK541, YTK1231 and YTK1268) with known numbers of GFP molecules per locus. YTK541, contains a tandem array of 10 copies of *CUP1* locus with 40 binding sites for the transcription activator Ace1p-GFP. *CUP1* is activated by Cu, and at the peak of activity *CUP1* array binds 120 molecules of GFP. In YTK1231, each lacO binding site may be associated with a dimer of the lac Repressor (LacI-GFP)^38^. Therefore, the array of 256 tandem lacO binding sites is associated with 512 LacI-GFP molecules. In YTK1268, the spindle pole body of the diploid yeast strain contains approximately 1000 molecules of Spc42-GFP. Strain genotypes are provided in Table S8, plasmids in Table S9 and primer sequences in Table S10.

#### S5.2 Media and growth conditions

For live transcript analysis, cells of YTK1799 were plated on CSM-URA plate (from −80 °C frozen glycerol stock) and grown for 48 h at 28 °C. 3 to 5 colonies were inoculated in 3 mL CSM-URA media (in 14 mL polypropylene tubes, Cat no. 352059, Falcon, Maxico) and grown for overnight at 28 °C, 230 RPM. 250 μL of this overnight grown culture was inoculated in 25 mL CSM-URA (in 250 mL flask) and grown at 28 °C, 230 RPM for 24 h. This flask was removed from the shaker and kept in refrigerator at 4 °C. We used this refrigerated culture for daily inoculations for a month to get consistent results (to avoid day to day variations in transcription induction kinetics due to difference in the age of the culture). From this refrigerated culture, we inoculated 60 μL in 3 mL of fresh CSM-URA media (in 14 mL polypropylene tubes, Cat no. 352059, Falcon, Maxico) in the morning and grew the cultures for 5 h at 28 °C, 230 RPM. Cells were harvested by centrifugation (2200 RPM for 1 min) and cells were placed under the CSM-URA agarose pad (100 μM CuSO_4_) for imaging. For the photobleaching correction and GFP calibration curve, strains YTK541, YTK1231 and YTK1268 were grown under the same conditions, except YTK541 and YTK1268 were grown in CSM-HIS media.

#### S5.3 Quantitative RT-PCR (RT-qPCR)

Samples were harvested at indicated time points after Cu induction. RNA was extracted (from yeast cells) using the ISOLATE II RNA Mini kit (Bioline, UK, Cat no. BIO-52072). cDNA was prepared using the iScript cDNA synthesis kit (BioRad, Cat no.: 1708891) starting with 1 mg of total RNA. Quantitative real-time PCR (qPCR) was performed as described^27^. For normalization, the expression of the housekeeping gene ACT1 was quantified. Primers used for this quantification are listed in Table S10 (T531, T532 for CUP1 and T1055, T1056 for ACT1). To confirm the absence of contaminating genomic DNA in cDNA preparations, reverse transcriptase negative (-RT) samples were used as a control, which produced the Ct value difference of >10 cycles between “-RT” and “+RT”samples, indicating a negligible amount of genomic DNA contamination in cDNA samples. mRNA extraction, cDNA synthesis, and qPCR were repeated at least twice, and qPCR was performed in duplicates for each experiment. Error bars indicate SEM.

#### S5.4 Microscope settings and imaging conditions

For imaging live cells, 5 h grown cultures were harvested by centrifugation (2200 RPM for 1 min) and 3 μL of cell pellet were placed in Lab-Tek II chambered coverglass (1.5 Borosilicate Gass, Nunc, ThermoFisher Scientific, MA, US), mixed with equal volume of 200 μM CuSO_4_ containing CSM-URA and covered by 1cm x 1cm CSM-URA agarose pad (100 μM CuSO_4_). A timer was started immediately upon mixing the cells with 200 μM CuSO_4_ containing CSM-URA. 3D time-lapse movies were acquired at the room temperature on the DeltaVIsion Elite Microscope, using 100x 1.4 NA oil immersion objective lens, sCMOS camera, FITC filter set (15 ms exposure, Ex 488/27; Em 505/45, Chroma Technology Corp, Bellows Falls, VT), 15 z-steps at every 400 nm, 1x1 binning and 1024x1024 pixels. Time-lapse movies with 3 s time interval were acquired for 90 s, followed by changing the field within next 90 s, imaging new field for another 90 s and so on. This imaging regime was repeated 9 times within 30 min to cover the entire slow cycle. E.g. to cover the entire slow cycle of 30 min with 3 s time interval, first set of movies were started for new field of cells after 3, 6, 9, 12, 15, 18, 21, 24, 27 min after copper induction and acquired for 90 s. Remaining 90 s (between 3 min and 6 min time points, between 6 min and 9 min time points, and so on) were used for moving to the next field of cells. Second set of movies were acquired for new field of cells after 4.5, 7.5, 10.5, 13.5, 16.5, 19.5, 22.5, 25.5 and 28.5 min after copper induction and acquired for 90 s to compensate the missing time points from the first set. Similarly, time-lapse movies for 12 s time interval were recorded for 5 min, followed by changing the field of view within the next 1 min. A first set of movies was started after 3, 9, 15, 21, 27 min after induction. A second set of movies was stared at 6, 12, 18, 24 min after induction. For the photobleaching correction, strain YTK1231 was imaged under the same condition with 3 s time interval.

**Table S8.**
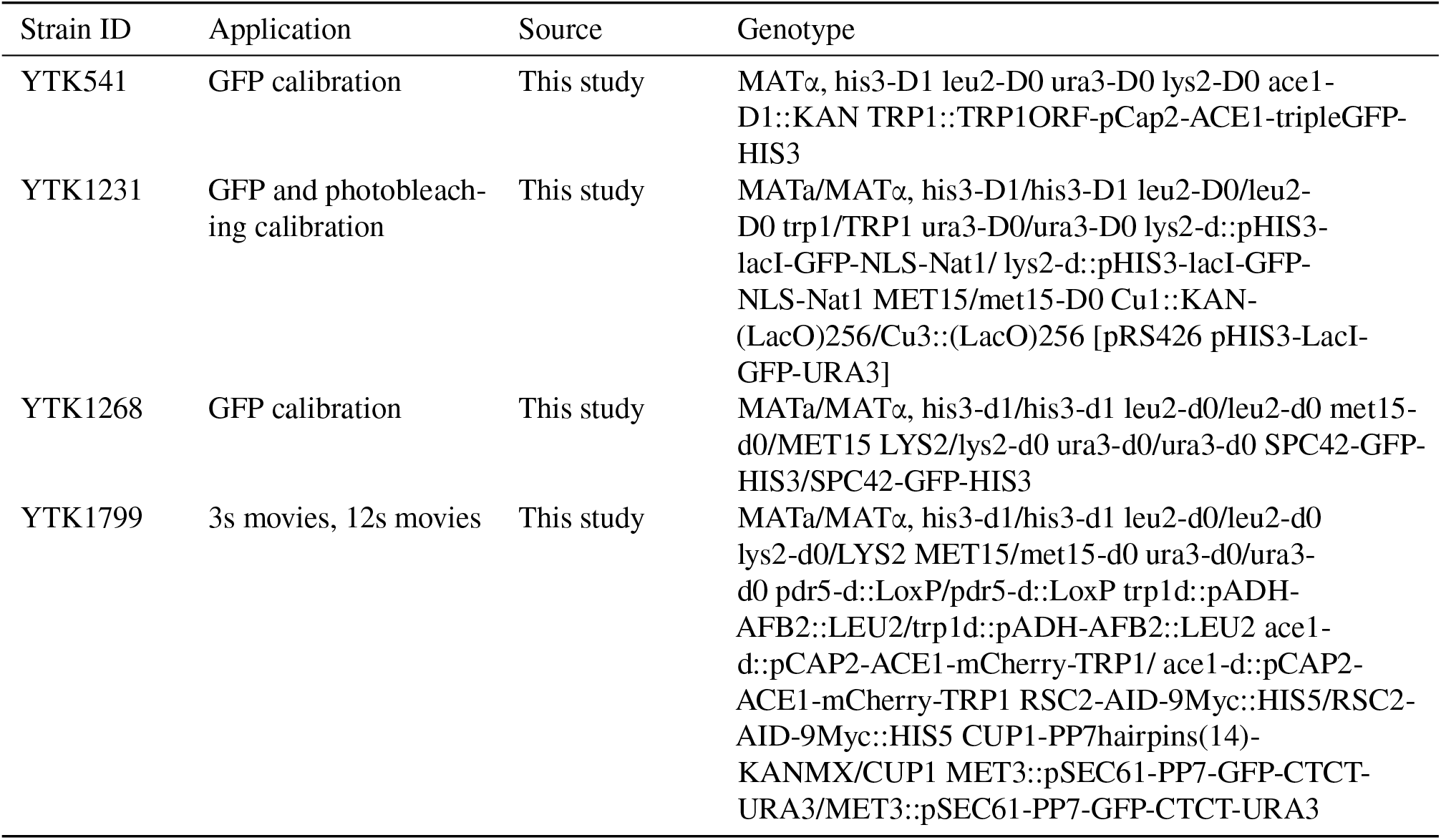
Yeast strains used in this study

**Table S9.**
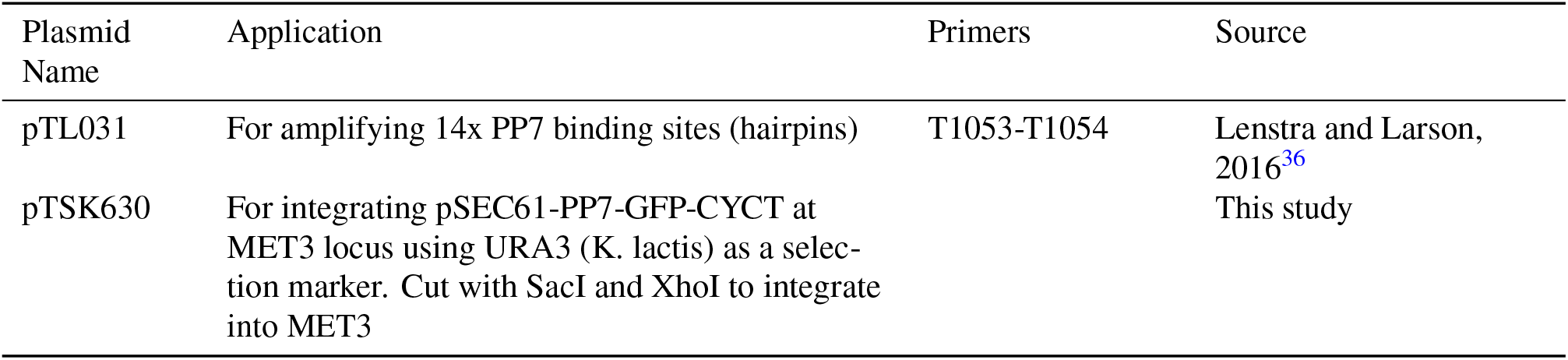
Plasmids used in this study

**Table S10.**
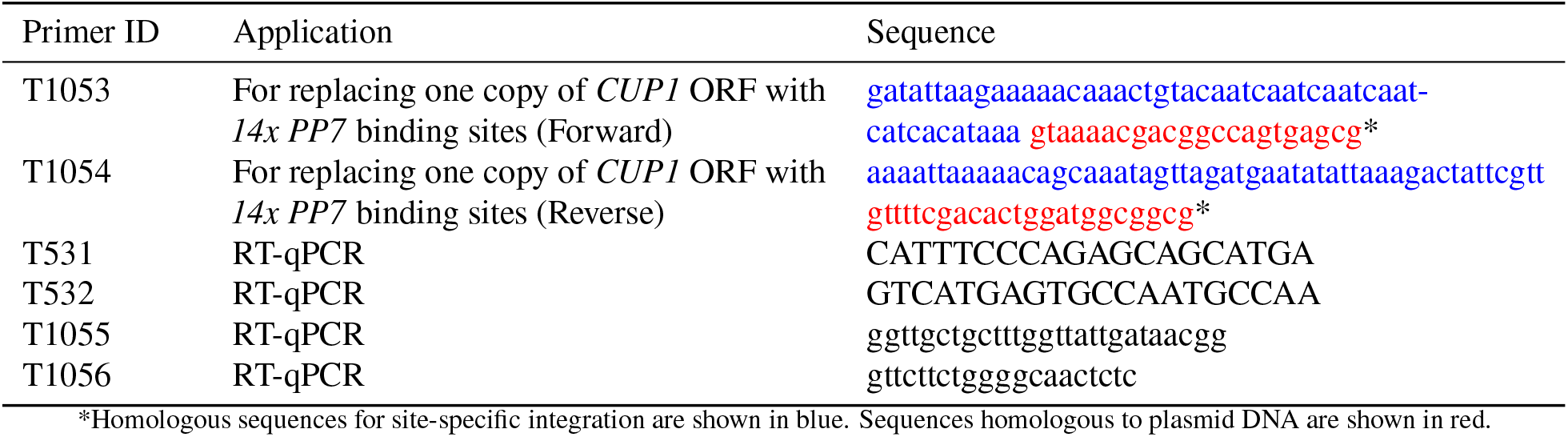
Sequences of the primers used in this study

**Table S11.**
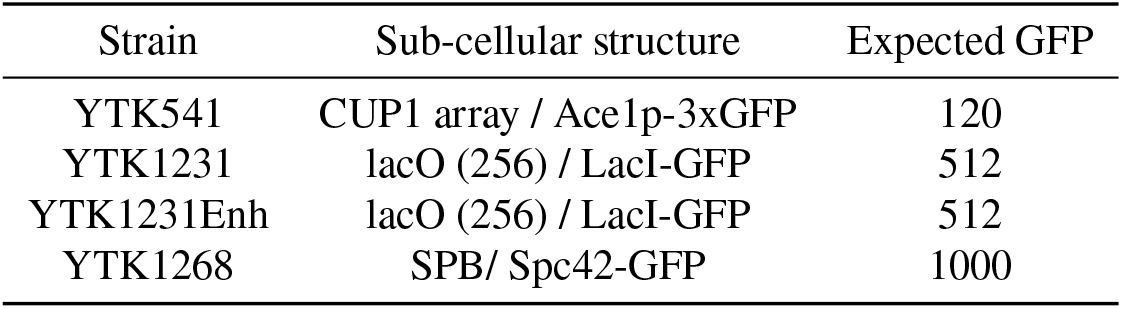
Structures used for GFP calibration

**Table S12.**
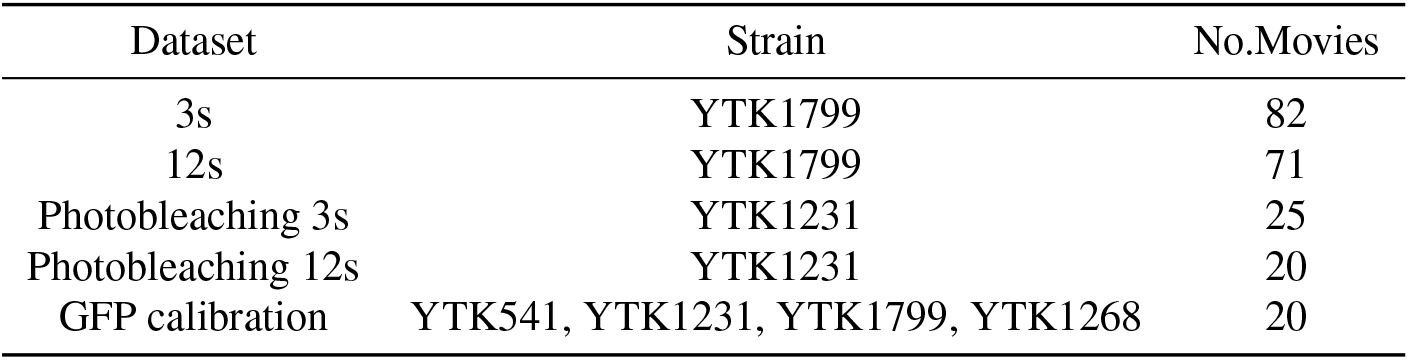
Datasets collected in this study

### S6 Data acquisition and pre-processing

#### S6.1 Cell selection

As a first step, we extracted movies of individual cells from the full movies. In order to extract cells with spots, we computed two projections: a maximum projection over all frames leading to a 2D movie and a maximum projection over all time frames and slices producing a single image. The latter single image allowed to identify all cells that develop an active transcription site during imaging. For the identified cells we drew ROIs in the initial time frame of the 2D video. These initial ROIs were tracked over time and used to extract 3D movies of individual cells. For each cell, the full image stack over time was stored for trace extraction. Fluorescence intensity traces were obtained as described in Sec. S6.2. A more detailed description of the custom tracker is given in Sec. S12.

#### S6.2 Trace extraction

To track fluorescence spots and quantify fluorescence levels, we developed a custom 3D method based on sequential filtering. The location of the spot within an image stack at time *t*_*k*_ is given by *r*_*k*_ = (*x*_*k*_, *y*_*k*_, *z*_*k*_). The spot is modeled as a diffraction limited point source such that the intensity 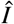 of the pixel at position *r* can be described as

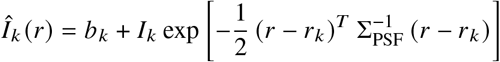

where the diagonal matrix Σ_PSF_ describes the shape of the point spread function of the optical system, *b*_*k*_ is a local background and *I*_*k*_ is the peak intensity of the spot that is related to the underlying intensity of the point source by a constant factor. In addition, we introduce a binary variable *s*_*k*_ that describes whether the spot is visible in frame *k*. This leads to a total state *x*_*k*_ = (*r*_*k*_, *b*_*k*_, *y*_*k*_, *s*_*k*_). The observation model is given by the predicted intensity per pixel with additive Gaussian noise. State estimation is carried out using the standard recursive filtering approach^43^. Due to the non-linear observation model, a Laplace approximation is used to evaluate filter updates. The details are given in S12. The filter provides estimates of *I*_*k*_ for *k* = 1, …, *n* which are treated as the noisy measurements *y*_1_, …, *y*_*n*_ for the main analysis. Traces showing dividing spots due to DNA replication were excluded from further analysis. In this study we do not take into account the noise from the fraction of the cells that completed the replication of the reporter gene, but did not separate sister chromatids, which may lead to transcription of two alleles in the same spot.

#### S6.3 Quality Control

In the images, some of the cells showed too high or too low GFP expression or were otherwise malfunctioning. In the 12s data set, we identified 3260 cells where an active TS is visible in at least one time frame during imaging. After tracking, all traces were compared manually to the corresponding cell video for quality checking. During this screening, we removed all potentially problematic traces, e.g. floating or dividing cells. The remaining 3036 high quality traces were used for further analysis. The same procedure was applied to the 3s data set leading to a selection of 3685 of an initial 4053 cells. A detailed summary of extracted cells per time interval is given in Table S13.

**Table S13.**
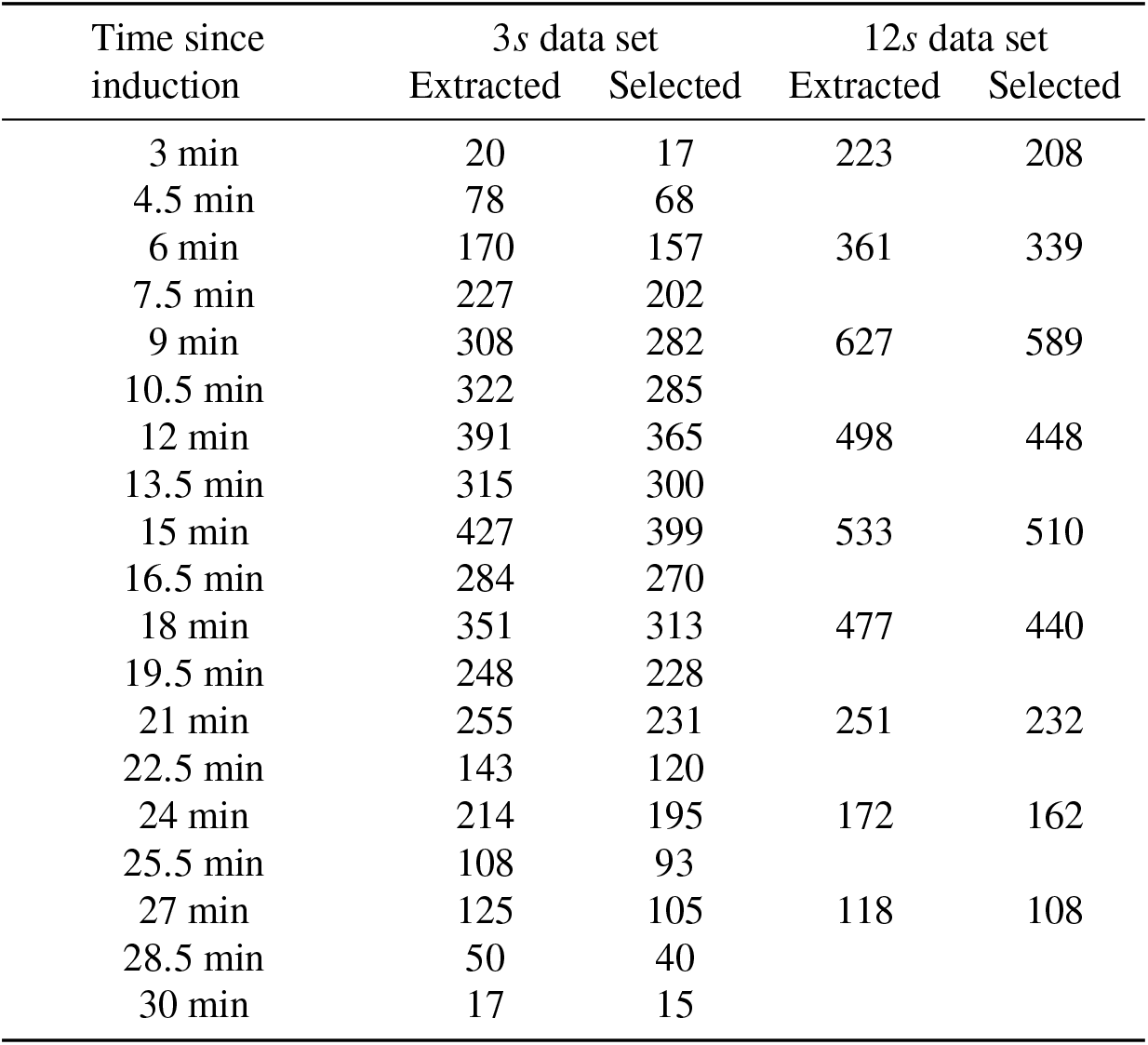
Cells extracted from the 3 *s* and 12 *s* data sets

### S7 Exploratory analysis

Before turning to the model-based Bayesian inference approach, we performed descriptive statistical analysis to confirm the cyclic activity of the *CUP1* promoter known from earlier studies.

#### S7.1 Total image brightness

The movies within different dataset were collected on different days. To ensure that these movies are comparable, we check the total intensity distributions of the first time frame of every movie. By focusing on the first time step, we can avoid confounding effects of bleaching. Boxplot representations of the intensity distributions (Fig. S11) reveal some variations in the intensity distribution. To understand these variations, it is important to note that the images consist mostly of extra-cellular background. Biological differences mainly affect the nuclear brightness and can thus not be responsible for large distribution shifts. Possible factors that could affect total brightness are the agarose medium in the background or variations in illumination or other factors of the optical system.

The most striking difference is that the movies collected on day 4 in the 3s dataset are significantly brighter than all other movies in this dataset. To make the dataset more homogenous, we exclude the day 4 movies from all further analysis. We also observe that the 12s dataset is significantly brighter than the 3s dataset. The reason for this difference is unclear as several months are between the recording dates. However, as we analyze the dataset separately and optical factors such as the gfp scaling are inferred from the dataset, we do not view this as a problem. The third observation we discuss here is that the brightness tends to decrease for movies take from the same coverslip. This is most likely due to some cross-bleaching from imaging neighboring positions. Since the effect is rather small, we will ignore it in the following analysis.

**Figure S11.**
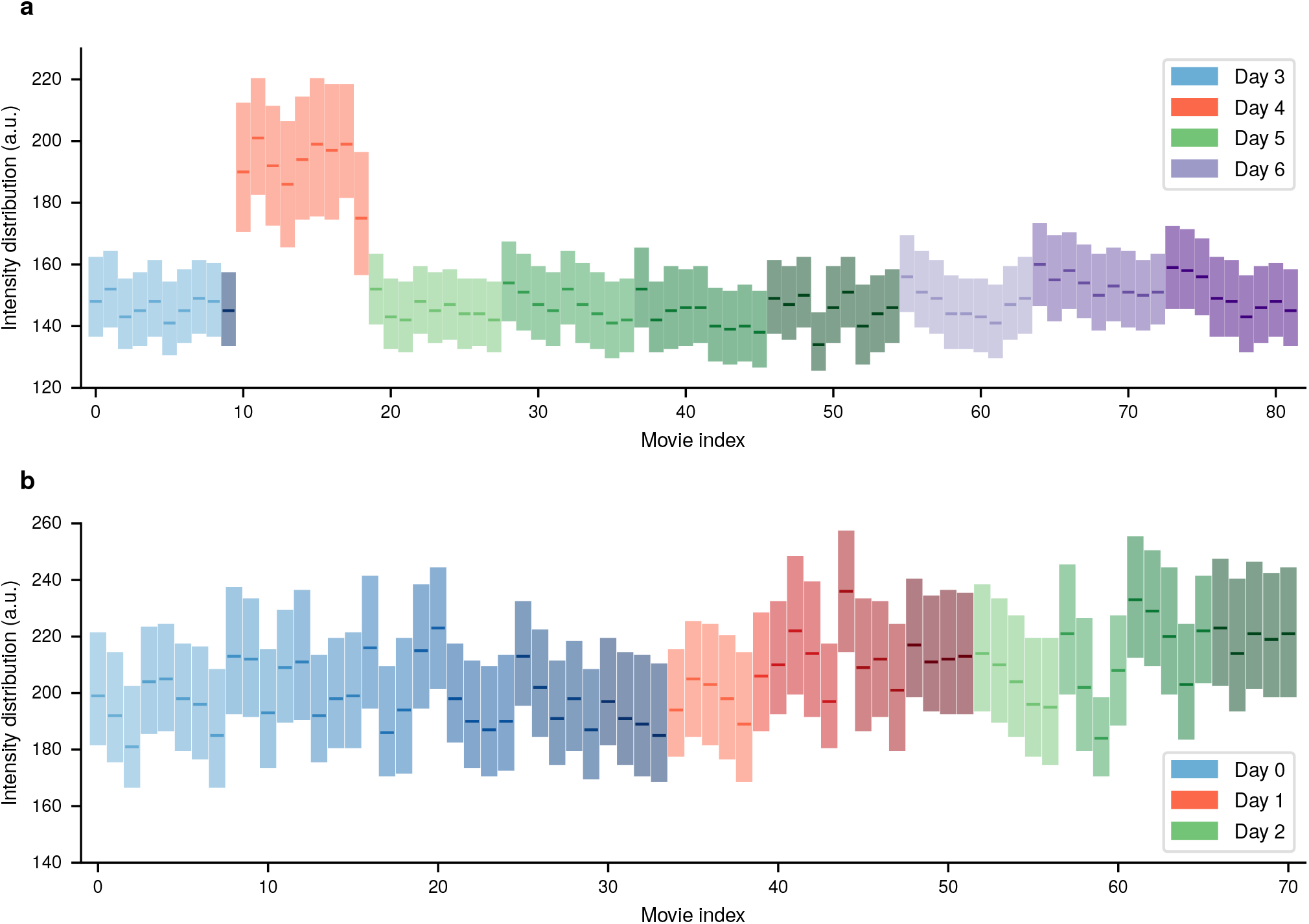
Pixel intensity distributions for the first time frame of every movie in form of a box plot shown for the 3 s dataset (**a**) and the 12 s dataset (**b**). Lower and upper border of the rectangles represent 25th and 75th percentile, respectively. The bold line within the box indicates the median. Every color family represents one experiment day, while the shading withing a color family indicates movies taken from the same coverslip

#### S7.2 Spot Brightness

We now take a first look at the spot brightness. Similarly as an the previous section, we consider only the first frame of every movie to avoid effects of bleaching. For every movie, the spot intensities of all active cells in the first frame were pooled. The corresponding distributions are shown in the form of boxplots for both 3s and 12s dataset in Fig. S12.

Especially in the 3s dataset one can observe a clear dependence of spot brightness on the time since induction. Since the movies taken from the same coverslip are ordered sequentially, they correspond to spot measurements every 3 minutes or every 6 minutes for 3 s dataset and 12 s dataset, respectively. The observed sharp increase followed by a longer decrease is well in accordance with measurements from RT-qPCR. In addition, the results from different coverslips are quite similar and show less variation than the total intensity distributions in Fig. S11. The reason for this is that spot intensities are measured relative to local nuclear background such that additive noise affecting the total brightness of images is filtered out by the spot tracking. The one exception from this are the movies from day 4, which we have already decided to treat as outliers due to their much higher overall brightness.

The dependence on time since induction is also visible in the 12 s dataset albeit less clear than for the 12 s dataset. One reason for this is that there are fewer measurements per coverslip due to the longer duration of the individual movies. In addition, some series of the movies are shifted by 3 minutes leading to a less regular appearance.

Finally, it seems that the spots in the 12 s dataset are overall brighter than in the 3 s dataset. Two possible explanations come to mind. The first possibility is that the cell cultures differ in activity which is possible due the time between the collection of the datasets. Alternatively, multiplicative noise affecting the total intensity distribution could explain the differences as it would not be filtered out by the spot tracking. In practice, it could also be a combination of both factors. We deal with this by estimating the observation model (cf. Sec. S9) parameters individually for every dataset. Any multiplicative noise will be captured by the estimated GFP scaling factor such that remaining differences can be attributed to Biological causes.

**Figure S12.**
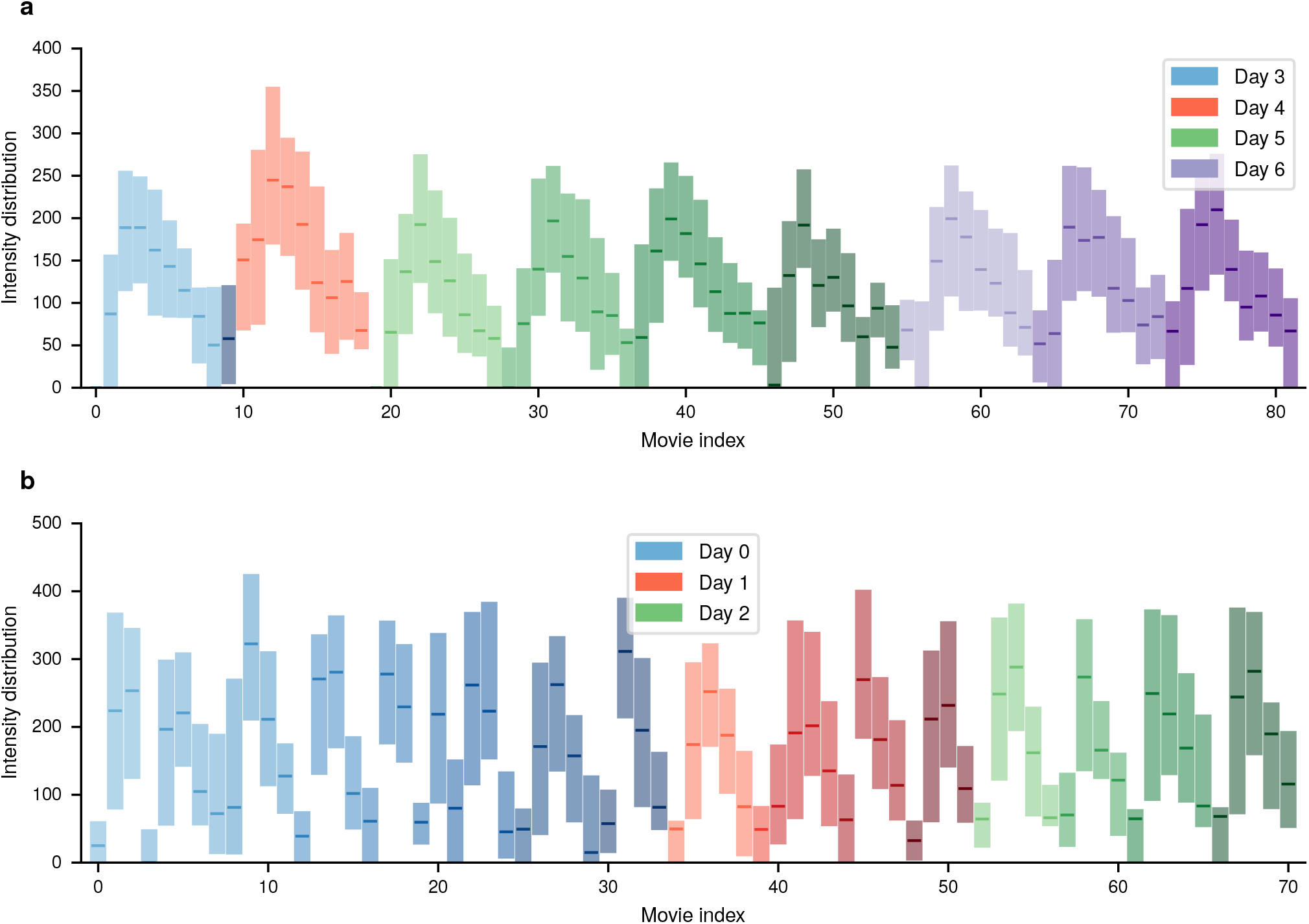
Pixel intensity distributions for the first time frame of every movie in form of a box plot shown for the 3 s dataset (**a**) and the 12 s dataset (**b**). Lower and upper border of the rectangles represent 25th and 75th percentile, respectively. The bold line within the box ndicates the median. Every color family represents one experiment day, while the shading withing a color family indicates movies taken from the same coverslip.

**Table S14.**
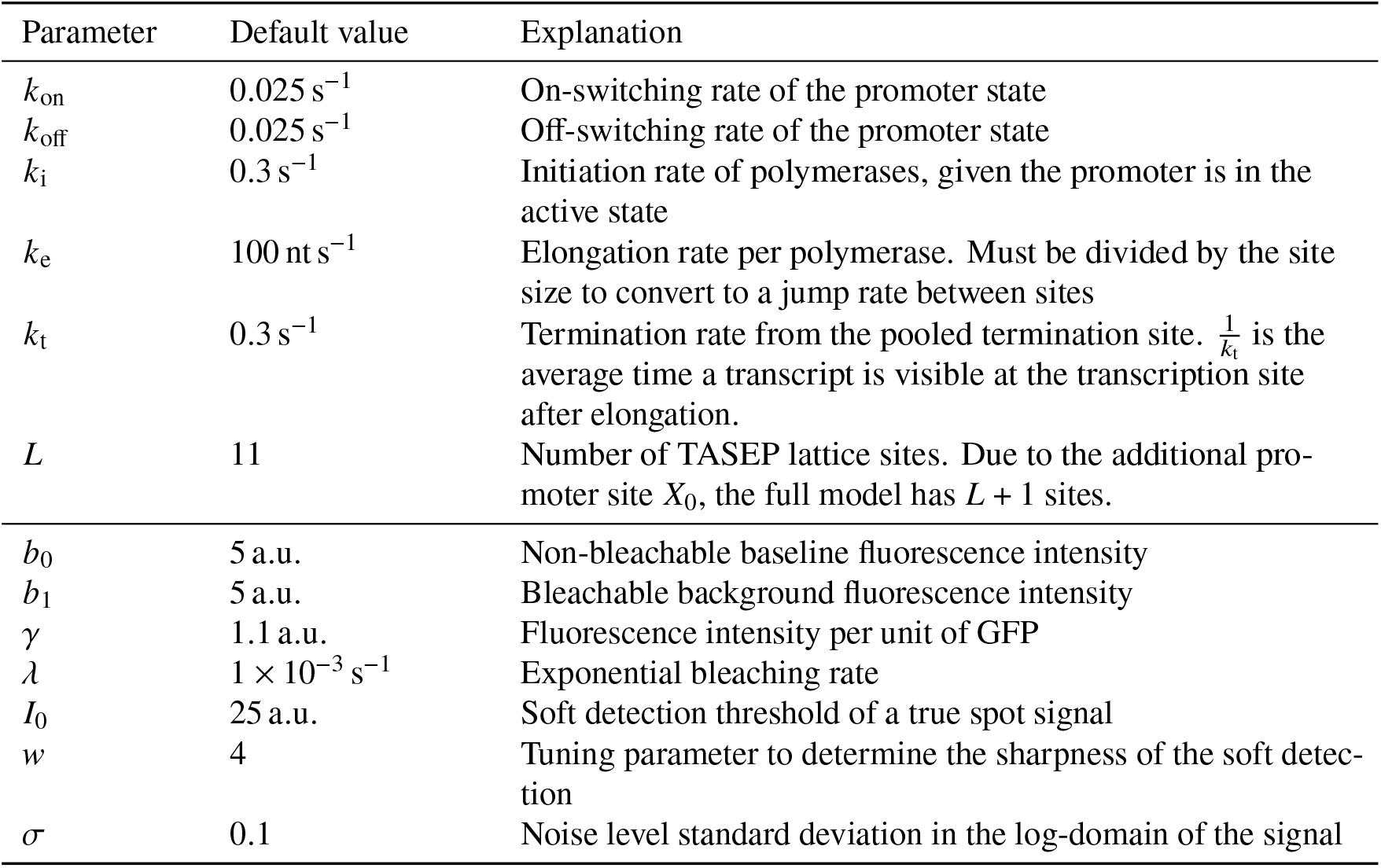
Overview of the model parameters

#### S7.3 Investigating time-dependant activity

The presence of non-stationary transcription dynamics can be already seen from the movie histograms in Fig. S12. We will now take a closer look by inspecting the average spot intensities pooled over all movies that started with the same time delay with respect to induction. As before, we will only consider the first time frame of each movie to avoid the effects of bleaching. The first row in Fig. S13 shows the average spot intensity over the cycle. For both 3 s dataset (left) and 12 s dataset (right) wie see a sharp rise of the intensity in the early part of the observed interval followed by a slower decay. While the general shape of both curves is similar, the spots in the 12s dataset are generally brighter as we have seen from the per-movie distributions (cf. Fig. S12). The second row of Fig. S13 shows the number of responding cells divided by the total number of cells per time window. To assign cells to the class of responders or non-responders, we used a standard Gaussian mixture classifier on the spot intensity distributions of the datasets. In order to avoid bias by the overall different brightness, we applied this classifier individually on each dataset. The resulting responder ratios of both datasets agree well.

**Figure S13.**
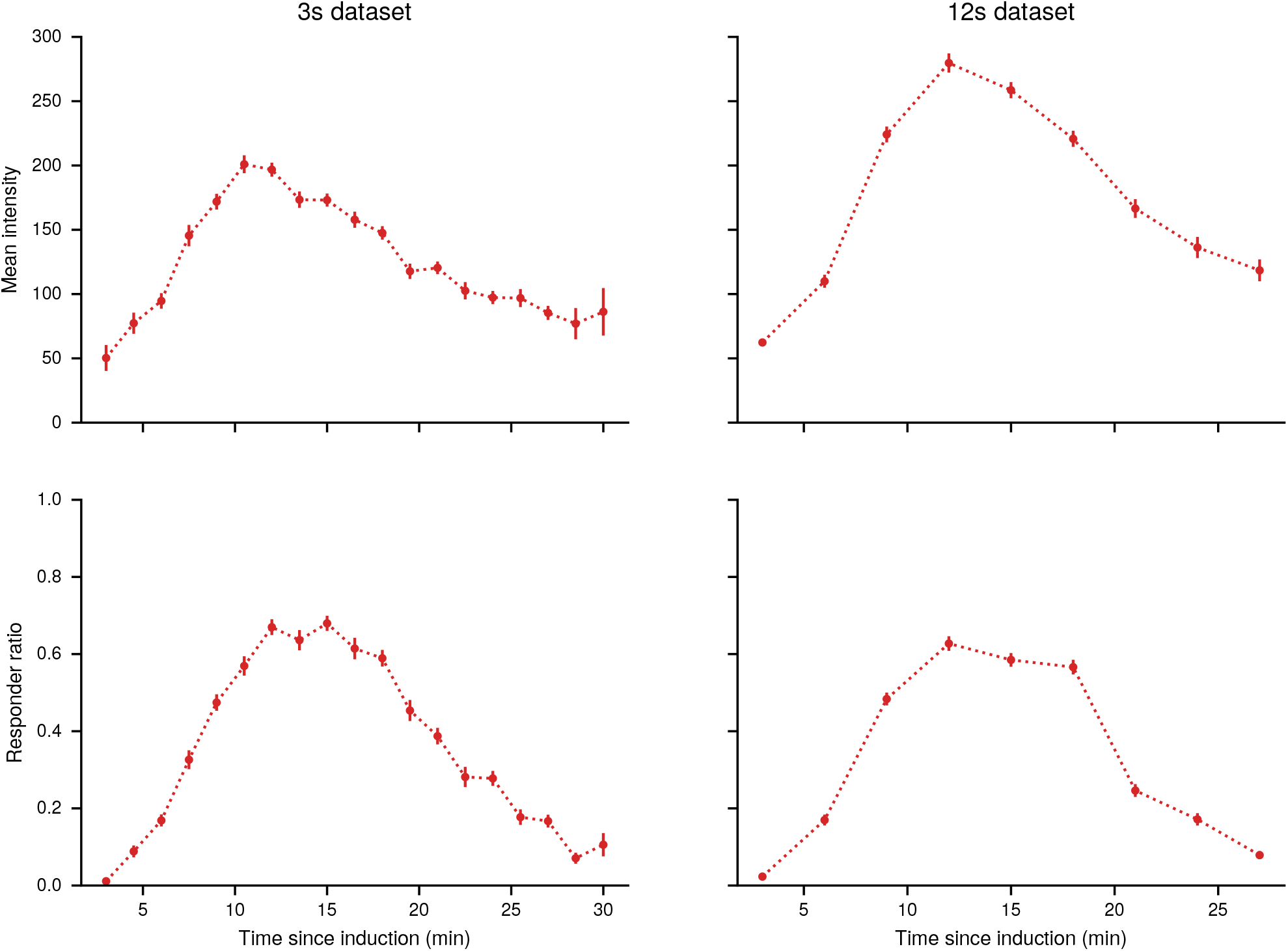
Dependence of summary statistics on the time since induction. The first row shows the mean of the spot intensities pooled over all videos with the same time delay since induction. The second row shows the number of responders divided by the total number of cells per time window. Error bars indicate standard error. Results for the 3 s dataset are also shown in the main text (Fig. 1f, g)

### S8 Model

#### S8.1 Kinetic Transcription Model

As described in the main text, the model is an example of a Markov jump process that satisfies the master equation (1). The transition function *Q* defines the probability of an event to happen in an infinitesimal interval *h*

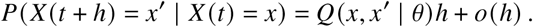

The transition function is fully specified by the vector of parameters *θ* = (*k*_on_, *k*_off_, *k*_i_, *k*_e_, *k*_t_)^⊤^ and the conditions under which transitions can occur, leading to

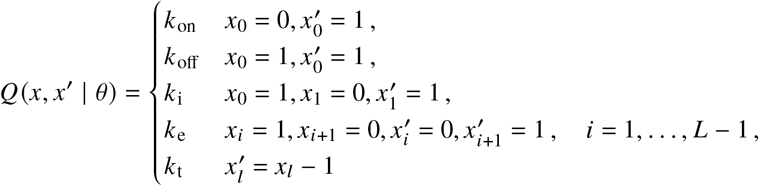

For parameter inference, we need to evaluate the system for many different parameter configurations. It is therefore convenient to represent the transition functions as

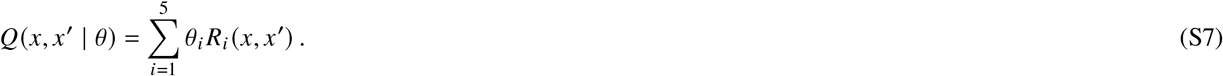

The operators *R*_*i*_ are independent of the parameters. By enumerating the states of the system, the probability *p* (*x, t*) can be represented by a vector **p**(*t*) and the transition function *Q* becomes a sparse matrix **Q**. With this, the master equation becomes

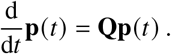

A formal solution of this system is given by the matrix exponential

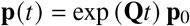

with initial distribution **p**_0_. For sparse **Q**, this can be efficiently solved for fairly large state spaces by the Krylov subspace approximation for matrix exponentials^41, 42^.

#### S8.2 Observation Model

The kinetic model discussed above is a continuous time model. In practice, one cannot observe such a systems in continuous time but rather at discrete sample times *t*_1_, …, *t*_*n*_. In addition, we do not observe the process *X*(*t*_*k*_) directly. Here, our measurement *Y*(*t*_*k*_) is provided by the total intensity of the fluorescence spots as measured by the tracking algorithm. In the following, we construct a model that relates *X* and *Y*. First, note that as the polymerase traverses the gene, an additional stem loop is added for every site until at some point the maximum number of stem loops is acquired. For the remaining part of the transcription process, the number of stem loops stays constant. After termination, the mRNA is released and rapidly diffuses away from transcription site. The corresponding spot is thus no longer visible and we observe a sharp drop in intensity. Hence, if *a* ∈ ℕ^*L*+1^ encodes the number of stem loops associated with the sites of the gene, the variable

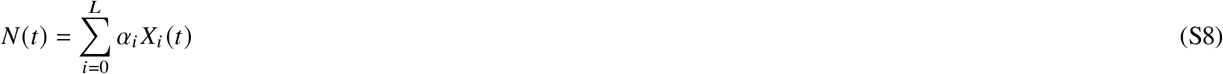

corresponds to the number of visible stem loops at time *t*. Assuming that stem loops are occupied by GFP fast compared to the elongation speed, the total spot signal can be described as

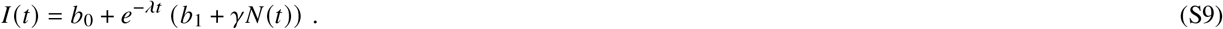

Here, *λ* is the bleaching factor and *b*_0_, *b*_1_ correspond to baseline background and a bleachable part of the background respectively. The factor *γ* encodes the intensity contribution per GFP molecule. During image acquisition and intensity estimation, the signal is corrupted by various forms of noise such as z-diffusion of the transcription site, photon counting noise on the camera chip, variations in the media, mismatch of the point-spread function with the Gaussian approximation, irregular background illumination, etc. We subsume all these effects into a single multiplicative noise variable leading to the

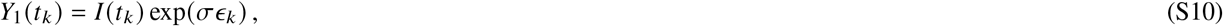

where *ϵ*_*k*_ are independent and standard normally distributed. While (S10) is a reasonable approximation for larger signals, it is not suitable for very small signals. The reason for this is that at low intensity, due to fundamental limitations of the spot detection, there is an increased probability that a spot is missed or that a local fluctuation is confused with a signal. To take account of this effect, we introduce the random background signal

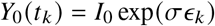

and an additional unobserved random variable *Z* (*t*_*k*_) ∈ {0, 1} with

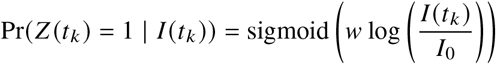

where sigmoid(*x*) = (1 + exp(−*x*))^−1^ and *w* is a response parameter. The final observation model is then given by

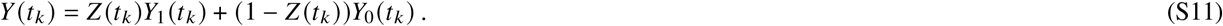

This can be understood as a soft threshold for spot detection. When the true spot intensity *I* is larger than *I*_0_, *Z* will likely be one and we measure the signal *Y*_1_ with high probability. When *I* is smaller than *I*_0_, *Z* is likely zero and we observe the spurious signal *Y*_0_ with high probability. The likelihood corresponding to (S11) is that of a mixture of two lognormal distributions with parameters log(*I*) and log(*I*_0_). An overview of all parameters involved in kinetic and observation model is given in Table S14.

#### S8.3 Elongation times

Many models for transcription assume independent movement of individual polymerases. This leads to a simple relation between elongation rate and elongation time as 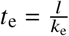 where *l* is the size of the translated region. Due to possible interactions between polymerases, this relation is not valid in the TASEP model. As shown in Fig. S14, the TASEP model requires a higher value compared to an independence model to produce the same expected elongation time. This difference becomes smaller for higher elongation rates, since a fast movement of individual polymerases decreases the probability of interaction. As an example, consider the dotted black line in Fig. S14 corresponding to an elongation time of 12 s. To produce such an elongation time, the independent model requires an elongation rate of 100 nt s^−1^, while the TASEP model requires and elongation rate of roughly 120 nt s^−1^.

**Figure S14.**
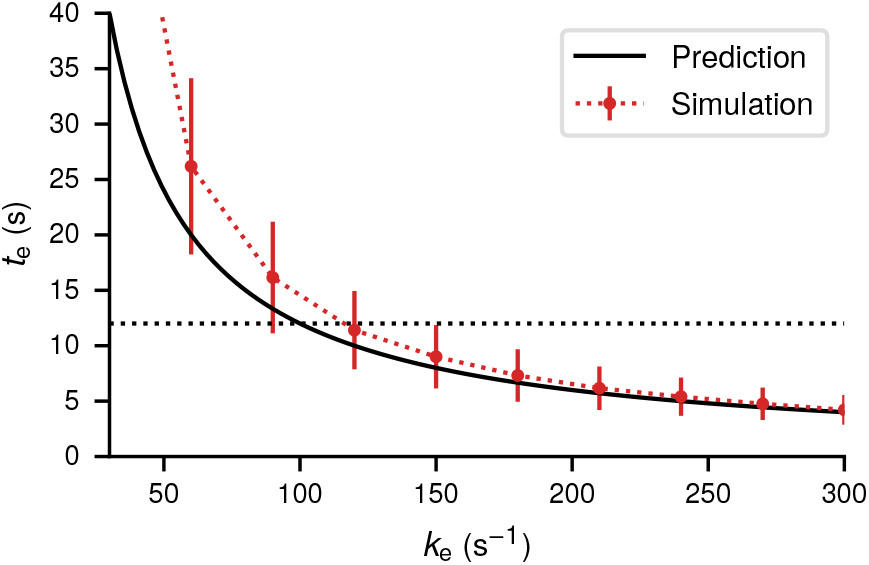
Expected elongation times of a gene template with size *l* = 1200 nt for different values of the elongation rate *k*_e_. The thick black curve indicates the value obtained from independent polymerase movement by the relation 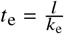. Red dots show the expected elongation time from simulations of a TASEP model with a site size of 120 nt. Error bars indicate the standard deviation of the distribution. Initiation rate *k*_i_ and termination rate *k*_t_ are fixed and chosen such that typically multiple polymerases are on the template and no traffic jams are caused by the termination site.

#### S8.4 Coarse-graining

The most natural quantization of the gene into sites would be to associate every site with a single nucleotide. This would, however, lead to more than 1000 sites and since the state space of the model scales as 2^*L*^ would make inference intractable. In addition, we only observe the system by one-dimensional summary statistic every few seconds such that most of the detailed dynamics are not captured. It is thus necessary to combine several nucleotides into a single site. A good candidate for such a coarse-graining is the DNA footprint of a stem loop, which is roughly 60 nt, as the appearance of a stem loop is the most fine-grained observable. For a DNA template consisting of 1200 nt, this would lead to *L* = 22 sites. While the master equation this model is still tractable, it requires significant computational effort such that only a small number of cells can be handled this way. As argued in the main text, it is important to pool many traces in order to overcome structural identifiability limitations of the system. As a compromise between tractability and resolution, we choose a partition size of 120 nt corresponding to two stem loops leading to a system of *L* = 12 sites.

The coarse-graining changes the waiting time distribution between appearance of two stem loops. If one starts from a fine-grained model where every site corresponds to a single nucleotide, the waiting time for jumps of size 120 bp is much more peaked compared to the exponential distribution. To investigate the robustness of the inference against this kind of mismatch, we simulated 100 trajectories from a fine-grained model described in^23^ Supp. M. The model is similar to the TASEP model we use for inference, but uses one site for every nucleotide. In addition, RNAP molecules have a footprint of 40 sites and individual stems loops appear every 60 sites with 14 stem loops in total. The number of sites was set to 1200 roughly corresponding to the gene investigates experimentally. The elongation rate was set to *k*_elong_ = 100 nt s^−1^, for all other parameters we used the standard values of the coarse-grained model. We generated 100 trajectories with 3 s time-lapse and performed pooled inference. For simplicity, we assume a constitutive promoter an drop the switching site. The results are presented in Fig. S15a. They indicate that while initiation and termination rate are inferred quite accurately, the estimated elongation rate is, however, significantly larger than the ground truth used to generate data in the fine-grained model. This can be explained by the exclusion property of the TASEP process. In the coarse-grained model, collisions will happen more frequently on average since the space is limited. In order to achieve a similar total elongation time, the rate has to be increased to account for the possible blocking. To verify this, we generated trajectories from fine-grained model and coarse-grained model and extracted the distribution of total transcription times (Fig. S15b). The mean elongation time predicted by the learned coarse-grained model is quite close to the mean elongation time of the fine-grained model. This demonstrates that even though there is a mismatch, the model learns physically meaningful quantities. However, we stress that this is merely an example to illustrate model mismatch. In practice, the distribution of elongation times predicted by the fine-grained model is likely too narrow as it ignores effects such as pausing, reverse steps, chromatin remodelling, etc. Thus, a coarse-grained model with wider waiting time distribution may be a better approximation to the real data, as long as these effects are not modeled explicitly.

#### S8.5 Overview of the model parameters

**Figure S15.**
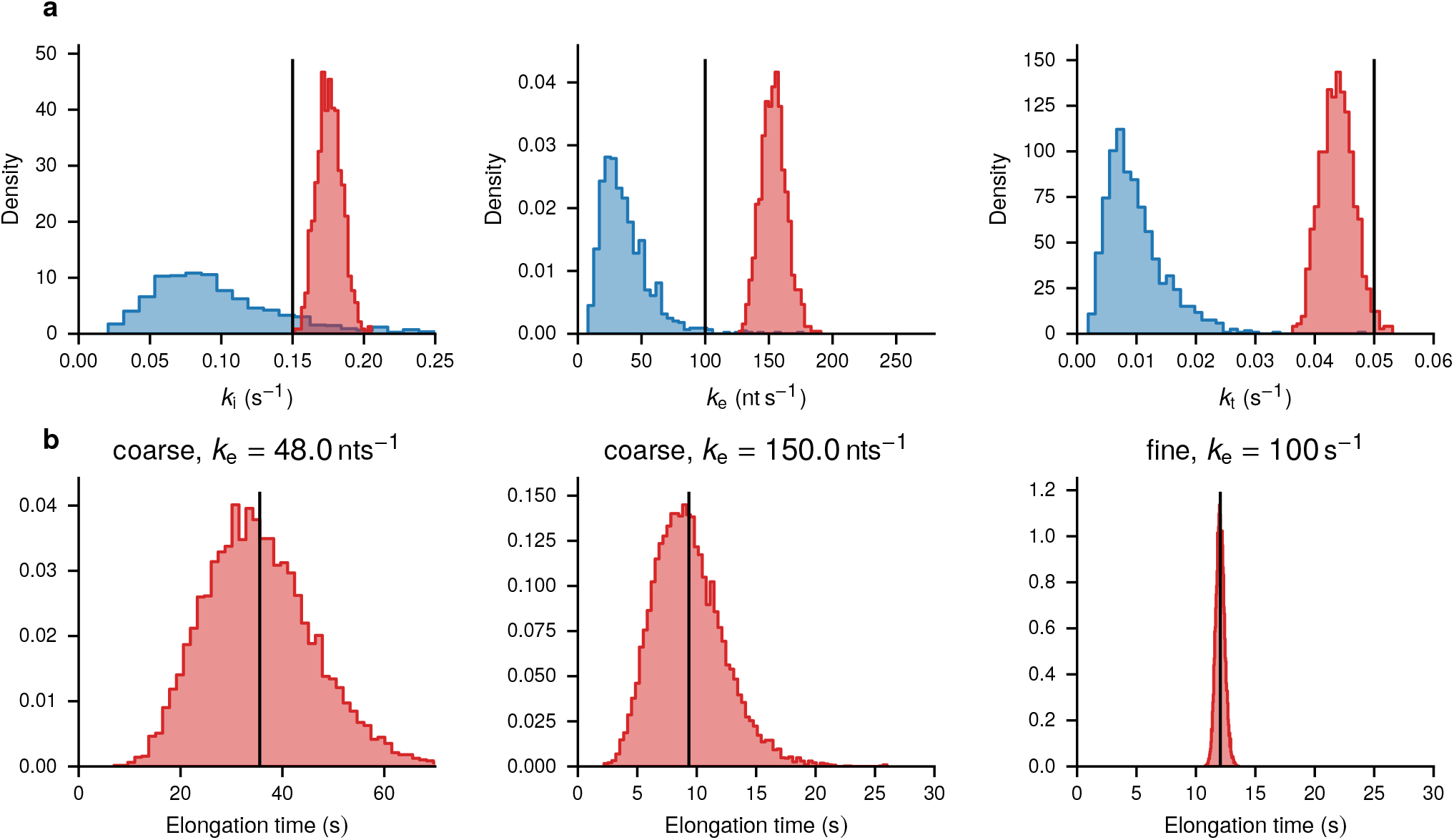
**a** Pooled posterior inference of a non-switching model on traces generated by the fine-grained TASEP model with a time laps of 3 s. Observation parameters are not shown but were also estimated during inference. The rows show histogram approximations of the prior distribution (blue) and the posterior distribution (red) for the model parameters. Black lines indicate the parameter value used to generate the data. **b** Distribution of total elongation times obtained from forward simulation of the coarse-grained model (left and middle) and the fine-grained model (right). The elongation right of the left plot is a typical value of the prior distribution, the elongation rate of the middle plot is a typical value of the posterior distribution. Black lines indicate the empirical mean.

### S9 Calibration

The observation model as described in Sec. S8.2 contains a number of unknown parameters. In particular, the bleaching rate *λ* and the GFP scaling factor *γ* can have a significant effect on the inference results. It is therefore helpful to obtain independent estimates of these quantities from dedicated control data sets which can be used as prior distributions in the live cell inference procedure. The last subsection deals with the point spread function (PSF) of the optical system. While the PSF is not directly contained in the generative model, it is used to extract the intensity measurements from the images.

#### S9.1 GFP-intensity scaling

For calibration measurements, we activated *CUP1* array in YTK541 with Cu and imaged the cells between 5 and 15 min of the Cu treatment, at the peak of their transcriptional activity to ensure that all the 40 binding sites are occupied with Ace1p-3xGFP fusion, and thus the total number of GFP per CUP1 array is 120. In YTK1231, we measured the brightness of the lacO/LacI-GFP array in telophase or G1 cells to ensure that we observe single non-duplicated array. LacI-GFP was present on multicopy plasmid, and some of the cells in the population were overexpressing LacI-GFP and thus displayed a high nuclear background. Therefore, we excluded the cells with abnormally bright nucleoplasm or arrays. In diploid cells of YTK1268, two peaks are observed in the SPB size distribution and about 5 % of the SPB in the population are significantly larger than average^39, 40^. This happens because the SPB grows over time through the cell cycle. At G2 stage the SPB is split in two. Therefore, we avoided newly split SPB and selected only single SPB in the cells in telophase or in G1, where maximal size of the SPB is expected. As a control we also measured the cells in G2 with two SPB well separated by spindle and observed diminished brightness of those structures, as expected. Also, we selected the sub-population of SPB based on the range of brightness excluding from the measurements the SPB that were abnormally bright. Spot intensities of all structures were measured as for the live cell experiments (see Sec. S6.2) but only for a single time point.

A field view for each of the different strains used for this is shown in Fig. S16a. A first overview of the calibration data as shown in Fig. S16b reveals that the linear approximation assumed in (S9) is reasonable. In addition, the noise increases with expected number of GFP confirming the multiplicative noise model (S10). For a more detailed analysis, note that we have single images rather than videos as calibration data. Thus, with *t* = 0 and (S9) reduces to

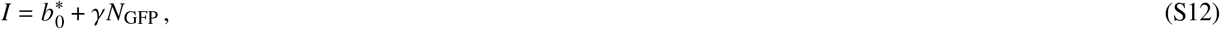

where 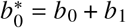 and *N*_GFP_ corresponds to the number of GFP associated with a particular structure. Together with the noise model (S10) and priors for 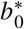 and *γ*, we can perform Bayesian inference by Hamiltonian Monte Carlo. The obtained posterior is shown in Fig. S16c. It is well approximated by a Gamma distributionΓ *α, β* with *α* ≈ 4226.4, *β* = 3822.8. This distribution is used as a prior for the main inference part.

#### S9.2 Bleaching Rate

To obtain an independent estimate of the bleaching rate, we recorded videos with 12 s intervals between observations with strain YTK1231 with *lac*O/LacI-GFP. Since GFP load is fixed for this structure, we expect any systematic changes in brightness over time to be caused by bleaching. This allows for an independent estimate of the bleaching factor. The mean intensity of this data over time is shown in Fig. S17a along with an exponential fit. We observe that the effect of bleaching is not very strong in the time window considered. To calibrate the observation model for Bayesian inference, we use Markov chain Monte Carlo sampling so get a posterior distribution over the bleaching factor that can be used as a prior for the main experiments. We again start from (S9). In contrast to the GFP calibration, time course data is available. However, *b*_1_ and *I* are not distinguishable due to the lack of spot dynamics leading to te reduced model

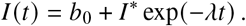

The corresponding Bayesian posterior for the parameter *λ* is shown in Fig. S17b leading to an estimate of *λ* = 0.007 s^−1^. For the 3 s interval movies, we do not use separate calibration data. Instead, we observe that light exposure is the main factor determining the bleaching rate. A four-fold increase in imaging frequency should therefore correspond to a four-fold increase in the bleaching rate. As this calibration only serves to construct a prior distribution, this estimate is deemed sufficient.

#### S9.3 Point Spread Function

We use a model-based approach to evaluate the spot intensity where the spot is described as a local background *b* plus a point source with intensity at (*x*_0_, *y*_0_). In the image space, this corresponds to an intensity profile of the form

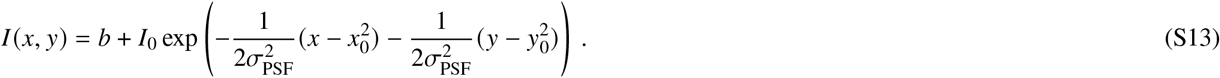

**Figure S16.**
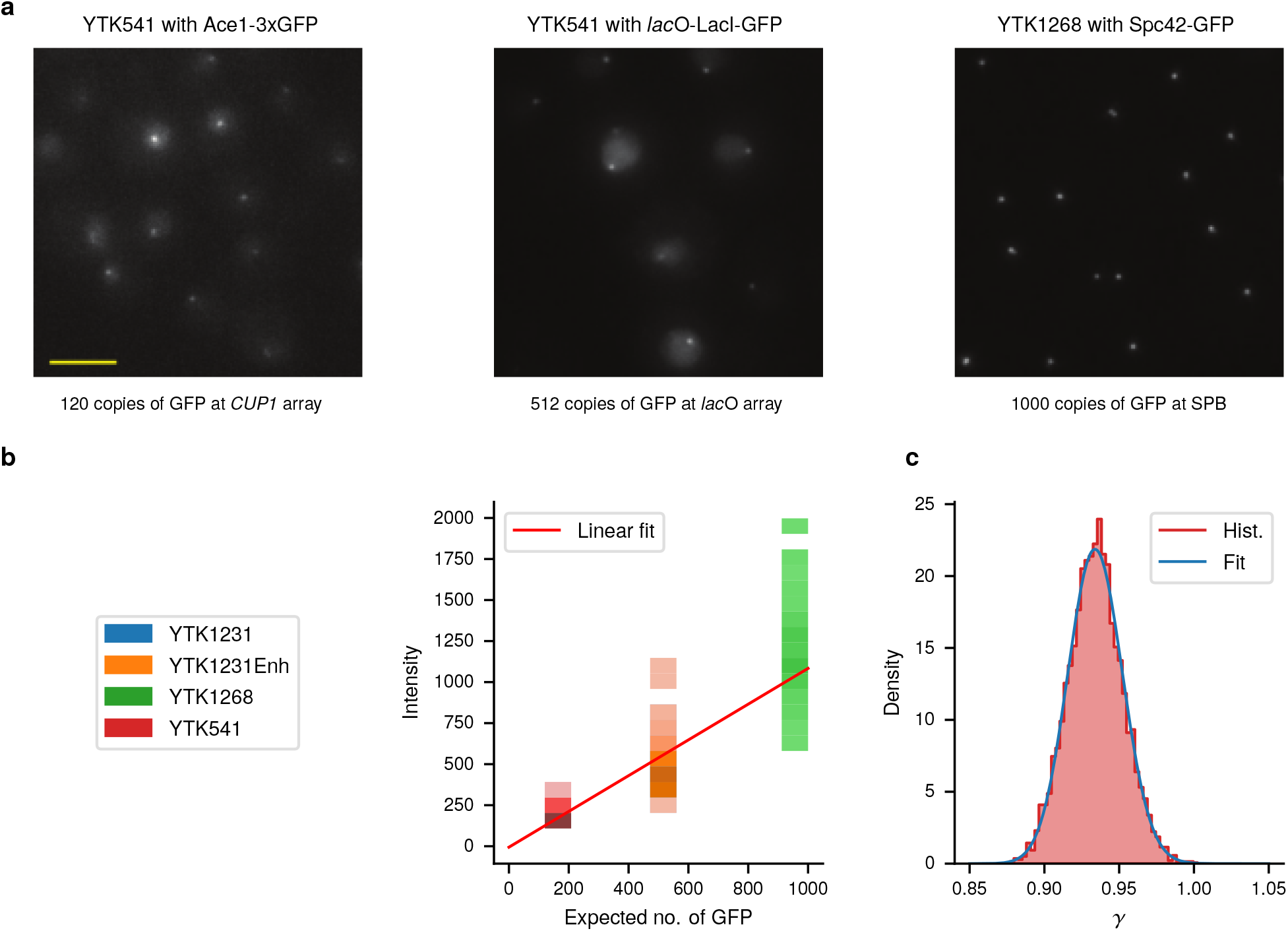
**a** Representative field views of cells with known numbers of GFP molecules per locus. **b** Color-coded histogram representations of the intensity distributions for the different strains. Darker colors indicate areas of higher density. The red line shows a least-squares fit of (S12) to the distribution. **c** Histogram of the posterior samples of *p* (*γ* | *I*_1_, …, *I*_*n*_) based on intensity estimates from *n* = 509 images (red). The blue curve corresponds to a gamma distribution fitted to the posterior samples.

The exponential term in (S13) corresponds to the point spread function with σ_*PSF*_ and peak intensity *I*_0_ describing the shape of the point in the image space. The parameter σ_*PSF*_ depends on the properties of the optical system. While there are approximate formulas, these estimates are often not very accurate in systems with high aperture. We therefore infer σ_*PSF*_ directly from a calibration dataset.

In fluorescence microscopy, the image is usually acquired by a CCD camera that discretizes the image space into a pixel grid. An example of a spot image from the strain *YTK1268* with spindle pole body is shown in Fig. S16a. The predicted intensity *p* (*i, j*) of the pixel at position *i, j* is given by

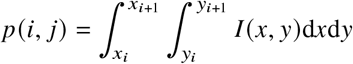

with pixel boundaries *x*_*i*_, *x*_*i*+1_, *y*_*i*_, *y*_*i*+1_. We assume now that the pixels have square shape and choose a coordinate system such that the pixel area is one. The intensity can then be expressed by

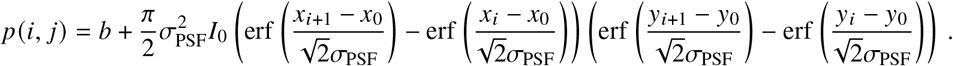

Finally, in the measurement process, the intensity is corrupted by multiplicative noise. The measured intensity 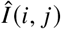 is thus given by

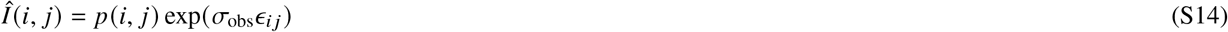

**Figure S17.**
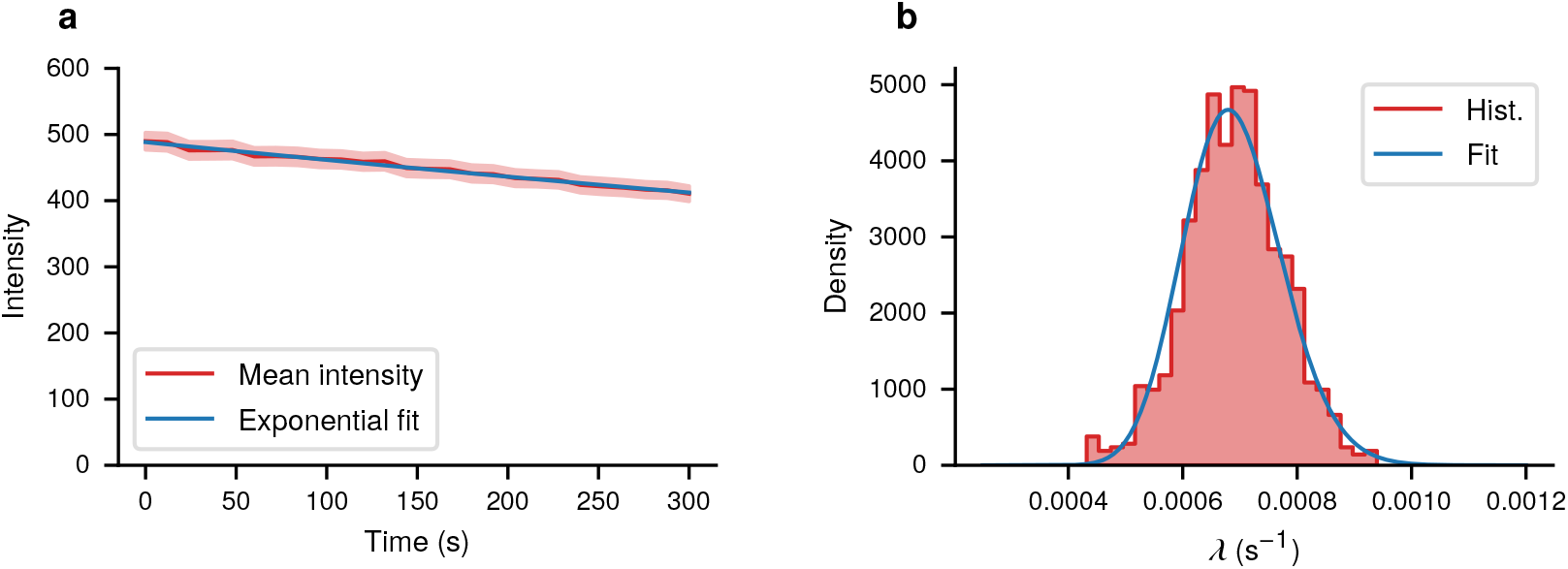
**a** Mean spot intensity of *n* = 252 cells of YTK1231 shown over time (red). The shaded region indicates the standard error. The blue line is a least squares fit of an exponential function. **b** Histogram of the posterior samples of *p* (*λ* | *I*_1_, …, *I*_*n*)_ based on intensity estimates from *n* = 252 videos (red). The blue curve corresponds to a gamma distribution fitted to the posterior samples.

where σ_obs_ is the noise level and *ϵ*_*ij*_ are i.i.d. standard normal random variables. By choosing priors for *b, I*_0_, *x*_0_, *y*_0_ and σ_PSF_, we have constructed a generative probabilistic model for spot images. We can then compute the posterior 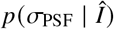 given an image 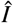. Generalization to multiple image samples is straightforward. A graphical model representation is shown in Fig. S18b. Using a non-informative prior, we run Hamiltonian Monte Carlo for inference. As shown in Fig. S18c, the resulting posterior is quite concentrated which justifies using a point estimate of σ_PSF_ ≈ 1.926 during spot intensity estimation.

**Figure S18.**
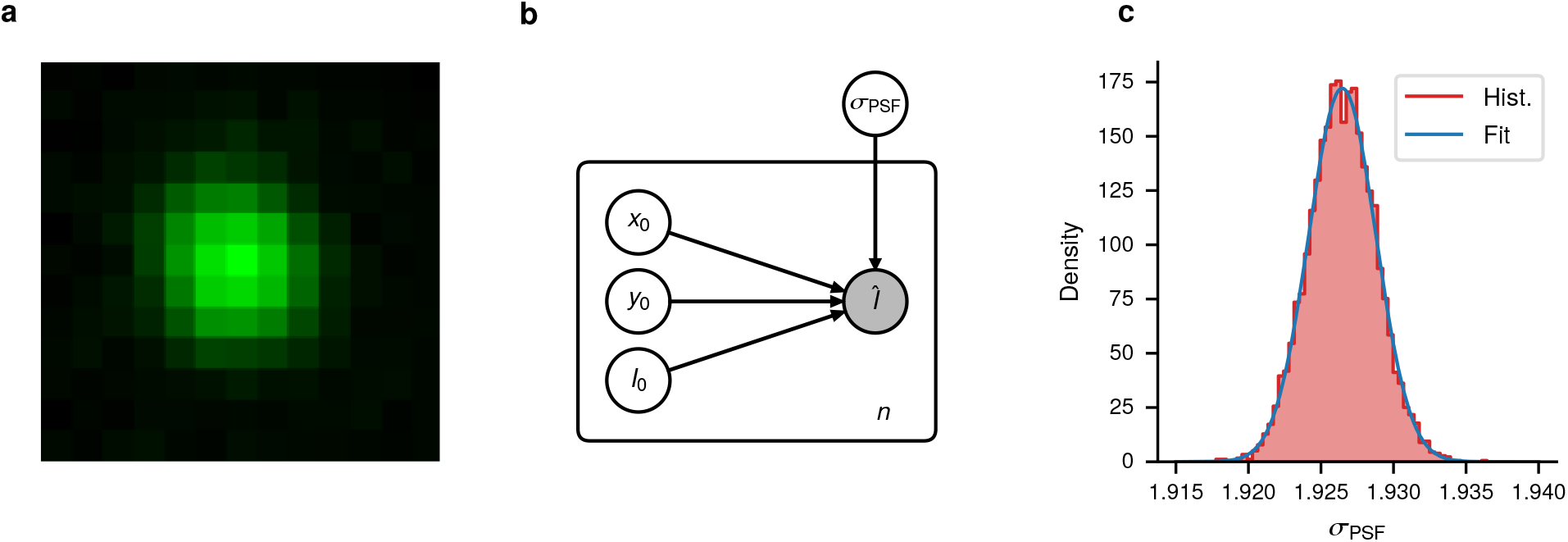
**a** Example of a spot image obtained from the strain YTK1268 with a spindle pole body. **b** Probabilistic graphical model for inferring the point spread function parameter σ_PSF_ jointly from *n* images. **c** Histogram of the posterior samples of 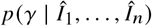 based on *n* = 509 images. The red line corresponds to a log-normal distribution fitted to the posterior samples.

### S10 Inference

#### S10.1 Fully Bayesian inference

Given an observed trace *y* = (*y*_1_, …, *y*_*n*_) we are interested in estimating the latent state *x*_[0,*T*]_, the model parameters *θ* and the observation parameters *ω*. In a Bayesian approach, this corresponds to computing the joint posterior *p* (*θ, ω, x*_[0,*T*] |_ *y*_1_, …, *y*_*n*_). Targeting this posterior directly with Monte Carlo methods is difficult. Exploiting the relation

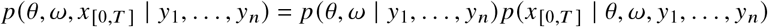

full posterior can be split joint into two parts: The marginal parameter posterior

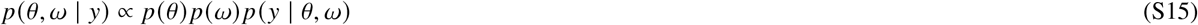

and the conditional state posterior *p*(*x*_[0,*T*]_ | *θ, ω, y*_1_, …, *y*_*n*_). Intuitively, the conditional state posterior characterizes the distribution of paths of the underlying continuous-time process that have most likely created the observed trace *y*. One can generate samples from such a conditional Markov process using the backward filtering forward sampling approach. This is based on the observation that for a Markov jump process with generator *Q*(*x, x*′) the smoothing process is equivalent to a modified process with time-dependent generator^46^

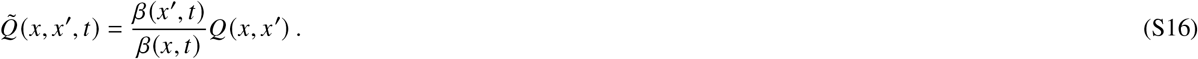

Here, *β*(*x, t*) = *p* (*y*_*k*_, …, *k*_*n*_ | *X* (*t*) = *x*) with *k* = min{*i* : *t*_*i*_ > *t*} is the probability density of future observations given the current state. Now *β*(*x, t*) satisfies a backward equation

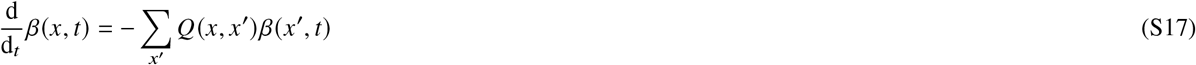

with reset conditions

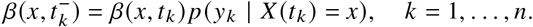

Sampling proceeds by solving (S17) backward in time and then using (S16) within a time-dependent version of the stochastic simulation algorithm^44^. The marginal parameter posterior (S15) is a continuous, finite-dimensional distribution and can, in principle, be tackled by a number of MCMC algorithms. A computational challenge, however, is that sampling from (S15) using MCMC requires evaluation of the marginal data likelihood *p*(*y* | *θ, ω*). The marginal likelihood can be evaluated using filtering theory for Markov processes. Consider the filter distribution *α* (*x, t*) = Pr(*x* (*t*) = *x* | *y*_1_, …, *y*_*k*_), *k* = max {*i* : *t*_*i*_ ≤ *t*} describing the best estimate of the current state of the system given all past observations. From general recursive filtering theory, we obtain that *α*(*x, t*) obeys the master equation in between the observation times, and at the observation times satisfies the update conditions

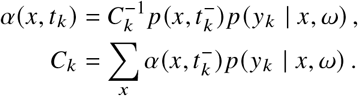

Furthermore

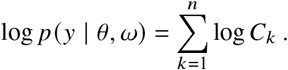

A derivation of the filter solution of the marginal likelihood is discussed in Sec. S10.2. To evaluate the gradient of the marginal likelihood we use the adjoin method and obtain

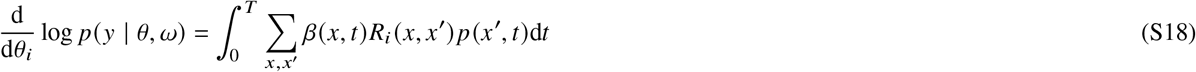

where *β*(*x, t*) is a backward filter as in (S17) but with modified reset conditions

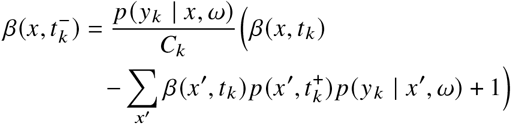

and *R*_*i*_ is the parameter independent part of the generator (cf. (S7)). A derivation of the modified backward filter is provided in Sec. S10.3. Having access to log *p*(*y* | *θ, ω*) and ∇_*θ*_ log *p*(*y* | *θ, ω*) allows to use efficient Monte Carlo methods such as Hamiltonian Monte Carlo47. In addition, the model can be integrated with probabilistic programming^48^ facilitating easy reuse and modification of our approach.

In practice, we use the marginal approach for parameter inference. If latent state inference is required, the full posterior can be reconstructed by resampling *θ, ω* from the parameter posterior and generate a trajectory from the smoothing process. These samples can be used to investigate arbitrary summary statistics of the latent process given the data, for example initiation times and local polymerase speed.

The discussion above refers to inference from a single trace. In case of a dataset 𝒟 = {*y* ^(1)^, …, *y* ^(*m*)^} of *m* traces, a joint analysis with a hierarchical model is possible. To generalize the above to such a scenario, we view each trace 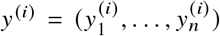 as one single random variable (cf. main text, Fig 2e,f). As *p*(*y*^(*i*)^ | *θ, ω*) is a finite dimensional distribution, constructing hierarchical models for a collection of traces follows the usual rules of probabilistic modeling and Bayesian inference^35^. Hierarchical modeling is discussed further in Sec. S10.4. The joint analysis requires evaluating forward and backward filters for every trace in the dataset in every step of MCMC. Therefore, full MCMC becomes infeasible for more then a few hundred traces. A viable alternative in this case is stochastic variational inference with mini-batching, that allows to efficiently compute the best approximation of the posterior within a parametric family of distributions^28^.

#### S10.2 Marginal likelihood by stochastic filtering

Formally, the marginal data likelihood is given by

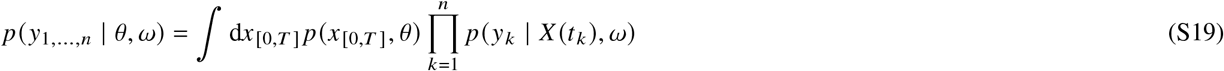

where the integral is over all sample paths of stochastic process *X t*. Using the Markov property, the path integral (S19) reduces to the finite dimensional sum

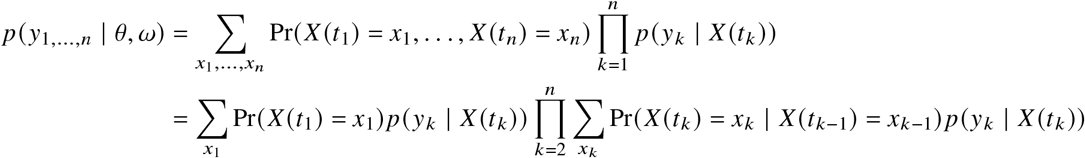

which is analogous to the likelihood of a hidden Markov model^43^. The above expression can be computed recursively by stochastic filtering. To see this, we introduce the forward filter

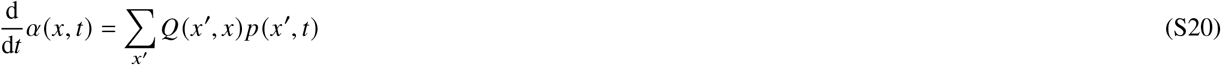

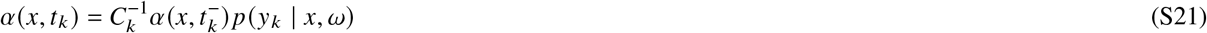

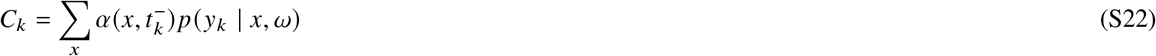

For the normalization constant *C*_*k*_ we have

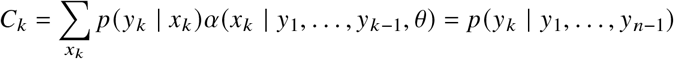

from which follows

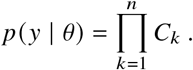

#### S10.3 Gradient of the marginal likelihood

To calculate the gradient ∇_*θ*_ *p*(*y* | *θ, ω*), we observe that the likelihood depends on the parameters only implicitly through the prediction step in (S20). More specifically, we require the derivative of

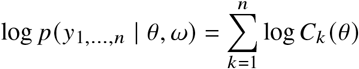

subject to

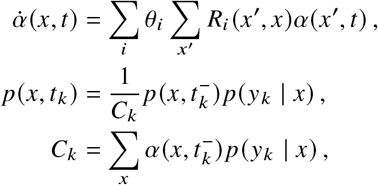

where we have used the parameter form of the transition function (S7) in the master equation within the constraint. To compute the gradient of such a constrained function, one can use variational calculus. First, the constrained problem is transformed to an augmented unconstrained functional known as the Lagrangian

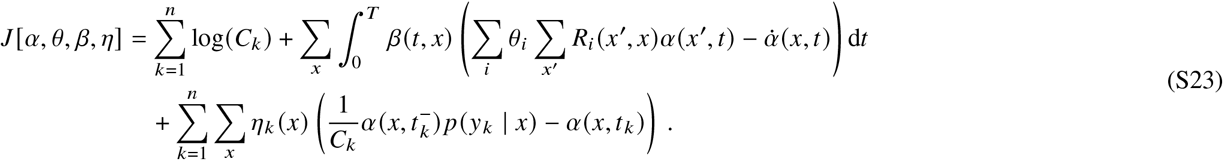

The gradient of the original function can be computed from the stationary conditions of the Lagrangian. In particular

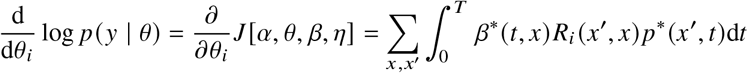

where *α* is the solution of (S20) for given *θ* and *β, η* satisfy the stationarity condition

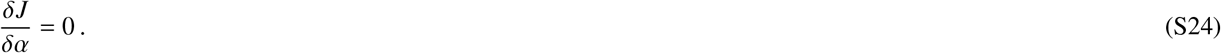

To calculate the functional derivative, we choose a suitable perturbation *δα* and linearize (S23) around *α*. After reorganizing terms, we get

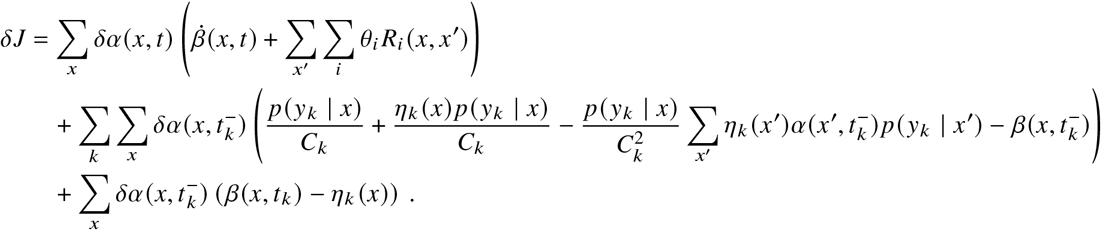

Since *δα* is arbitrary, the terms in brackets must vanish to satisfy (S24). This leads to the adjoint state equation

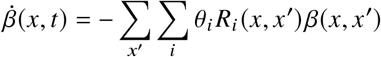

along with the jump conditions

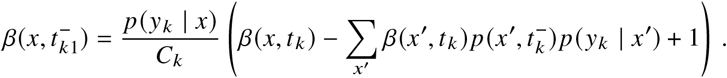

For applying HMC, we also require the gradient ∇ _ω_ *p*(*y* | *θ*, ω). This gradient is straightforward to compute since *ω* only affects the observation likelihood

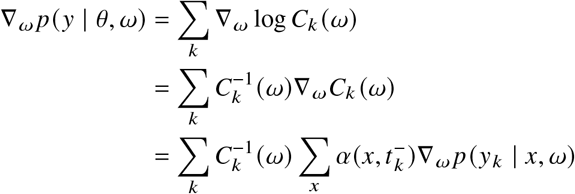

which corresponds to a weighted average of the gradients of the observation likelihood.

#### S10.4 Hierarchical modelling

Consider a collection of traces 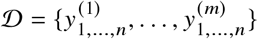. As discussed in *Methods — Bayesian inference*, we can consider the discretely observed trace *y*_1,…,*n*_ as a single random variable generated from a distribution *p*(*y*_1,…,*n*_ | *θ, ω*). This allows to apply the usual rules for probabilistic graphical models and Bayesian inference to construct hierarchical models. For convenience, we provide a few more explicit examples in this section.

##### Pooling of multiple traces

The simples model for a collection of traces 𝒟 is that of identical and independently distributed samples. This means that two parameter vectors *θ, ω* are shared by all the traces. We are interested in the posterior

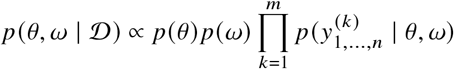

where *p*(*θ*) and *p*(*ω*) are appropriate priors for transcription and observation parameters, respectively, and each term in the product is a marginal likelihood for a single trajectory as in (S19). This model uses identical parameters for all time windows meaning that it allows no cycle dependence and is considered as a baseline. More complex models will be scored against the fully pooled model.

##### Per-window pooling

Consider now traces 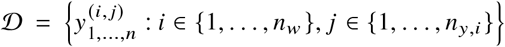 where *n*_*w*_ is the number of windows and *m*_*i*_ is the number of traces in window *i*. For every window, we assume independent parameters *θ* ^(*i*)^, *ω*^(*i*)^. In this case, the inference problem decomposes into individual posteriors for every window

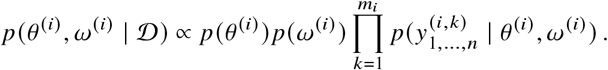

This corresponds to repeating the simple pooling independently for each window. While straightforward to apply, this approach does not exploit shared information between traces of different windows, which can lead to poor accuracy if the number of traces per window is small.

##### Mixed approach

In practice, some of the parameters may depend on the cycle while others are shared between all traces. Throughout, we assume a global observation parameter *ω*. The kinetic transcription parameters *θ* are split into global parameters *θ*_*g*_ and local parameters 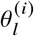 that are allowed to vary for different windows. The corresponding posterior is of the form

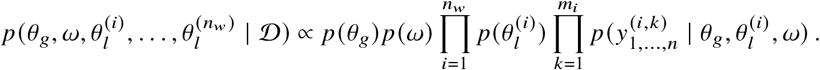

By performing inference with different partitions into local and global parameters and comparing the different models, this provides some insight into which parameters are affected by the cycle.

#### S10.5 Parameter identifiability

The presented model contains 5 kinetic parameters and additional 7 observation parameters. In addition, the latent stochastic process *X*(*t*) is a fairly high-dimensional lattice model (≈ 2 × 10^5^ states in the configuration used for most experiments) that is observed by fluorescence intensity, which is a one-dimensional quantity. This setting raises the question of parameter identifiability, which we investigated based on simulated data. Preliminary numerical experiments indicated that in particular the GFP intensity calibration factor *γ* has a major impact on inference quality. If an uninformative prior is used for *γ*, the system is not practically identifiable with a realistic number of traces. We therefore used calibration measurements to construct a tight prior for *γ* (see Sec. S9.1). However, even with good prior knowledge of *γ*, a single trace is not very informative. Pooling of multiple traces can increase accuracy significantly. This was demonstrated in the main paper using the example of the initiation rate (cf. Fig. 1f). The full results with posteriors for all parameters are given in Fig. S19.

#### S10.6 Computing infrastructure

Numerical experiments using the real data were run on the Hessian High Performance Computer (HHLR) located at TU Darmstadt. A typical run of the variational inference was performed on a single compute node consisting of 96 Intel Xeon Platinum 9242 processors for 24 hours which allowed for roughly 2000 gradient steps. The experiments based on simulated used fewer traces and were thus run on the local cluster of Self-Organizing Systems Lab on a 22 core machine with Intel Core Haswell architecture.

#### S10.7 Custom Software

The core of the code consists of the following parts: A model builder that allows to straightforwardly implement arbitrary CTMC models, a simulation module for data generation, an integrator for solving the master equation based on the Krylov subspace approximation of the matrix exponential^41, 42^, programs for evaluating the data likelihood and the corresponding gradient with respect to the parameters and a simulator for the state posterior. The backend of the code is implemented in C++ and was compiled into a native Python extension using the Pybind11. Matrix operations are implemented using the Eigen library. The code has been parallelized using OpenMP such that it can process multiple traces simultaneously. For inference, our python extension has also been interfaced with the PyTorch package and the probabilistic programming language Pyro^45^. This gives access to state of the art inference algorithms and simplifies implementing hierarchical models. The code is available at XXX.

**Figure S19.**
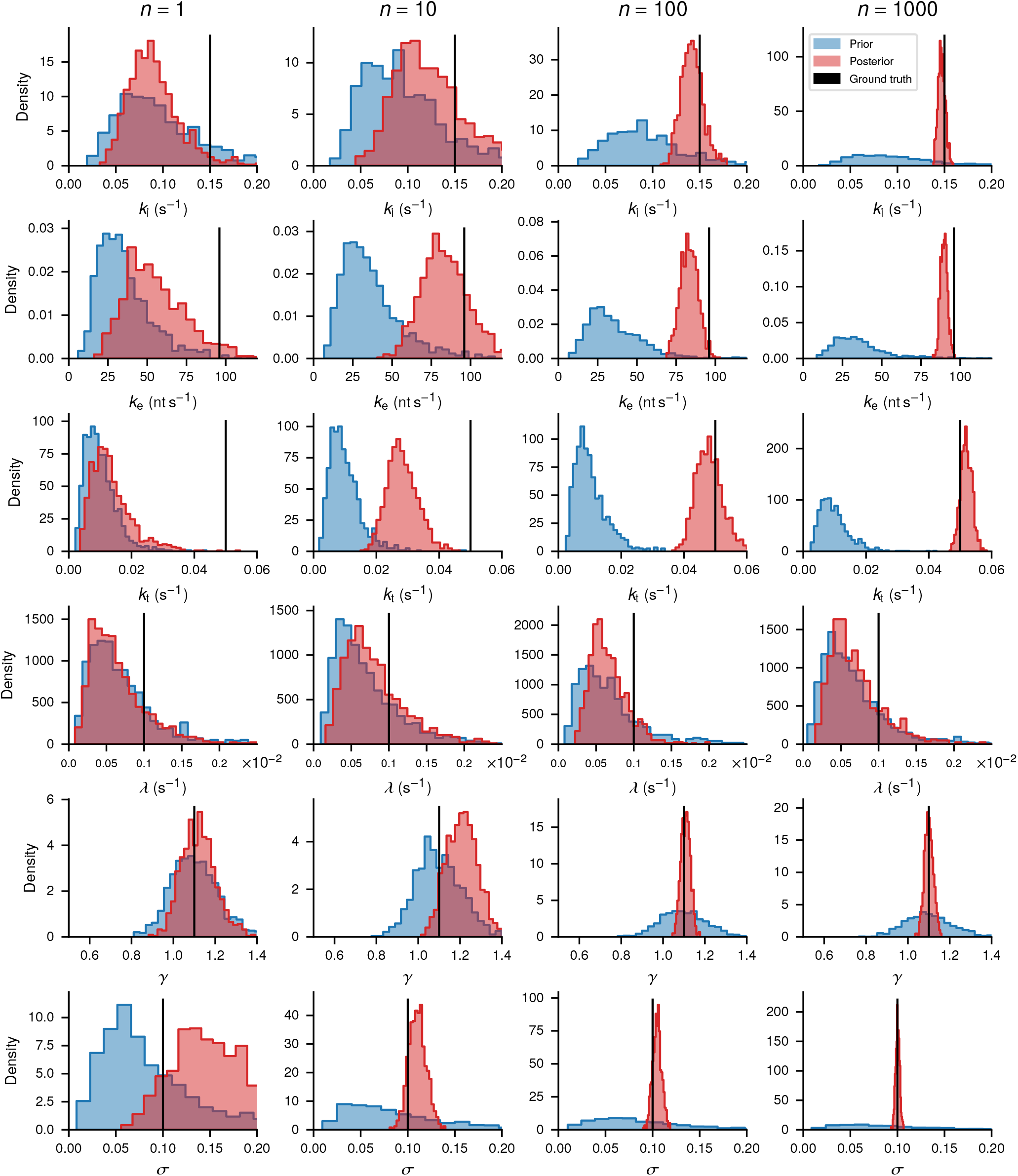
Joint posterior inference with different numbers of pooled traces (cf. Sec. S10.4) based on simulated data. The rows show histogram approximations of the prior distribution (blue) and the posterior distribution (red) for the model parameters. Black lines indicate the parameter value used to generate the data. The columns show realizations of the experiment with different numbers of trajectories. Some of the observation parameters are not shown, as results look similar.

**Figure S20.**
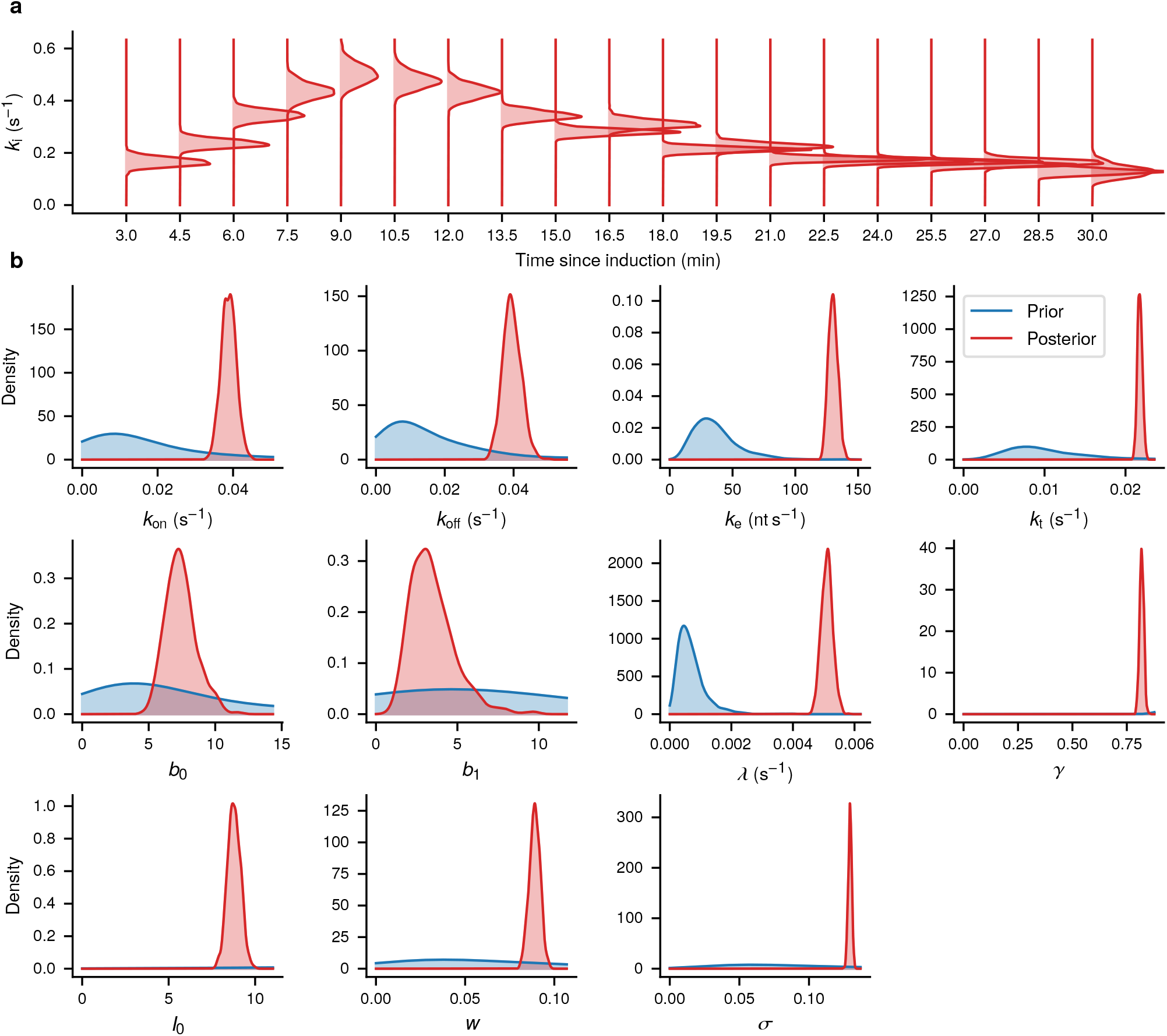
Posterior of the model with local initiation rate and remaining parameters global. This is an extended version of Fig. 4e,f of the main paper. **a** Local initiation rate per time window since induction. **b** Global kinetic and observation model parameters.

### S11 Model selection

In Bayesian inference there are two major approaches to model evaluation and comparison^35^. The first approach is based on the idea that we can perform posterior inference not only over parameters but also over models. Assuming there are a number of hypothesis *H*_1_, …, *H*_*m*_ corresponding to *m* models, some prior probabilities over the models *p*(*H*) and data 𝒟, we can compute *p* (*H* | 𝒟) to find the most probable model given the data. Typically, a uniform prior over the models is chosen which leads to *p* (*H* | 𝒟) *∝ p* (𝒟|*H*). In particular, if we are interested in comparing the odds of two models, we get

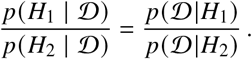

The fraction on the r.h.s. is called the Bayes factor, the term *p*(𝒟\*H*) is called the marginal likelihood or evidence. In a parametric model the marginal likelihood is given by

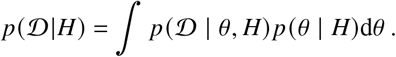

Therefore, the marginal likelihood can be used to score the performance of different models for a given dataset. The main advantage of the marginal likelihood as a model evaluation criterion is that it automatically penalizes model complexity. If two models explain data equally well, the one with fewer parameters will show a higher score. Unfortunately, the marginal likelihood is difficult and costly to evaluate, so typically approximations have to be used. A comprehensive review of such approximation techniques can be found in^**?**^. If inference is performed using a variational approach, an approximation to the marginal likelihood is obtained for free in form of the evidence lower bounds (ELBO). Classical variational inference is based on the idea to approximate the posterior by a distribution *q* in a tractable family 𝒬. The optimal approximation *q*^*^ is obtained by minimizing the Kullback-Leibler divergence

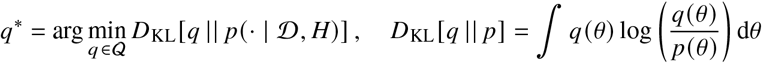

It can be shown that the above objective function decomposes as

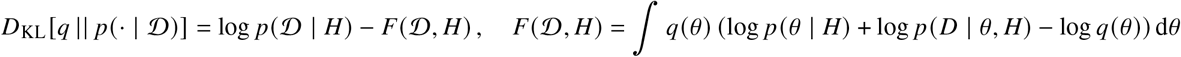

The quantity *F* (𝒟, *H*) is the ELBO and the maximization of the ELBO is equivalent to minimizing the KL divergence to the posterior. Furthermore, if the variational family *Q* is sufficiently expressive, *D*_KL_ [*q*^*^ || *p* (· | 𝒟)] is close to zero and we get

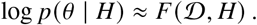

This immediately suggest an approximation of the Bayes factor as

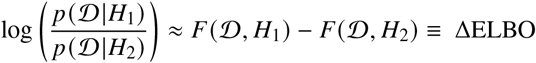

A conceptual drawback of evidence-based model selection is that it is only a relative measure of performance. If all models are bad and one model is slightly less bad, the latter one can still appear as a clear winner by evidence score. It is therefore advisable to combine evidence-based model selection with goodness of fit checks. In a Bayesian framework, the natural way to do this is by the posterior predictive distribution

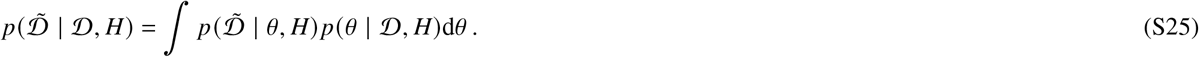

Intuitively, 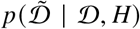 is the distribution of data simulated from the fitted model. Goodness of fit in this framework is assessed by comparing the similarity of certain summary statistics *T*(𝒟) of the true data with the same summary statistics of the posterior predictive distribution. More explicitly, for a discrepancy measure *D* (*T*(𝒟), 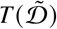 for two realization of the summary statistic, the predictive score is given by

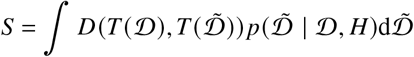

In practice, the integral with respect to the predictive distribution is performed by sampling from the parameter posterior *p* (*θ* | 𝒟, *H*) and simulating a dataset 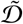 using the model 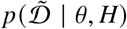. The choice of these summary statistics depends on the problem and should reflect features that are deemed important. In many applications, low order moments such as the mean or quantiles of the empirical distribution are chosen and the discrepancy measure *D* is, e.g. the *L*_2_-norm. Here, we are interested in the evolution of the intensity distribution over time. Consider a collection of traces 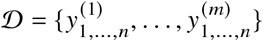. Let 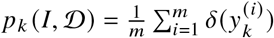 be the empirical intensity distribution at time *t*_*k*_. In this case, we are interested in the distribution of fluorescence intensity over time. As summary statistic we choose the collection of empirical distributions at the different time points, i.e. *T* = (*p*_*k*_)_*k*=1_^*n*^. The same collection of distributions is computed for a simulated dataset 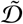. To compare individual distributions, we use the Wasserstein metric 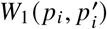. The Wasserstein metric is rooted in optimal transport theory. Intuitively, it describes the minimal amount of work required to transform one distribution into another one by moving infinitesimal pieces of mass at a time^50^. Averaging over individual time points leads to

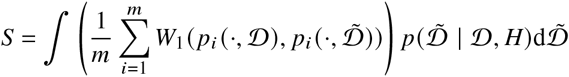

This statistic is computed separately for all windows and then averages over the windows. We do not provide the explicit formula is the notation becomes quite cumbersome. Finally, after computing *S* for all models of interest we can rank the models by how well the average predicted intensity distribution agrees with the empirical distribution.

Table S15 shows ΔELBO and the discussed posterior predictive score for various model configurations evaluated on both datasets. The relative ELBO is always provided for the simplest model, which can be considered as a baseline, for every dataset. A look at ΔELBO reveals that in both datasets there is a significant preference of the bursting models over the constitutive models. In addition, the time-dependent models score higher than the fully global model. Most of this gain is explained by a time-varying initiation rate. If other dynamic parameters are allows to vary, there is an additional but smaller gain in evidence. Interestingly, the fully local model shows a predictive performance close to other well fitting models but with lower evidence. This indicates that the additional degrees of freedom are not very useful for explaining the experimental data. In conclusion, S15 supports a bursting model with time-dependent parameters with most of the time dependence explained by the initiation rate.

**Table S15.**
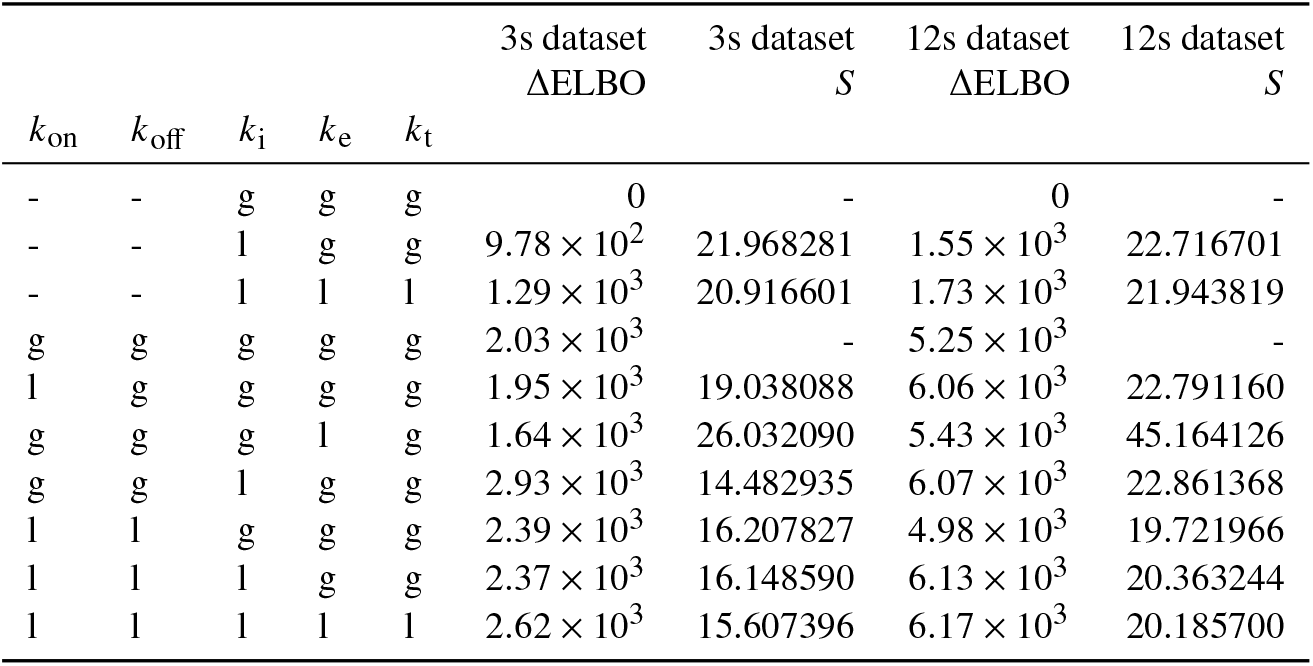
Bayesian model selection by approximate evidence and predictive scores based on Wasserstein metric. The index column on the right indicate which parameters are shared for all cells (g) and which are local for every time window (l). The dash indicated missing values, meaning that the corresponding model is not bursting. The quantity ΔELBO is shown with respect to the simplest model (shared parameter, no bursting) for every dataset. Therefore, higher values are better. *S* quantifies the average deviation of the intensity distribution between simulated and real dataset. Thus, lower values are better.

### S12 Tracking algorithm

Transcription site tracking in live cell fluorescence microscope imagery has to overcome three problems. First, fluorescent spots vary in intensity and may disappear altogether due to the underlying dynamics of the transcription process. Second, the background is non-homogenous and varies over time due to cellular clutter. Third, accumulations of fluorescent protein may cause spurious spots. Standard spot extraction algorithms typically run a detection step on each time frame and than combine possible spot candidates to a trajectory using a scoring scheme. We follow a joint approach combining detection and tracking within a stochastic filtering framework.

#### S12.1 Sequential Filtering Framework

The central idea is that the current position of the spot and the current intensity will provide information on the likely places to find the spot in the next frame. We denote the position of the spot in frame *k* as *r*_*k*_ = (*x*_*k*_, *y*_*k*_, *z*_*k*_)^⊤^. In addition, we define *I*_*k*_ and *b*_*k*_ as the spot and local background intensity, respectively. This leads to a full state **x**_*k*_ = (*r*_*k*_, *b*_*k*_, *I*_*k*_). We assume the evolution of these quantities between two consecutive images as given by

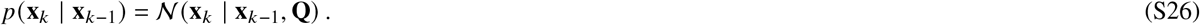

Eq. (S26) corresponds to a diffusive motion of the TS and and encourages the intensity variables at different times to be close. To deal with sudden vanishing and re-appearance of spots, we introduce a binary switching variable *s*_*k*_ representing the visibility. The observation 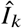 corresponds to the 3D image stack at step *k*. We connect the latent state **x**_*k*_ to the observed image 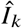 by a point spread function model as described in Sec. S9.3

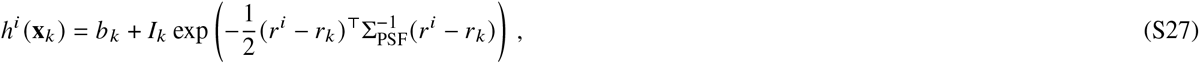

where *r*^*i*^ corresponds to the center of pixel *i*. In contrast to Sec. S9.3, we use a three-dimensional PSF model with 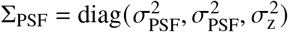. We also refrain from integrating over the pixel area and use the center of the pixel directly with the Gaussian profile. Together with a multiplicative noise as in (S14) we obtain

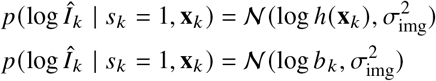

where the second equation indicates that only noisy background is observed when visibility is zero. This leads to a hidden Markov model with spot position and intensity as latent state and image stacks as observations (cf. Fig. S21). Computing 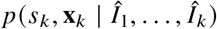, the distribution of the latent state given all observations up to step *k*, is known as the filtering problem and can be reduced to sequential prediction and update steps^43^.

**Figure S21.**
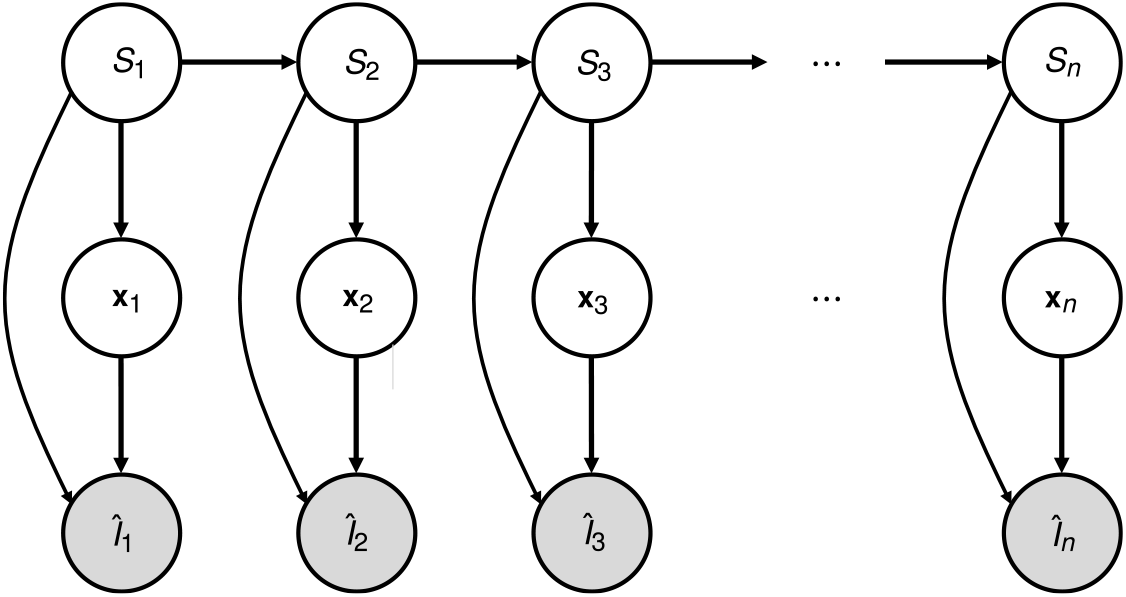
Probabilistic graphical model representation of the spot tracking problem with visibility variables *s*_*k*_, spot state **x**_*k*_ and noise observed image stack *I*_*k*_ .

#### S12.2 Prediction Step

Assume that at step *k* − 1 we have access to the filtering distribution

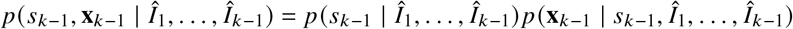

where 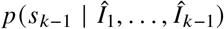 is a Bernoulli distribution and we assume that 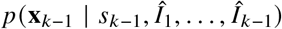 follows a Gaussian distribution. In this case, the prediction step can be computed explicitly as

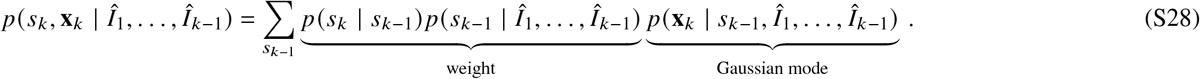

The last term on the r.h.s. can be computed analytically under the Gaussian assumption. Thus, (S28) becomes a Gaussian mixture with modes at the previous filter estimates for visible and invisible state and weights determined by the activity estimate.

#### S12.3 Approximate Update Step

Given the prediction step, the update step is given by

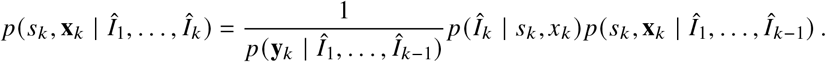

The normalizer is given by

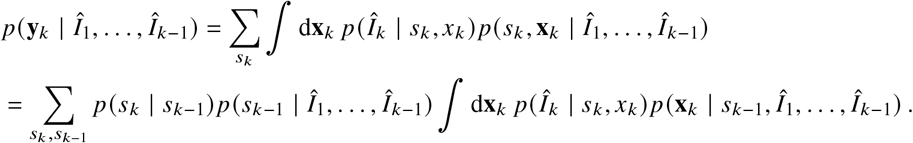

For the model discussed here, the update step cannot be solved in closed form due to the non-linear observation model (S27). In addition, the binary state *s*_*k*_ causes the filtering distribution to become a mixture with 2^*k*^ components. To obtain a tractable approximation, we combine two approximations. First, we observe that the non-linear observation likelihood is strongly peaked.

Thus it is reasonable to use a Laplace approximation

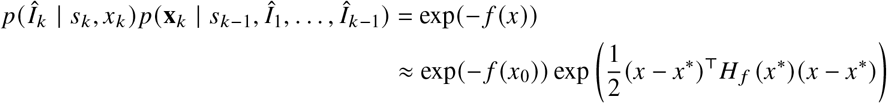

where

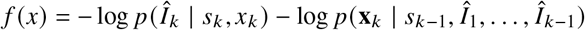

and *x*^*^ = arg min_*x*_ *f*(*x*). This leads to a representation of the filter in form of a Gaussian mixture distribution at every time. In order to keep the mixture from growing, we perform a mixture reduction via moment matching after every step.

## Notes

### Competing Interest Statement

The authors have declared no competing interest.

